# Benchmarking concentration and direct extraction methods for wastewater-based surveillance of eight human respiratory viruses: implications for rapid application to novel pathogens

**DOI:** 10.1101/2024.11.27.625007

**Authors:** Audrey Liwen Wang, Minxi Jiang, Allie Nguyen, Staci R. Kane, Monica K. Borucki, Rose S. Kantor, Kara L. Nelson

## Abstract

Wastewater monitoring is rapidly expanding to provide surveillance for a growing number of epidemic and endemic pathogens. To provide early warning and support rapid response to a novel virus through wastewater surveillance, it would be ideal to understand in advance which concentration and extraction methods are likely to be effective for dPCR-based methods, depending on virus characteristics. In this study, we spiked wastewater samples with eight human respiratory viruses and processed them with four commercial methods that concentrate and/or extract nucleic acids from both liquid and solid fractions (Promega, Nanotrap, InnovaPrep), or only the solid fraction of wastewater (Solids). Our findings provide encouraging evidence that all four concentration/extraction methods combined with dPCR could detect an emerging virus in wastewater, although they differed in sensitivity. The pattern of recovery efficiency for adenoviruses, coronaviruses, and influenza A viruses was consistent across methods, while distinct patterns were observed for coxsackieviruses. Promega produced higher median recovery efficiencies based on dPCR for all viruses except for coxsackieviruses, even though it had the highest dPCR inhibition. We suggest caution in applying Nanotrap to new targets, based on the low recovery of coxsackievirus B5 compared to the other viruses. We also quantified the endogenous indicators PMMoV and Carjivirus (formerly crAssphage), illustrating how normalization could either improve or worsen the comparison of virus concentrations measured by different methods. These findings can guide the selection of concentration and extraction methods for wastewater monitoring based on the properties of target viruses, thus enhancing pandemic preparedness.

**Synopsis Statement:** Benchmarking scalable concentration and extraction methods on respiratory viruses with diverse properties facilitates the rapid application of wastewater-based surveillance to emerging viruses.

## 1. Introduction

Since the COVID-19 pandemic, there has been a rapid expansion of wastewater-based surveillance (WBS) for monitoring population-level infectious disease trends in communities^1–4^. Many countries have launched wastewater monitoring programs for SARS-CoV-2 based on detection of nucleic acids by PCR, and have added additional pathogen targets to respond to public health priorities^5–7^. Human respiratory viruses are one of the most common causes of pandemics due to their fast transmissibility and high infectivity through respiratory droplets and aerosols^8,9^. To prepare for future pandemics, it would be ideal to have knowledge about which methods are likely to be effective for detecting novel targets in wastewater, to facilitate faster public health intervention. However, sensitive detection of viruses in wastewater remains a challenge, as different virus types, concentration and extraction methods, and wastewater characteristics can affect viral nucleic acid recoveries.

Interactions between the concentration/extraction methods and virus properties can lead to large variability in recoveries between virus types. For example, virus envelope types may influence the susceptibility to reagents as well as wastewater constituents such as detergents, solvents, and disinfectants^10,11^. Additionally, the characteristics of surface functional groups of viruses can affect their adsorption and partitioning to particulates in wastewater^12–14^. Adsorption in turn may affect method-specific recovery depending on the fraction(s) of a wastewater sample targeted by each method (liquid, solid, or both fractions)^15,16^. Finally, virus integrity (intact virus versus extraviral nucleic acids) and genome type (RNA versus DNA genome)^17^ could affect recovery, given that some methods extract total nucleic acids directly, while others first isolate viruses and then perform extraction.

Numerous method comparison studies highlight the challenges that result from differences in the nucleic acid recovery efficiency and dPCR sensitivity of each method^18–21^. Viruses are naturally concentrated in settled solids and high-throughput solids-based methods have been developed^22,23^. Meanwhile, for whole wastewater samples, viruses must be concentrated to achieve sufficient detection sensitivity. Prior to the COVID-19 pandemic, multiple concentration strategies were used including polyethylene glycol precipitation, ultracentrifugation, skim milk flocculation, and electronegative membrane concentration^20,24^. Several methods have since emerged that are compatible with high-throughput formats and require less processing volume to reach similar sensitivity^25–29^. The majority of method comparison studies for WBS have focused on coronaviruses and their surrogates^18–20,28,30,31^, while a small number compared the recovery of other virus targets with diverse properties^15,26,32,33^. To facilitate the rapid application of WBS to novel viruses, there is a need for additional method comparisons for viruses that span a larger range of physical and biological properties.

Here, we performed benchmarking of four commercial methods that are amenable to high-throughput formats: 1) Promega Wizard Enviro TNA Kit (“Promega”) for direct extraction of whole wastewater, 2) AllPrep PowerViral Kit for direct extraction of centrifuged solids (“Solids”), 3) Nanotrap magnetic beads, which is an affinity-based concentration method for whole wastewater (“Nanotrap”), and 4) InnovaPrep Concentrating Pipette Select for concentrating whole wastewater (“InnovaPrep”). We evaluated the recovery of eight respiratory viruses from four virus groups spanning a range of physical and biochemical properties: coronaviruses (SARS-CoV-2, OC43), influenza A viruses (H1N1, H3N2), adenoviruses (AdV-2, AdV-5), and coxsackieviruses (CV-A6, CV-B5).

The recovery of two endogenous fecal indicators (PMMoV and Carjivirus) were also compared. We further examined interactions between virus properties, method mechanisms, and varied wastewater characteristics to interpret the results. Based on the observed patterns of nucleic acid recovery across different methods and virus types, we provide recommendations for optimizing these techniques for the detection of other existing or emerging viruses that possess similar physical and biological properties.

## 2. Materials and Methods

Influent wastewater samples from three wastewater treatment plants (WWTPs) serving varying population sizes were collected at two time points (**Figure 1a**). Eight viruses were mixed and spiked into each 40-mL wastewater sample (**Figure 1b**). The forms of each virus stock (infectious, encapsidated but noninfectious, and extraviral nucleic acids only) were measured to provide insight into their recovery efficiencies (**Figure 1b**). The nucleic acid concentrations recovered by each of the four methods were quantified using digital PCR (dPCR) (**Figure 1c**).

**Figure 1.**
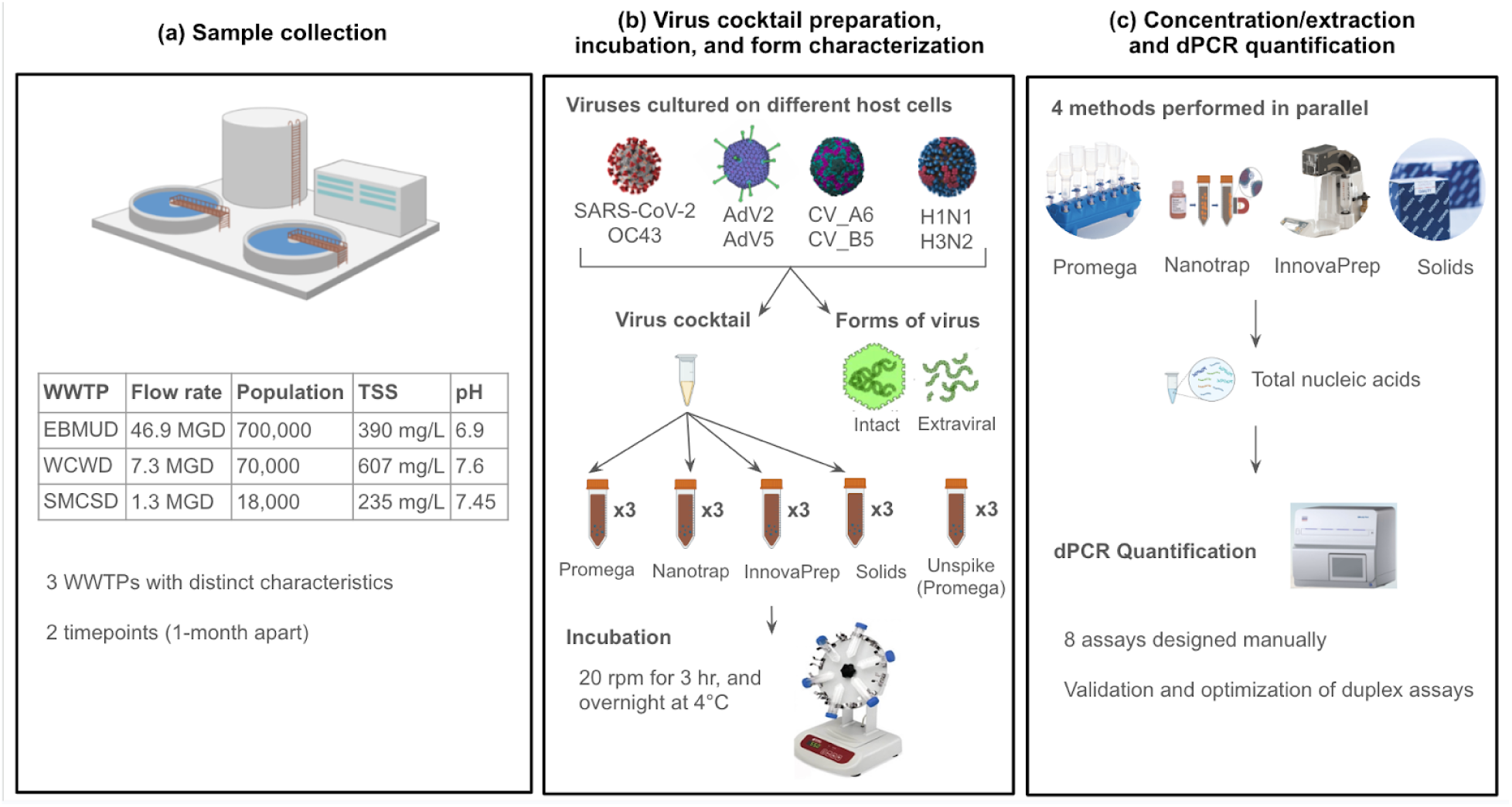
Overview of the experimental design.

### 2.1 Wastewater sample collection

24-hour composite samples of influent wastewater were collected from three WWTPs in the greater San Francisco Bay Area. The sampling locations were the East Bay Municipal Utility District (EBMUD), West County Wastewater District (WCWD), and Sausalito Marin City Sanitary District (SMCSD), which correspond to large (∼700,000) medium (∼70,000), and small (∼18,000) populations, respectively. For each facility, samples were collected on two dates, approximately one month apart: EBMUD samples on 7/26/2023 and 8/30/2023, WCWD samples on 7/31/2023 and 9/5/2023, and SMCSD samples on 8/2/2023 and 9/19/2023. This resulted in a total of six batches of samples. All samples were transported to the laboratory on ice and stored at 4 °C until further processing. Wastewater metadata including flow rate, biological oxygen demand (BOD_5_), total suspended solids (TSS), and pH were provided by wastewater agencies (**Table S1**).

### 2.2 Acquisition and culturing of viruses for spiking

Two strains or species from each virus group—coronavirus (CoV), influenza A virus (IAV), adenovirus (AdV), and coxsackievirus (CV)—were selected for this study. The selected viruses are common human respiratory viruses and span a range of physical and biological properties (**Table 1**). Each virus was propagated using the specified cell lines and culture conditions based on the previous protocols. Details about virus stocks and host cell information are provided in **Table S2**, and the propagation conditions and titer determination are described in the **Supplementary Methods**. For all viruses, the harvested cells were centrifuged at 2000 x *g* for 5 min to remove cell debris, and the supernatants were aliquoted into 1-mL tubes and stored at -80°C; no further purification steps were performed. Finally, heat-inactivated SARS-CoV-2 (isolate: USA-WA1/2020, NR-52286) was obtained from Biodefense and Emerging Infections (BEI) Resources.

**Table 1.** Physical and biological properties of the spiked-in viruses.

| Virus family | Strains/species | Genome type | Virus structure | Genome size (kb) | Virion size (nm) |
| --- | --- | --- | --- | --- | --- |
| Coronaviruses | SARS-CoV-2, OC43 | ssRNA | enveloped | 30 | 60-140 |
| Influenza A viruses | H1N1, H3N2 | ssRNA | enveloped | 13.5 | 80-120 |
| Enteroviruses | Coxsackievirus A6, B5 | ssRNA | non-enveloped | 7.6 | 22-30 |
| Adenoviruses | Type 2, 5 | dsDNA | non-enveloped | 35.5 | 90-100 |

### 2.3 Measurement of the forms of virus in the stocks and used for spiking

We determined the fractions of each virus stock that were: a) infectious, b) non-infectious but encapsidated, and c) extraviral. The infectious form was quantified by the median tissue culture infectious dose (TCID_50_/mL)^34^, while the other forms were determined based on dPCR with and without nuclease (DNase or RNase) pretreatment (**Figure S1A**). AdV virus stocks (DNA genome) were pretreated with DNase, and the rest of the virus stocks (RNA genome) were pretreated with RNase. The pretreatment process eliminated extraviral nucleic acids from the virus stock and ensured it was (>99%) encapsidated. All nuclease pretreatment experiments were performed in triplicate. Controls included triplicates of extracted viral nucleic acids treated with nuclease. DNase pretreatment followed a previously established protocol^17,35^, and RNase pretreatment was also conducted according to a standard method^17,36^ Full descriptions of both procedures can be found in the **SI Methods**.

Virus stocks were extracted using the solids protocol of AllPrep PowerViral kit (Qiagen) but with a modification of adding carrier RNA (Applied Biosystems™). Briefly, 200 µL of virus stock was added to PowerBead tubes, followed by the addition of PM1 and β-mercaptoethanol solution. The tubes were vortexed for 10 min, then centrifuged at 13,000 g for 1 min at room temperature. The supernatant was collected, and 6 µL of carrier RNA solution (1 µg/µL) was added to achieve a final concentration of approximately 0.01 µg/µL. The solution was incubated at room temperature for 5 minutes. Inhibitor Removal Solution (IRS) was then added, and the remaining steps of the protocol were followed as instructed. All virus concentrations were quantified with dPCR.

### 2.4 Virus cocktail preparation and wastewater spike-in

Using titer information from dPCR, a single virus cocktail was prepared as an equal-titer mixture of eight virus stocks with approximately 10^6^ gene copies of each virus. Aliquots of the virus cocktail (n=12) were stored at -80 °C until use. For each experiment, wastewater samples were homogenized and distributed into 15 replicates of 40 mL each. One virus cocktail aliquot was thawed to add to 12 wastewater aliquots (for four methods in triplicates), and 200 µL of the virus cocktail was then extracted using the modified solids protocol of AllPrep PowerViral kit (Qiagen), following the exact procedure described for virus stocks extraction outlined in **Method 2.3**.

Three unspiked wastewater aliquots were used to determine endogenous virus concentrations via the Promega method (procedures described below in **Method 2.5**). All 15 wastewater aliquots were rotated at 20 rpm for 3 h at room temperature using Multi-Purpose Tube Rotators (Fisher Scientific) to reach equilibrium partitioning of viruses to solids^37^, and then stored overnight at 4 °C. The next day, samples were rotated again at 20 rpm for 20 min at ambient temperature to re-equilibrate prior to concentration and extraction.

### 2.5 Concentration and direct extraction methods

Four concentration and extraction methods that employed different mechanisms were performed in parallel (**Figure S2**). To standardize the four protocols, the AllPrep PowerViral DNA/RNA Kit (Qiagen) was used for all extraction steps. The starting material was 40 mL of whole wastewater for all methods, and each method resulted in 100 μL of purified nucleic acids. The purified nucleic acids were aliquoted and stored in -20 °C and went through one freeze-thaw before dPCR testing.

**Promega Wizard Enviro TNA Kit for direct extraction of whole wastewater (“Promega”)** was performed according to the manufacturer’s instructions. Briefly, 0.5 mL of protease solution was added to the 40 mL wastewater sample and was mixed by inversion. The sample was incubated for 30 min at ambient temperature, and then large particles were removed via centrifugation at 3,000 x *g* for 10 min. Binding buffers and isopropanol were added to the resulting supernatant, and the mixture was passed through a PureYield binding column. Following two wash steps, the total nucleic acids were eluted in 1 mL of nuclease-free water and further purified on a PureYield Minicolumn to produce a final volume of 100 µL purified nucleic acids.

**AllPrep PowerViral DNA/RNA Kit for direct extraction of centrifuged solids (“Solids”)** was performed according to the manufacturer’s solids extraction protocol, which started with centrifuging a 40-mL wastewater sample at 20,000 x *g* for 10 min. The total weights of the centrifuged solids (ranging from 0.22 to 0.82 g) were recorded **(Table S3)**. Then, 0.22-0.25 g (wet weight) of the solids pellet was loaded in the Powerbead tubes with the addition of PM1 and Beta-mercaptoethanol solution for 10-min vortexing. The resulting supernatant was loaded onto the MB Spin Column following two washes and final elution, yielding 100 µL purified nucleic acids.

**Nanotrap magnetic beads for whole wastewater concentration (“Nanotrap”)** was performed according to the Nanotrap Microbiome A Protocol (APP-091 Ceres Nanosciences SKU), which was compatible with the downstream liquid extraction protocol of Qiagen AllPrep PowerViral DNA/RNA Kit. The starting wastewater volume increased from 35 mL (original protocol) to 40 mL, and all other reagents were correspondingly scaled. Briefly, 600 µL of Nanotrap Microbiome A Particles and 115 µL Nanotrap Enhancement Reagent 2 were added to the whole wastewater sample and mixed by inversion. The sample was incubated for 30 min at ambient temperature. Then, a magnetic rack was applied to separate the Nanotrap particles from the solution. The supernatant was discarded and the Nanotrap particles were resuspended in 1 mL of nuclease-free water. A magnetic rack was further applied to separate out the Nanotrap particles and the supernatant was discarded. Particles were resuspended in 600 µL of PM1 and Beta-mercaptoethanol solution, followed by 95 °C heating for 10 min to lyse the cells. Subsequent nucleic acid extraction steps followed the protocol.

**InnovaPrep Concentrating Pipette Select™ for liquid concentration (“InnovaPrep”)** began by adding 5% of Tween 20 (Thermo Scientific) to the 40 mL wastewater sample and mixed by inversion to release virus particles adsorbed to solids^38^. The sample was then centrifuged at 7000 x *g* for 10 min, and the supernatant was collected and passed through a Ultrafilter Concentrating Pipette Tip (CC08004 Unirradiated, InnovaPrep) to concentrate the viruses.The virus concentrate was eluted by Elution Fluid - Tris (InnovaPrep) into viral concentrate (ranging from to 170 to 720 µL, **Table S3**). Then, 170-200 µL of the viral concentrate was extracted according to the Qiagen AllPrep PowerViral liquid extraction protocol, yielding 100 µL purified nucleic acids.

### 2.6 Quantification of extracts by dPCR and inhibition testing

All virus targets were quantified using the QIAcuity Four Platform Digital PCR System (Qiagen). The virus assays were designed using PriMux^39^ and Primer3Plus^40^ (**Table S4).** All materials and conditions were summarized in **Table S5**. The reaction mixtures for RNA viruses (**Table S5**) were prepared using the QIAcuity OneStep Advanced Probe Kit (Qiagen), while QIAcuity Probe Master Mix (Qiagen) was used for DNA viruses. The virus assays were duplexed for the same virus group. The reaction mixtures were loaded to either 8.5k plates for quantifying spiked viruses or 26k plates for quantifying endogenous viruses. Priming and imaging used default settings and thermal cycling conditions are reported in **Table S5**. Partition volume is 0.2 nL. Valid partition counts ranged from 7,165 to 8,293 per well for 8.5k plates and 11,638 to 25,492 per well for 26k plates. The operational limit of detection was treated as ≥3 positive partitions per well. The positive control was linearized plasmid DNA (SARS-CoV-2) or gBlock standards from Integrated DNA Technologies, and the negative control was nuclease-free water. All positive controls showed clear separation between positive and negative partitions, while negative controls showed zero positive partition **(Figure S3)**. Data were analyzed using the QIAcuity Suite Software V1.1.3 (Qiagen) with automated settings for threshold and baseline, followed by manual inspection. To test for inhibition, each extracted sample was run at 1:5 dilution with the aim to reduce the concentration of potential inhibitors. The Environmental Microbiology Minimum Information Checklist^41^ is provided in **Table S6**.

### 2.8 Data analysis

All data analysis was performed in Python (v3.10.12) using package scipy.stats (v1.11.4) and scikit-posthocs (v.0.9.0), and statistical significance was determined at a 95% confidence interval (p < 0.05). Kruskal-Wallis H-test was employed to compare differences among concentration and extraction methods within the same virus type. Dunn’s test with Bonferroni correction was used as a post-hoc test following a significant result in a Kruskal-Wallis test. Virus concentration in purified total nucleic acids (C_purified_TNA_, gc/µL-TNA) was determined according to **Equation 1**, and the recovered concentration from 40 mL of wastewater (C_recovered_, gc/mL-WW) was calculated based on **Equation 2**.

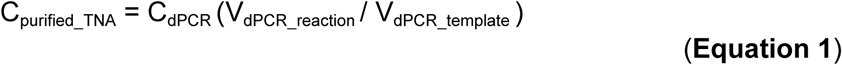

Where C_dPCR_ is the concentration reported by dPCR (gc/µL), V_dPCR_reaction_ is the final reaction volume (µL) for dPCR, V_dPCR_template_ is the template volume (µL). The volumes were reported in **Table S5**.

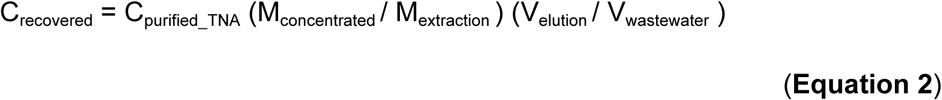

Where M_concentrated_ refers to the total mass (g) of solid pellets or concentrated liquid (µL) obtained from the Solids method centrifugation or the InnovaPrep concentration step, respectively. The mass was recorded in **Table S3 (column I)**. M_extraction_ represents the actual mass of the solid or liquid sample used for the subsequent extraction step, which is 0.22-0.25 g for the Solids method and 170-200 µL for the InnovaPrep method. V_elution_ is 100 µL, which is the final elution volume from the extraction kit. V_wastewater_ is 40 mL, representing the volume (mL) of one raw wastewater sample.

Recovery efficiency (%) of each virus was calculated as follows:

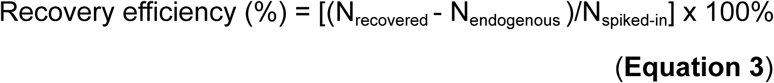

Where N_recovered_ is the number of spiked gene copies recovered by each concentration/extraction method from a 40-mL wastewater sample. N_endogenous_ is the number of endogenous gene copies present in the background of a 40-mL wastewater sample, which was measured using the Promega method. Finally, N_spiked-in_ is the initial number of gene copies spiked into a 40-mL wastewater sample, which was measured by extracting viruses from the virus cocktail.

## 3. Results

Eight human viruses (**Table 1**) were spiked into raw wastewater samples from three treatment facilities at two time points (**Figure 1**). Samples were homogenized and incubated to reach equilibrium, and then processed in parallel by four virus concentration and/or extraction methods.

### 3.1 Recovery efficiency varied by method and by virus

Recovery efficiencies were calculated for each method and each virus (**Equation 3**; **Figure 2A**) from the dPCR measurements. The input virus concentrations varied somewhat across the six experiments and the eight viruses (**Figure S4A**), and were accounted for in the calculation of recovery efficiency. The concentrations of endogenous wastewater viruses were confirmed to be >100-fold lower than the spike-in level (**Figure S4B**). Several measurements of recovered spiked virus quantities were below the dPCR limit of detection (**Figure S4C**). For fecal indicator viruses, instead of recovery efficiency, the amount of each virus recovered was directly compared because the starting concentrations were not known (**Figure 2B**).

**Figure 2:**
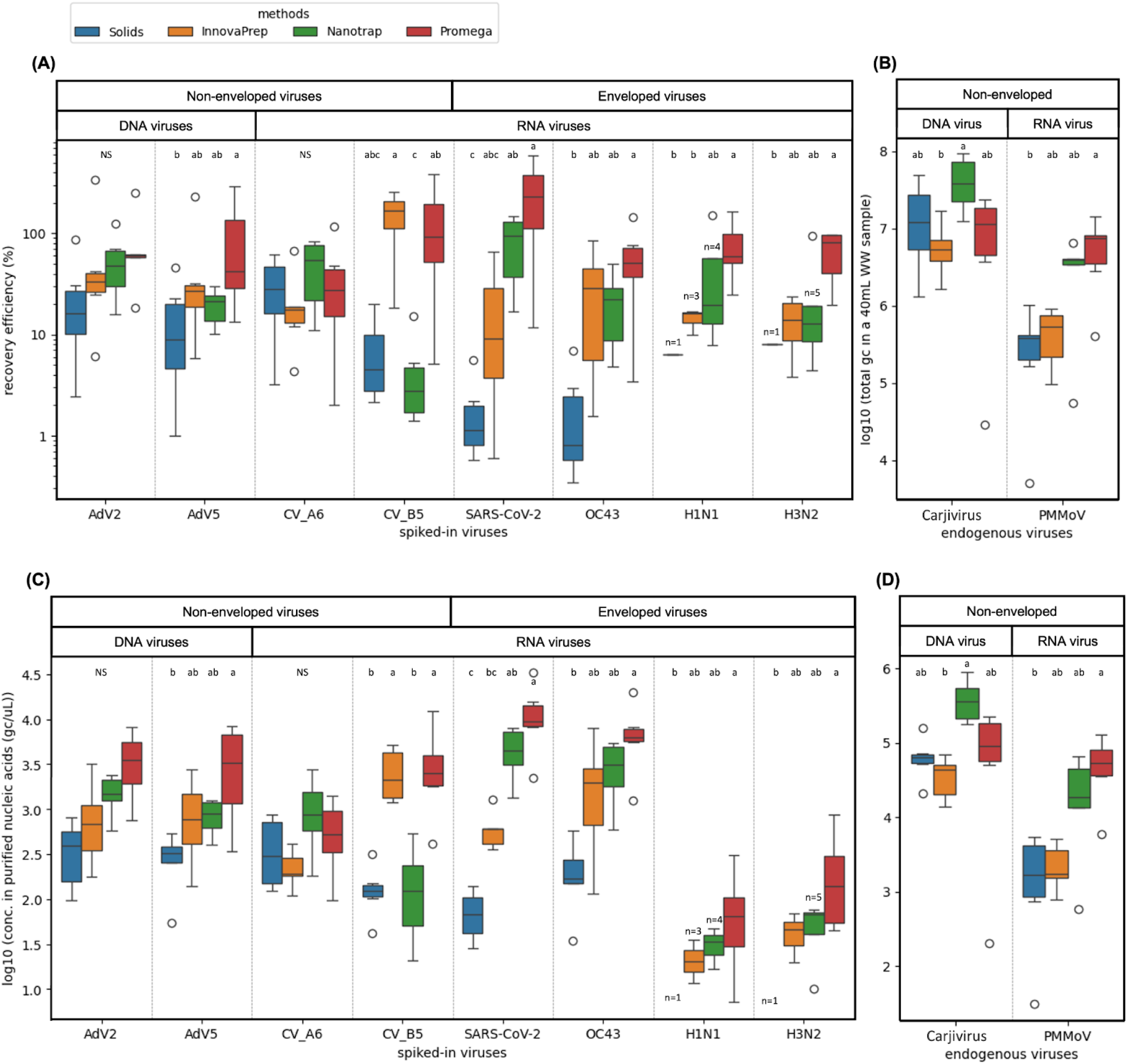
Recovery efficiency (%) of the **(A)** spiked-in viruses and **(B)** endogenous fecal indicators across four methods. Concentration of the **(C)** spiked-in viruses and **(D)** endogenous fecal indicators in purified nucleic acids (gc/µL) across four methods, which implies the limit of detection (LOD) varied for each method and virus. For all figures, boxes and whiskers indicate the interquartile range (IQR) and minimum / maximum values within 1.5 times the IQR, respectively, across all samples (n=6 for two time points and three wastewater sources; three biological replicates from each wastewater were first averaged by the geometric mean). The total number of samples (n) is shown when less than 6, due to concentrations below the dPCR detection limit. Outliers are displayed as individual points. Significance letters are shown based on the results of the post-hoc Dunn’s test for pairwise comparisons of methods within each virus type. Any methods sharing the same letter are not significantly different from each other. NS = no significant differences.

Mean recovery efficiencies ranged from approximately 0.34% (OC43 extracted by Solids) to around 400% (SARS-CoV-2 extracted by Promega) (**Figure 2A**). The recovery efficiencies that exceed 100% highlight one of the challenges in accurately quantifying virus recovery, which is further discussed in **Discussion 4.2**. Among the enveloped viruses, strains from the same virus group displayed a similar recovery pattern across the four methods, and Promega performed significantly better than Solids (p_dunn_ < 0.05, **Table S7**). Among the non-enveloped viruses, the two adenoviruses showed similar recovery trends across different methods, but the two coxsackieviruses differed. Recovery of CV-A6 was not significantly different by any method, while recovery of CV-B5 was lowest with Nanotrap and highest with Promega and InnovaPrep. Lastly, the pattern of recovery for endogenous PMMoV across methods was similar to that of adenoviruses, coronaviruses, and influenza A viruses (**Figure 2B**; Promega > Solids, p_dunn_ = 0.013, **Table S7**). Meanwhile for endogenous Carjivirus (formerly crAssphage), all methods performed similarly, though InnovaPrep appeared slightly less efficient (statistically significant only between Innovaprep and Nanotrap).

### 3.2 Sensitivity and inhibition varied across methods

To understand differences in method sensitivity from a practical perspective, concentrations of each virus genome in the purified nucleic acids (the input material for dPCR) were calculated for all methods (**Equation 1**; **Figures 2C and 2D**). The Solids and InnovaPrep methods were characterized by low sensitivity relative to other methods for most targets (**Figure 2B**); the exception was CV-B5, which was well-recovered by InnovaPrep. Limitations of the extraction kit prevented extraction of the entire sample for these methods. The average effective volume processed was 27 mL for InnovaPrep and 30 mL for Solids, compared to 40 mL for Promega and Nanotrap (**Table S3**). Additionally, we observed variable levels of inhibition across different wastewater sources, dependent on the dPCR assay (**Figure S5**). Where inhibition was observed, it was highest with the Promega method.

### 3.3 Forms of viruses in the input differed

Given that the form of a virus in wastewater could affect method performance here and for future viruses of concern, we determined the fractions of each viral stock that were: a) infectious, b) non-infectious but intact (protected from digestion by nucleases), and c) extraviral (**Figure 3**). The infectious form was quantified by TCID_50_, while the other forms were determined based on dPCR with and without nuclease pre-treatment (**Figure S2**). Both coxsackieviruses were primarily intact but non-infectious (CV-A6 = 75.6±1.45%; CV-B5 = 77.8±1.1%). Adenovirus Type-2 and Type-5 contained 58.5±1.51% and 45.8±1.24% extraviral material as a percentage of total viral genome copies, respectively. Interestingly, all of the nuclease-protected AdV5 was infectious (54.3±1.14%), whereas only around 5% out of the total AdV2 was infectious. Influenza A viruses were primarily extraviral (both strains around 75%), while nearly all of the OC43 stock consisted of nuclease-protected but non-infectious forms (87.7±1.13%). Lastly, the heat-inactivated SARS-CoV-2 was primarily extraviral (83.8±1.4%), while the TCID_50_ was certified to be zero by the vendor. We note that virus forms may have changed during incubation in the wastewater, but we did not attempt to account for potential changes.

**Figure 3:**
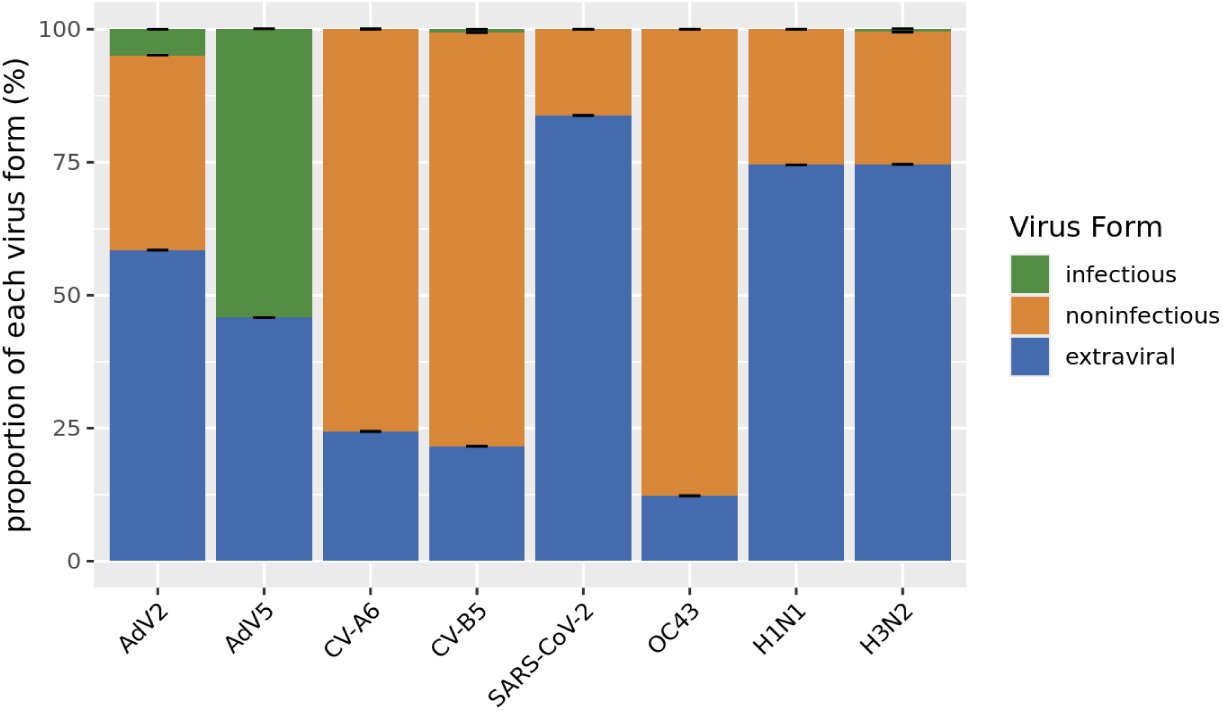
Virus stocks were characterized to determine intact infectious (green), intact non-infectious (orange), and extraviral (blue) fractions. To determine these fractions, three measurements were taken: TCID_50_ (intact infectious), intact nucleic acids after DNAse/RNase treatment (= encapsidated form = intact infectious plus intact non-infectious), and total nucleic acids (= all three forms). The intact non-infectious form was the difference between the encapsidated form and TCID_50_. The extraviral form was obtained from subtracting total nucleic acids with encapsidated form. TCID_50_ was not determined for SARS-CoV-2, which was purchased as an inactivated stock. The standard deviation for each proportion was calculated using error propagation.

### 3.4 Solids partitioning estimates differed by method

For comparability with other methods comparison studies, we calculated virus concentrations on a per-mass basis (gc/gTSS for the Solids method and gc/mL wastewater for the other methods). When compared on a per mass basis, all targets were higher in Solids samples than whole wastewater samples extracted by InnovaPrep, Promega, and Nanotrap (**Figure S6**). To determine which viruses were most enriched in the solids fraction of wastewater on a mass equivalent basis, we calculated ratios of mass-equivalent gene concentration of Solids method (gc/gTSS) to the whole wastewater methods (gc/mL WW) (**Table 2**). Note that these ratios are not distribution coefficients^37^, because all three of the whole wastewater methods were designed to capture viruses from both the liquid and the solids fractions of the sample. As expected, the ratios depended on the whole wastewater methods, as has been shown previously^23^, with the lowest ratios found for Promega. Ratios also varied up to 100-fold between viruses, e.g., for Solids:InnovaPrep the ratio was 4550 mL/g for CV-A6 and 104 mL/g for Influenza A H1N1. Surprisingly, most ratios were lower for enveloped viruses, which were expected to have higher solids association than the non-enveloped viruses. We note that there is low confidence in the IAV concentrations measured by the Solids method because many samples were below the detection limit (**Figure 2C**).

**Table 2:**
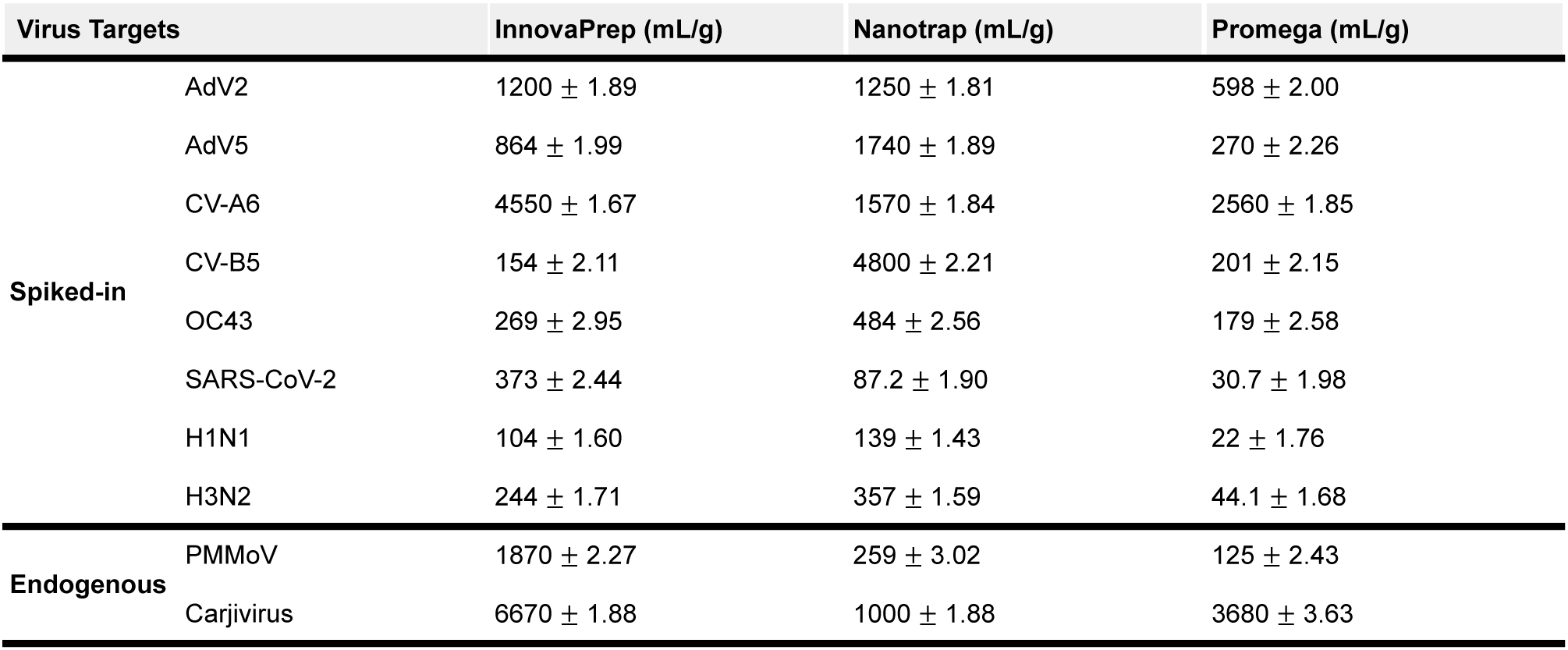
Mass-equivalent gene concentration ratios (mL/g) of Solids to InnovaPrep, Nanotrap, and Promega for spike-in viruses and endogenous fecal indicators. Each value shown is the geometric mean of six samples (n=6 for two time points and three wastewater sources; three biological replicates from each wastewater were first averaged by the geometric mean as well), except for several IAV samples that were below dPCR detection limit. The geometric standard deviation was calculated using error propagation. The ratios were calculated on a mass-equivalent basis and the values were rounded to three significant figures.

## 4. Discussion

### 4.1 Interactions between virus characteristics and the underlying mechanism of each method may affect recovery

To understand the observed recovery efficiencies, we explore how the capsid properties and forms of the different viruses may have interacted with the specific mechanisms employed in the four methods.

#### 4.1.1 Promega

The high recoveries by Promega reported previously for SARS-CoV-2 were also observed for all viruses studied here^25,29^. Three factors may contribute. First, the direct extraction approach eliminates the need for a prior concentration step, which may reduce losses compared to multiple-step concentration-extraction approaches^21^. Second, Promega is designed to recover viruses from both the liquid and the solid fractions of a 40-mL wastewater sample. While the specific mechanism for recovery of target from the solids is not reported by the manufacturer, it is possible that the protease treatment prior to solids removal causes lysis of viral protein capsids and release of nucleic acids, as has been previously demonstrated with other proteases^42,43^. Third, the method may capture extraviral nucleic acids present in the original sample, via binding to the silica column. Thus, the Promega method appears less biased toward different viral forms and virus partitioning to solids. Notably, Promega also had the highest levels of dPCR inhibition for RT-dPCR assays (**Figure S5**), which may also explain why the recoveries of coxsackieviruses and OC43 were not higher. Additional inhibitor removal steps may improve the results for this method.

#### 4.1.2 Nanotrap

This method generally had the second highest recovery, after Promega. Nanotrap particles are magnetic hydrogel particles coupled with high affinity baits that bind to a broad range of proteins^44,45^. The composition of the baits determines the targets^46–48^. For instance, Nanotrap particles containing the affinity dye cibacron blue were most effective at capturing rift valley fever virus (RVFV) and human immunodeficiency virus type 1 (HIV-1)^46^, while particles with red baits were more effective for concentrating influenza A viruses^48^. The low recovery of CV-B5 by Nanotrap in our study was an exception to the generally good recovery for the other viruses; it is possible that the mixture of baits in the Microbiome A reagent had a lower affinity for this virus due to its capsid structure. Given that the recovery efficiency of CV-A6 and CV-B5 differed by around 20-fold, caution should be exercised in extrapolating the performance of Nanotrap to new viruses, even those that are closely related. While we did not remove solids via centrifugation prior to using the Nanotrap beads, other laboratories may, and our results may not reflect the recoveries achieved with a solids removal step.

#### 4.1.3 InnovaPrep

InnovaPrep concentrates viruses via size-exclusion^49^, and we selected the Concentrating Pipette Tips characterized as ultrafiltration (specific pore size not reported for product CC08004). Despite the possibility that the pores might not capture extraviral nucleic acids as effectively as intact viruses, we did not observe a trend between lower recovery efficiency and percentage of extraviral nucleic acids; a more controlled study would be needed to isolate this factor. InnovaPrep yielded higher recovery efficiency for CV-B5 than for other viruses, including CV-A6 (**Figure 2A**). Possible reasons for the higher recovery efficiency of CV-B5 relative to other viruses include that it had lower adsorption to wastewater particles, was more effectively released from wastewater particles by the addition of Tween 20 prior to solids removal, or was more stable in the wastewater and in the presence of Tween 20. Our findings are consistent with a recent study that found CV-B5 was not effectively removed by activated sludge treatment via sorption to biological flocs or degradation, compared to several other viruses that had higher association with biological flocs^14^. More generally, enteroviruses have been shown to have widely different susceptibility to chemicals (such as chlorine^50^) due to differences in their capsid structures, reinforcing that different behavior can be observed among closely related viruses.

#### 4.1.4 Solids

The low recovery efficiencies reported by the Solids method (**Figure 2A**) reflect the fact that the mass of solids in a 40 mL sample is relatively small compared to the mass of liquid. The result aligns with a recent study^51^ which found that assaying the liquid fraction of the influent wastewater led to more sensitive virus detection due to the low solids content of the influent wastewater, despite viruses being more concentrated in the solid fraction. Of all the targets in this study, CV-A6 and Carjivirus were recovered the most efficiently by the Solids method (relative to the whole wastewater methods), which could be due to higher partitioning to solids, and/or lower recovery by the whole wastewater methods. Interestingly, in a companion paper on wastewater sequencing, we found that Promega and InnovaPrep resulted in a greater reduction in bacteriophage sequences compared to Solids and Nanotrap, which we attributed to the solids removal steps^52^. Perhaps a large fraction of the bacteriophage, including Carjivirus, were present inside or attached to the host bacteria, and were not effectively released from the solids by the protease (Promega) or surfactant (InnovaPrep) steps prior to solids removal.

### 4.2 Challenges in conducting and interpreting methods comparison studies

Some of the recovery efficiencies measured in this study were greater than 100%, which highlights one challenge with spike-in studies - it is not possible to quantify the “true” concentration of target in the spike-in solution. We used the Qiagen AllPrep PowerViral kit to quantify the virus spike-in solution (prior to addition to wastewater), whereas recovery of the spiked virus targets after mixing with wastewater was performed with the Promega kit, and the recovery appears to have been higher. Recoveries higher than 100% have been reported previously and attributed to the same phenomenon^19,21^. Our results also illustrate that it is not possible to apply the recovery efficiency measured for a spike-in proxy virus (e.g., bovine coronavirus) to accurately account for the recovery of a target virus, because even for the same method, the recovery efficiency varies depending on the target^21^.

Spike-in studies are often used to evaluate the recovery of targets that are not consistently present as endogenous viruses in sufficient concentrations in wastewater. However, lab-cultured viruses added to wastewater may not perfectly mimic the behavior of endogenous viruses, even though we followed methods that have been used by others to allow partitioning of the added viruses to solids^37^. The fact that we observed patterns of recovery for the two endogenous indicators (Carjivirus and PMMoV) that matched those of the spiked-in viruses lends confidence to our results. Further, variations exist in the procedures of each of the methods we used, which could have an impact on the recovery efficiencies. For example, we did not employ RNA shield or grinding balls in our Solids method^23^, which could have led to lower recovery efficiencies. As noted in **Discussion 4.1.2**, some laboratories remove solids via centrifugation before using Nanotrap beads^53^, which would be expected to reduce the recovery efficiency. The method used for extracting nucleic acids could influence both the recovery efficiency and the inhibition; based on our results, the PowerViral kit produced extracts without significant PCR inhibition.

### 4.3 Implications for practice

#### 4.3.1 Any of the methods has potential to detect emerging pandemic viruses

Our findings provide encouraging evidence that all four of the methods tested would be able to detect an emerging virus in wastewater. Currently, most surveillance programs are based on observing changes in concentration relative to a baseline value (e.g., CDC NWSS wastewater viral activity level^74^), which avoids the need to account for differences in recovery efficiency between methods, labs, and targets. To enable early detection it would be ideal to use methods with high sensitivity. While we found that Promega was generally the most sensitive of the methods we tested, it was also the most prone to PCR inhibition. The sensitivity of the other methods tested here can be increased by processing larger sample volumes and by pooling results from more than one replicate. Virus stability in wastewater may also affect the suitability of a given method for early detection^13,14^: the Promega method appears to recover extraviral nucleic acids, which could be important if an emerging virus lyses during its transport through the sewer. Broadening the scope beyond viruses, each of the tested methods has the potential to be applied to other classes of pathogens (e.g., bacteria, protozoan cysts, fungi); the Promega, Innovaprep, and Solids methods are not specific to viruses, and the Nanotrap method can be modified by incorporating Nanotrap B particles. Note also that while our findings should generalize to other types of quantitative PCR, optimal methods to prepare samples for sequencing endpoints have been shown to be different^35^.

#### 4.3.2 Use of endogenous indicators to improve continuous monitoring

Endogenous fecal indicators are commonly used to normalize the fecal strength of the wastewater (e.g., accounting for dilution by infiltrating groundwater or stormwater). In addition, endogenous indicators can potentially be used to account for differences in recovery efficiency between methods so that wastewater concentrations can be compared across datasets generated with different methods. A novel finding from this work is that the pattern of recovery observed for PMMoV reflected that of the adenoviruses, coronaviruses, and influenza A viruses.

Thus, normalizing the concentrations of these targets (by dividing by PMMoV) measured by any of the four methods tested here may make the values more comparable to each other across methods. Similarly, based on our results, Carjivirus appears to be an appropriate indicator to normalize CV-A6 concentrations for differences in recovery efficiency. In contrast, normalizing concentrations of CV-B5 measured by Nanotrap and Innovoprep by dividing by the respective concentrations of Carjivirus would actually make the concentrations measured by these different methods less comparable.

### 4.4 Future research directions

For the application of WBS, little is known about the forms of virus shed by infected individuals^54,55^, which could impact the persistence of the viral nucleic acid signals in the sewer system^17^ as well as during sample collection and processing. We recommend further research to understand the contribution of extraviral nucleic acids to fecal shedding values. Most studies quantifying recovery efficiencies by concentration and extraction methods do not report the forms of virus in the samples used, which could vary dramatically for spike-in studies depending on the virus culture and purification methods. We recommend that this information be included in future studies. While we found that the proportions of the dPCR signal due to extraviral nucleic acids varied significantly across the targets we studied, we could not isolate the influence these differences might have had on the observed recovery efficiencies of the four methods. A more controlled study, in which the proportions of intact virus and extraviral nucleic acids vary over a wide range while keeping other factors constant, would be needed to understand the impacts for each method.

## Acknowledgements

Funding was provided by the UCOP Lab Fees CRT Award (L22CR4507). We thank Khi Lai at EBMUD, Randy Hart at SMCSD, and Geraldine Gonzales at WCWD for their assistance with wastewater sample collection, Van Trinh and Joaquin Jamieson for help with dPCR and TNA extraction, and Jordan Boeck for help with virus culturing. Vero/TMPRSS2 cells were generously provided by Sean Whelan at the Washington University in St. Louis to Dr. Borucki. The SARS-CoV-2 isolate USA-WA1/2020 (heat-inactivated, NR-52286) was deposited by the Centers for Disease Control and Prevention and obtained through BEI Resources, NIAID, NIH.

## Notes

### Competing Interest Statement

The authors have declared no competing interest.

### Summary of Updates

Add myself (Audrey LiWen Wang) as a first-author to the "Author list".

