## Supplementary_Materials for "Benchmarking concentration and direct extraction methods for wastewater-based surveillance of eight human respiratory viruses: implications for rapid application to novel pathogens"

##### Supplementary Tables:

**Table S1.** Wastewater characteristics (metadata) on each sampling day

**Table S2.** Virus stocks acquisition and host cells information

**Table S3.** Sample metadata, effective volume, and dPCR concentration (see excel file)

**Table S4.** dPCR forward primer, reverse primer, probe, and dye of every virus target

**Table S5.** dPCR cycling conditions and reaction mix recipe

**Table S6.** dMIQE checklist for RT-dPCR experiments

**Table S7.** Statistical analysis of **(A-B)** recovery efficiency, **(C-D)** purified total nucleic acid concentration, and **(E-F)** total gene copies in a 40mL wastewater sample

##### Supplementary Figures:

**Figure S1. (A)** Graphical presentation of the calculation of different forms of viruses. **(B)** Concentrations of infectious virus from $TCID_{50}$/mL, intact virus concentration, and total virus concentration

**Figure S2**. Brief overview of methods workflow.

**Figure S3.** Examples of partition fluorescence plots of dPCR positive and negative control

**Figure S4. (A)** Initial virus concentrations in 40 mL of wastewater based on the concentration of the pure virus cocktail; **(B)** Endogenous virus concentration in 40 mL of wastewater quantified from the unspiked samples; **(C)** Recovery efficiency (%) of the spiked-in viruses across four methods

**Figure S5**. dPCR inhibition for each wastewater source **(A)** EBMUD; **(B)** WCWD; **(C)** SMCSD.

**Figure S6.** Virus concentration reported in mass gram basis across four methods with Solids reported in gc/gTSS and the rest in gc/mLWW

##### Supplementary Methods:

**Method A.** Acquisition and culturing of viruses for spiking

**Method B.** Measurement of the forms of virus in the stocks used for spiking

**Table S1.** Wastewater characteristics (metadata) on each sampling day

| Sampling date | Wastewater site | Flow rate (MGD) | Population | $BOD_5$ (mg/L) | TSS (mg/L) | pH | Per capita flow (L/person/ day) |
|---|---|---|---|---|---|---|---|
| 07/26/2023 | EBMUD | 46.9 | ~700,000 | 330.0 | 320.0 | 7.1 | 253.6 |
| 07/31/2023 | WCWD | 7.3 | ~70,000 | 332.0 | 556.0 | 7.6 | 394.8 |
| 08/02/2023 | SMCSD | 1.3 | ~18,000 | 170.0 | 220.0 | 7.3 | 273.4 |
| 08/30/2023 | EBMUD | 45.9 | ~700,000 | 340.0 | 440.0 | 6.7 | 247.6 |
| 09/05/2023 | WCWD | 7.1 | ~70,000 | 514.0 | 658.0 | 7.6 | 375.4 |
| 09/19/2023 | SMCSD | 1.2 | ~18,000 | 190.0 | 290.0 | 7.6 | 252.4 |

**Table S2.** Virus stocks and host cells information

| Virus family | Strain information | Source and catalog ID | Abbreviation in this manuscript | Host cells |
|---|---|---|---|---|
| Coronaviruses | SARS-Related Coronavirus 2, Isolate USA-WA1/2020, Heat Inactivated | BEI catalog no. NR-52286 | SARS-CoV-2 | N.A |
| | Human coronavirus OC43 strain unknown | BEI catalog no. NR-56241 | OC43 | Vero cells |
| Influenza A viruses | A/California/04/2009 (H1N1)pdm09 | BEI catalog no. NR-13658 | H1N1 | MDCK cells |
| | A/Netherlands/823/1992 (H3N2) | BEI catalog no. NR-49235 | H3N2 | MDCK cells |
| Enteroviruses | Human Coxsackievirus A6 strain Gdula | ATCC VR-1801 | CV-A6 | Rhabdomyosarcoma cells |
| | Human Coxsackievirus B5 strain Faulkner | ATCC VR-185 | CV-B5 | Vero cells |
| Adenoviruses | Human Adenovirus 2 strain unknown | Viral and Rickettsial Diseases Laboratory (VRDL); co-author strain collection | AdV2 | A549 cells |
| | Human Adenovirus 5 strain unknown | | AdV5 | A549 cells |

**Table S3:** Sample metadata, effective volume, and dPCR concentration (see excel file)

**Table S4.** dPCR forward primers, reverse primers, probes, and dyes of every virus target. MGB stands for minor groove binder which stabilizes probe-target hybridization and increases melting temperature, allowing the use of shorter probes

| Virus target | Duplex with | Forward primer sequence | Reverse primer sequence | Probe sequence | Dye |
|---|---|---|---|---|---|
| **SARS-CoV-2** | OC43 | GACCCCAAAATCAGCGAAAT | TCTGGTTACTGCCAGTTGAATCTG | ACCCCGCATTACGTTTGGTGGACC | FAM |
| **OC43** | SARS-CoV-2 | TATTGTTCCATGGGTTATGTAC | TCATGCACCTGGTCATAA | /56-TAMN/GG CGG TTT T/ZEN/G GAC ATG TTT ATG ATT T/3IABkFQ/ | TAMRA/ZEN/BkFQ |

| | | | | | |
|---|---|---|---|---|---|
| **H1N1** | H3N2 | TTACCAGATTTTGGCRATCTAYT | CCAGGGAGACTASCARTACCA | ACWGTYGCCAGTTC - MGB | FAM/MGB |
| **H3N2** | H1N1 | GCTCAGAGTGGGGAAAGCTATG | TTGGCATAGTCACGTTCAGC | CACGAATCAGATTACAA | VIC/MGB |
| **CV_A6** | CV_B5 | TGACGTGCTGAATGACACAG | AGCCCCTTGGTGAAATTTGC | TGTACCGCTCGGGCTTTTGC | VIC/MGB |
| **CV_B5** | CV_A6 | AAAACTGGCGCACATGAGAC | AACTGGCTCCGTGAATTTCC | AGACTTCACGCAGGACCCGG | FAM/MGB |
| **AdV2** | AdV5 | ACTAAACTTGGAGCGGGT | AGAACACCGTTTTGGTCAA | AAATGATGACAAACTTACCCTGTGGA | FAM |
| **AdV5** | AdV2 | TTGTGCCATCGGTCTACT | GTGGACCAGGTGTTTCAG | GACCTCCCGGCCACTATCC | Texas Red (ROX) |
| **PMMoV** | N.A | GAGTGGTTTGACCTTAACGTTTGA | TTGTCGGTTGCAATGCAAGT | CCTACCGAAGCAAATG | FAM/MGB |
| **Carjivirus** | N.A | CAGAAGTACAAACTCCTAAAAAACGTAGAG | GATGACCAATAAACAAGCCATTAGC | AATAACGATTTACGTGATGTAAC | Texas Red (ROX) |

**Table S5.** dPCR reaction mix recipe and thermal cycling conditions

| **DNA virus assays** | 8.5k plate | | 26k plate | |
|---|---|---|---|---|
| | singpleplex | duplex | singpleplex | duplex |
| Sample template (ul) | 3 | 3 | 10 | 10 |
| 4x Probe PCR Master Mix (ul) | 3 | 3 | 10 | 10 |
| 10X Primer/probe mix (ul) - virus 1 | 1.2 | 1.2 | 4 | 4 |
| 10X Primer/probe mix (ul) - virus 2 | / | 1.2 | / | 4 |
| Nuclease free H2O (ul) | 4.8 | 3.6 | 16 | 12 |
| Total (ul) | 12 | 12 | 40 | 40 |
| **RNA virus assays** | 8.5k plate | | 26k plate | |
| | singpleplex | duplex | singpleplex | duplex |
| Sample template (ul) | 3 | 3 | 10 | 10 |
| 4x Probe PCR Master Mix (ul) | 3 | 3 | 10 | 10 |
| 100X One-Step RT Mix (ul) | 0.12 | 0.12 | 0.4 | 0.4 |
| 20X Primer/probe mix (ul) - virus 1 | 0.6 | 0.6 | 2 | 2 |
| 20X Primer/probe mix (ul) - virus 2 | / | 0.6 | / | 2 |
| Nuclease free H2O (ul) | 5.28 | 4.68 | 17.6 | 15.6 |
| Total (ul) | 12 | 12 | 40 | 40 |

| **# cycles** | **Temperature °C** | **Duration** |
|---|---|---|
| 1 X | 50 | 40 min |
| 1 X | 95 | 2 min |
| 40 X | 95 | 5 s |
| | 60 | 30 s |

**Table S6.** Environmental Microbiology Minimum Information (EMMI) Checklist for dPCR

### Environmental Microbiology Minimum Information Checklist

#### Study Description

Study: Benchmarking concentration
Date: 22-Sep-2024
Completed by: Audrey Li-Wen Wa

| Environmental Sampling | Sample Treatment | Sample Reduction | Nucleic Acid Extraction | Reverse Transcription | PCR Detection | Analysis |
|---|---|---|---|---|---|---|
| Composite raw wastewater sampling | [x] Performed<br>Protease or Tween20 followed by centrifugation | [x] Performed<br>Concentration by affinity-based bead capture, ultrafiltration, or centrifugation | - Silica column<br>- Heat / mechanical / enzymatic lysis | [x] Performed<br>1-step RT | [ ] qPCR [x] dPCR<br>dPCR | - Threshold adjustment<br>- Data analysis |

#### Control Checklist

| | Environmental Sampling | Sample Treatment | Sample Reduction | Nucleic Acid Extraction | Reverse Transcription | PCR Detection |
|---|---|---|---|---|---|---|
| Step performed | [x] | [x] | [x] | [x] | [x] | [x] |
| **Negative Controls** | | | | | | |
| Step has control info | [ ] | [ ] | [ ] | [ ] | [ ] | [x] |
| # control replicates | 0 | 0 | 0 | 0 | 0 | 2 |
| Control result reported | [ ] | [ ] | [ ] | [ ] | [ ] | [x] |
| Data handling reported | [ ] | [ ] | [ ] | [ ] | [ ] | [x] |
| **Positive Controls** | | | | | | |
| Control introduced | [ ] | [ ] | [ ] | [ ] | [ ] | [x] |
| Internal/External | Internal | Internal | Internal | Internal | Internal | External |
| Independent/Parallel | Independent | Independent | Independent | Independent | Independent | Parallel |
| Step has control info | [ ] | [ ] | [ ] | [ ] | [ ] | [x] |
| # control replicates | 0 | 0 | 0 | 0 | 0 | 2 |
| Control result reported | [ ] | [ ] | [ ] | [ ] | [ ] | [x] |
| Data Handling reported | [ ] | [ ] | [ ] | [ ] | [ ] | [x] |

#### Process Checklist

##### Environmental Sampling
- [x] Sampling Procedure
- [x] Number of samples
- [x] Sample amount, mean, range
- [x] Sampling locations, dates, times

##### Sample Treatment
- [x] Performed
- [x] Treatment procedure
- [x] Reagents

##### Sample Reduction
- [x] Performed
- [x] Reduction procedure
- [x] Reagents
- [x] Concentration Factor

##### Nucleic Acid Extraction
- [x] Extraction procedure
- [x] Amount extracted, amount obtained
- [x] Extract storage conditions

##### qPCR or dPCR
- [ ] Target gene name, amplicon length
- [x] Thermocycling temperatures and times
- [x] Master mix: composition, vendors, concentrations
- [ ] Additives: vendors, concentrations
- [x] Template amount added, pre-treatment (if any)
- [x] Primers: sequences, concentrations, vendors, references
- [ ] Amplicon confirmation method (probe, melt curve, etc)
- [x] Probe sequence, concentration, vendor, reference
- [x] Instrumentation
- [x] Equivalent volume of sample analyzed by PCR
- [x] Inhibition assessment procedure
- [ ] Inhibition control description (if used)
- [x] Number samples tested and found inhibited

##### Reverse Transcription
- [x] Performed
- [x] One or two step
- [ ] cDNA storage conditions (if two step)
- [x] Reaction temperatures and times
- [x] Reaction reagents and concentrations
- [x] Priming method
- [x] Reaction volume, added template amount
- [x] Inhibition assessment procedure
- [ ] Inhibition control description (if used)
- [x] Number samples tested and found inhibited

##### Analysis – dPCR
- [x] Threshold settings
- [ ] Technical replicates, number, well merging
- [x] Partitions measured, number, mean, variance
- [x] Partition volume
- [ ] Target copies per partition, mean, variance
- [x] Program used for dPCR analysis
- [x] Explanation of control results, example plots

##### Analysis – qPCR
- [ ] Method for handling failed negative controls
- [ ] Technical replicates, number, calculations
- [ ] Calibration standards: description and source
- [ ] Method of quantifying standards
- [ ] Calibration curve slope
- [ ] Calibration curve R2
- [ ] Lowest standard measured or 95% LOD
- [ ] Cq value determination method

**Table S7:** Statistical analysis of (A-B) recovery efficiency, (C-D) purified total nucleic acid concentration, and (E-F) total gene copies in a 40mL wastewater sample. (A),(C),(E) p-values of Kruskal-Wallis H-tests that compare differences among concentrations and extraction methods within the same virus type from were shown, with three colors indicating different significance level: yellow is $P < 0.05$, orange is $P < 0.01$, and red is $P < 0.001$ (B),(D),(F) p-values of the Dunn test for pairwise comparisons with Bonferroni correction was further employed following a significant result in the Kruskal-Wallis test. Statistical significance was determined at a 95% confidence interval, with three colors also representing different levels of significance.

**(A)** Kruskal-Wallis test for Figure 2A and 2B (recovery efficiency)

| virus | p-value |
|---|---|
| SARS-CoV-2 | 8.20e-04 |
| H1N1 | 2.68e-03 |
| H3N2 | 1.09e-03 |
| OC43 | 8.54e-03 |
| AdV2 | 3.69e-01 |
| AdV5 | 2.82e-02 |
| CV_A6 | 4.37e-01 |
| CV_B5 | 1.04e-03 |

**(B)** Dunn Test for Figure 2A and 2B (recovery efficiency)

| AdV5 | Promega | InnovaPrep | Solids | Nanotrap |
|---|---|---|---|---|
| Promega | 1.00e+00 | 1.00e+00 | 2.91e-02 | 1.48e-01 |
| InnovaPrep | 1.00e+00 | 1.00e+00 | 7.85e-01 | 1.00e+00 |
| Solids | 2.91e-02 | 7.85e-01 | 1.00e+00 | 1.00e+00 |
| Nanotrap | 1.48e-01 | 1.00e+00 | 1.00e+00 | 1.00e+00 |

| CV-B5 | Promega | InnovaPrep | Solids | Nanotrap |
|---|---|---|---|---|
| Promega | 1.00e+00 | 1.00e+00 | 1.65e-01 | 1.51e-02 |
| InnovaPrep | 1.00e+00 | 1.00e+00 | 6.82e-02 | 4.89e-03 |
| Solids | 1.65e-01 | 6.82e-02 | 1.00e+00 | 1.00e+00 |
| Nanotrap | 1.51e-02 | 4.89e-03 | 1.00e+00 | 1.00e+00 |

| SARS-CoV-2 | Promega | InnovaPrep | Solids | Nanotrap |
|---|---|---|---|---|
| Promega | 1.00e+00 | 7.66e-02 | 1.22e-03 | 1.00e+00 |
| InnovaPrep | 7.66e-02 | 1.00e+00 | 1.00e+00 | 5.65e-01 |
| Solids | 1.22e-03 | 1.00e+00 | 1.00e+00 | 2.25e-02 |
| Nanotrap | 1.00e+00 | 5.65e-01 | 2.25e-02 | 1.00e+00 |

| OC43 | Promega | InnovaPrep | Solids | Nanotrap |
|---|---|---|---|---|
| Promega | 1.00e+00 | 1.00e+00 | 4.89e-03 | 1.00e+00 |
| InnovaPrep | 1.00e+00 | 1.00e+00 | 1.83e-01 | 1.00e+00 |
| Solids | 4.89e-03 | 1.83e-01 | 1.00e+00 | 1.83e-01 |
| Nanotrap | 1.00e+00 | 1.00e+00 | 1.83e-01 | 1.00e+00 |

| H1N1 | Promega | InnovaPrep | Solids | Nanotrap |
|---|---|---|---|---|
| Promega | 1.00e+00 | 4.28e-02 | 1.75e-03 | 2.79e-01 |
| InnovaPrep | 4.28e-02 | 1.00e+00 | 1.00e+00 | 1.00e+00 |
| Solids | 1.75e-03 | 1.00e+00 | 1.00e+00 | 6.17e-01 |
| Nanotrap | 2.79e-01 | 1.00e+00 | 6.17e-01 | 1.00e+00 |

| H3N2 | Promega | InnovaPrep | Solids | Nanotrap |
|---|---|---|---|---|
| Promega | 1.00e+00 | 2.63e-01 | 3.63e-04 | 2.50e-01 |
| InnovaPrep | 2.63e-01 | 1.00e+00 | 2.76e-01 | 1.00e+00 |
| Solids | 3.63e-04 | 2.76e-01 | 1.00e+00 | 2.90e-01 |
| Nanotrap | 2.50e-01 | 1.00e+00 | 2.90e-01 | 1.00e+00 |

| Carjivirus | Promega | InnovaPrep | Solids | Nanotrap |
|---|---|---|---|---|
| Promega | 1.00e+00 | 5.19e-02 | 3.61e-01 | 2.44e-01 |
| InnovaPrep | 5.19e-02 | 1.00e+00 | 1.00e+00 | 2.64e-03 |
| Solids | 3.61e-01 | 1.00e+00 | 1.00e+00 | 8.52e-01 |
| Nanotrap | 2.44e-01 | 2.64e-03 | 8.52e-01 | 1.00e+00 |

| PMMoV | Promega | InnovaPrep | Solids | Nanotrap |
|---|---|---|---|---|
| Promega | 1.00e+00 | 4.35e-01 | 1.22e-03 | 1.00e+00 |
| InnovaPrep | 4.35e-01 | 1.00e+00 | 3.30e-01 | 1.00e+00 |
| Solids | 1.22e-03 | 3.30e-01 | 1.00e+00 | 1.07e-01 |
| Nanotrap | 1.00e+00 | 1.00e+00 | 1.07e-01 | 1.00e+00 |

**(C)** Kruskal-Wallis test for Figure 2C and 2D (conc. in purified TNA)

| virus | p-value |
|---|---|
| AdV2 | 1.03e-01 |
| AdV5 | 1.26e-02 |
| CV_A6 | 9.26e-02 |
| CV_B5 | 7.70e-04 |
| SARS-CoV-2 | 2.09e-04 |
| OC43 | 2.69e-03 |
| H1N1 | 3.92e-03 |
| H3N2 | 3.99e-03 |

**(D)** Dunn test for Figure 2C and 2D (conc. in purified TNA)

| AdV5 | Promega | InnovaPrep | Solids | Nanotrap |
|---|---|---|---|---|
| Promega | 1.00e+00 | 1.00e+00 | 7.55e-03 | 1.00e+00 |
| InnovaPrep | 1.00e+00 | 1.00e+00 | 3.00e-01 | 1.00e+00 |
| Solids | 7.55e-03 | 3.00e-01 | 1.00e+00 | 1.83e-01 |
| Nanotrap | 1.00e+00 | 1.00e+00 | 1.83e-01 | 1.00e+00 |

| CV-B5 | Promega | InnovaPrep | Solids | Nanotrap |
|---|---|---|---|---|
| Promega | 1.00e+00 | 1.00e+00 | 1.73e-02 | 1.97e-02 |
| InnovaPrep | 1.00e+00 | 1.00e+00 | 2.56e-02 | 2.91e-02 |
| Solids | 1.73e-02 | 2.56e-02 | 1.00e+00 | 1.00e+00 |
| Nanotrap | 1.97e-02 | 2.91e-02 | 1.00e+00 | 1.00e+00 |

| SARS-CoV-2 | Promega | InnovaPrep | Solids | Nanotrap |
|---|---|---|---|---|
| Promega | 1.00e+00 | 1.96e-02 | 3.73e-04 | 1.00e+00 |
| InnovaPrep | 1.96e-02 | 1.00e+00 | 1.00e+00 | 3.61e-01 |
| Solids | 3.73e-04 | 1.00e+00 | 1.00e+00 | 1.96e-02 |
| Nanotrap | 1.00e+00 | 3.61e-01 | 1.96e-02 | 1.00e+00 |

| OC43 | Promega | InnovaPrep | Solids | Nanotrap |
|---|---|---|---|---|
| Promega | 1.00e+00 | 1.09e-01 | 2.20e-03 | 6.17e-01 |
| InnovaPrep | 1.09e-01 | 1.00e+00 | 1.00e+00 | 1.00e+00 |
| Solids | 2.20e-03 | 1.00e+00 | 1.00e+00 | 3.20e-01 |
| Nanotrap | 6.17e-01 | 1.00e+00 | 3.20e-01 | 1.00e+00 |

| H1N1 | Promega | InnovaPrep | Solids | Nanotrap |
|---|---|---|---|---|
| Promega | 1.00e+00 | 5.19e-02 | 2.64e-03 | 2.44e-01 |
| InnovaPrep | 5.19e-02 | 1.00e+00 | 1.00e+00 | 1.00e+00 |
| Solids | 2.64e-03 | 1.00e+00 | 1.00e+00 | 8.52e-01 |
| Nanotrap | 2.44e-01 | 1.00e+00 | 8.52e-01 | 1.00e+00 |

| H3N2 | Promega | InnovaPrep | Solids | Nanotrap |
|---|---|---|---|---|
| Promega | 1.00e+00 | 4.35e-01 | 1.22e-03 | 1.00e+00 |
| InnovaPrep | 4.35e-01 | 1.00e+00 | 3.30e-01 | 1.00e+00 |
| Solids | 1.22e-03 | 3.30e-01 | 1.00e+00 | 1.07e-01 |
| Nanotrap | 1.00e+00 | 1.00e+00 | 1.07e-01 | 1.00e+00 |

| Carjivirus | Promega | InnovaPrep | Solids | Nanotrap |
|---|---|---|---|---|
| Promega | 1.00e+00 | 5.19e-02 | 3.61e-01 | 2.44e-01 |
| InnovaPrep | 5.19e-02 | 1.00e+00 | 1.00e+00 | 2.64e-03 |
| Solids | 3.61e-01 | 1.00e+00 | 1.00e+00 | 8.52e-01 |
| Nanotrap | 2.44e-01 | 2.64e-03 | 8.52e-01 | 1.00e+00 |

| PMMoV | Promega | InnovaPrep | Solids | Nanotrap |
|---|---|---|---|---|
| Promega | 1.00e+00 | 2.63e-01 | 3.63e-04 | 2.50e-01 |
| InnovaPrep | 2.63e-01 | 1.00e+00 | 2.76e-01 | 1.00e+00 |
| Solids | 3.63e-04 | 2.76e-01 | 1.00e+00 | 2.90e-01 |
| Nanotrap | 2.50e-01 | 1.00e+00 | 2.90e-01 | 1.00e+00 |

**(E)** Kruskal-Wallis test for Figure S2C (total gene copies in a 40mL WW sample)

| virus | p-value |
|---|---|
| AdV2 | 2.12e-01 |
| AdV5 | 5.20e-02 |
| CV_A6 | 4.16e-01 |
| CV_B5 | 8.76e-04 |
| SARS-CoV-2 | 2.09e-04 |
| OC43 | 4.13e-03 |
| H1N1 | 7.86e-03 |
| H3N2 | 8.66e-03 |

**(F)** Dunn test for Figure S2C (total gene copies in a 40mL WW sample)

| CV-B5 | Promega | InnovaPrep | Solids | Nanotrap |
|---|---|---|---|---|
| Promega | 1.00e+00 | 1.00e+00 | 6.82e-02 | 2.56e-02 |
| InnovaPrep | 1.00e+00 | 1.00e+00 | 2.56e-02 | 8.71e-03 |
| Solids | 6.82e-02 | 2.56e-02 | 1.00e+00 | 1.00e+00 |
| Nanotrap | 2.56e-02 | 8.71e-03 | 1.00e+00 | 1.00e+00 |

| SARS-CoV-2 | Promega | InnovaPrep | Solids | Nanotrap |
|---|---|---|---|---|
| Promega | 1.00e+00 | 3.71e-02 | 4.43e-04 | 1.00e+00 |
| InnovaPrep | 3.71e-02 | 1.00e+00 | 1.00e+00 | 6.66e-01 |
| Solids | 4.43e-04 | 1.00e+00 | 1.00e+00 | 2.89e-02 |
| Nanotrap | 1.00e+00 | 6.66e-01 | 2.89e-02 | 1.00e+00 |

| OC43 | Promega | InnovaPrep | Solids | Nanotrap |
|---|---|---|---|---|
| Promega | 1.00e+00 | 9.18e-01 | 1.96e-03 | 1.00e+00 |
| InnovaPrep | 9.18e-01 | 1.00e+00 | 1.83e-01 | 1.00e+00 |
| Solids | 1.96e-03 | 1.83e-01 | 1.00e+00 | 1.48e-01 |
| Nanotrap | 1.00e+00 | 1.00e+00 | 1.48e-01 | 1.00e+00 |

| H1N1 | Promega | InnovaPrep | Solids | Nanotrap |
|---|---|---|---|---|
| Promega | 1.00e+00 | 1.57e-01 | 4.32e-03 | 2.44e-01 |
| InnovaPrep | 1.57e-01 | 1.00e+00 | 1.00e+00 | 1.00e+00 |
| Solids | 4.32e-03 | 1.00e+00 | 1.00e+00 | 1.00e+00 |
| Nanotrap | 2.44e-01 | 1.00e+00 | 1.00e+00 | 1.00e+00 |

| H3N2 | Promega | InnovaPrep | Solids | Nanotrap |
|---|---|---|---|---|
| Promega | 1.00e+00 | 3.20e-01 | 4.17e-03 | 7.33e-01 |
| InnovaPrep | 3.20e-01 | 1.00e+00 | 8.66e-01 | 1.00e+00 |
| Solids | 4.17e-03 | 8.66e-01 | 1.00e+00 | 3.89e-01 |
| Nanotrap | 7.33e-01 | 1.00e+00 | 3.89e-01 | 1.00e+00 |

**(A)**

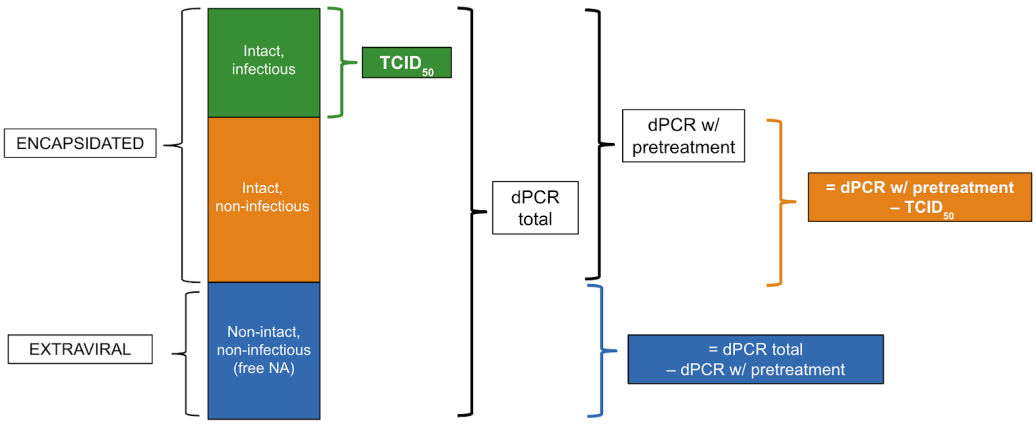


**(B)**

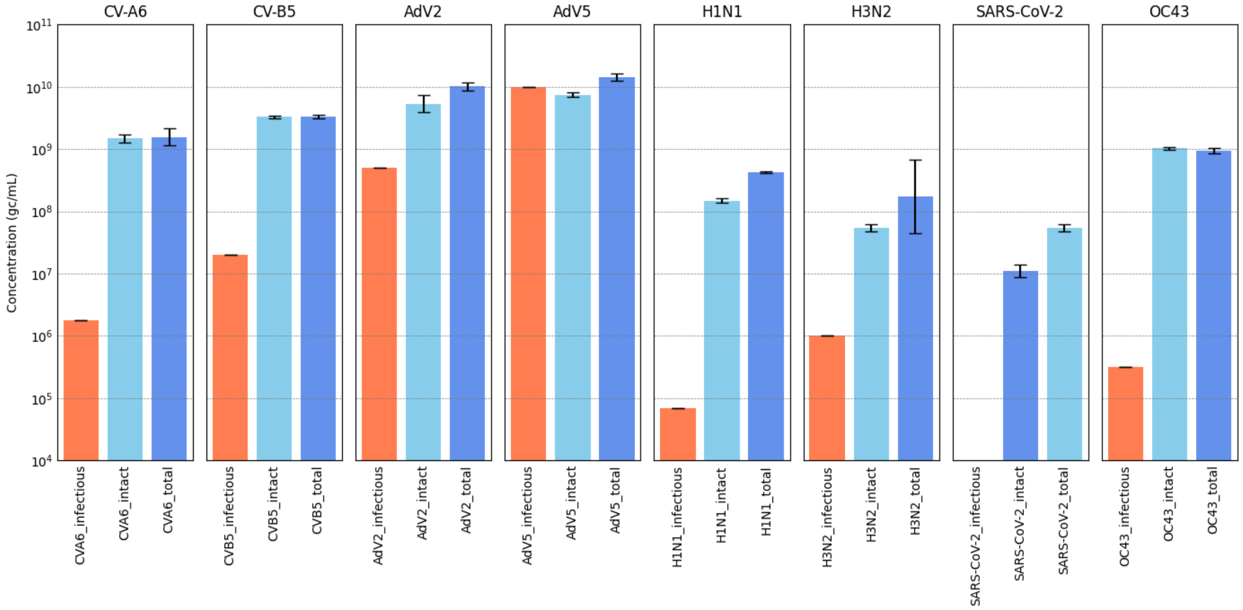


**Figure S1** (**A**) Schematic of the calculation of different forms of viruses. (**B**) Concentrations of infectious virus from TCID50/mL (orange), intact virus concentration (light blue) measured after DNAse/RNase treatment pretreatment, and total virus concentration without any treatment (dark blue). TCID50/mL was not determined for SARS-CoV-2, which was purchased as an inactivated stock.

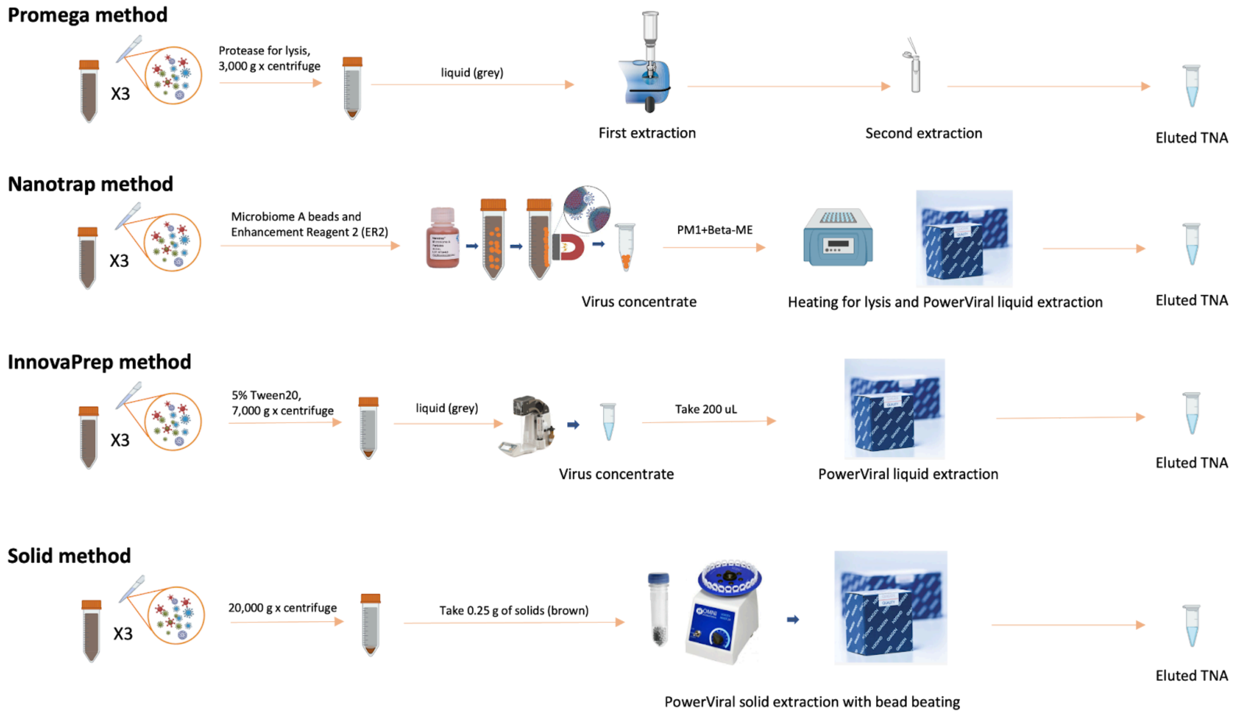


**Figure S2.** Schematic workflow of each method. The workflow is modified based on a SI Figure in a complementary paper[1].

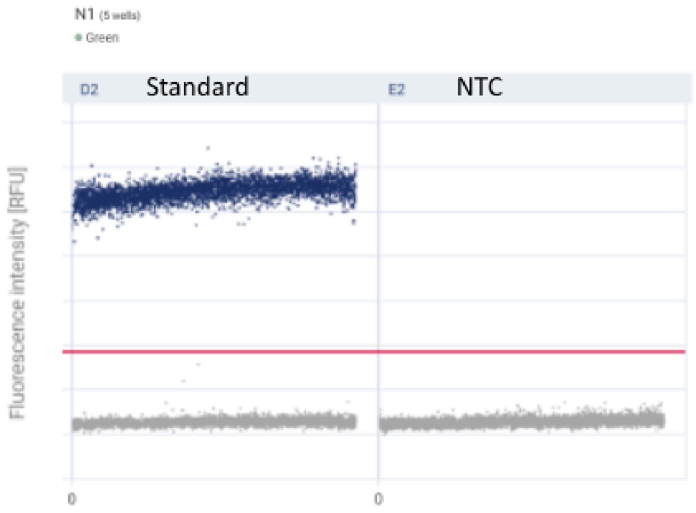


**Figure S3.** Examples of partition fluorescence plots of dPCR positive and negative control for the CDC N1 assay for SARS-CoV-2

(A)

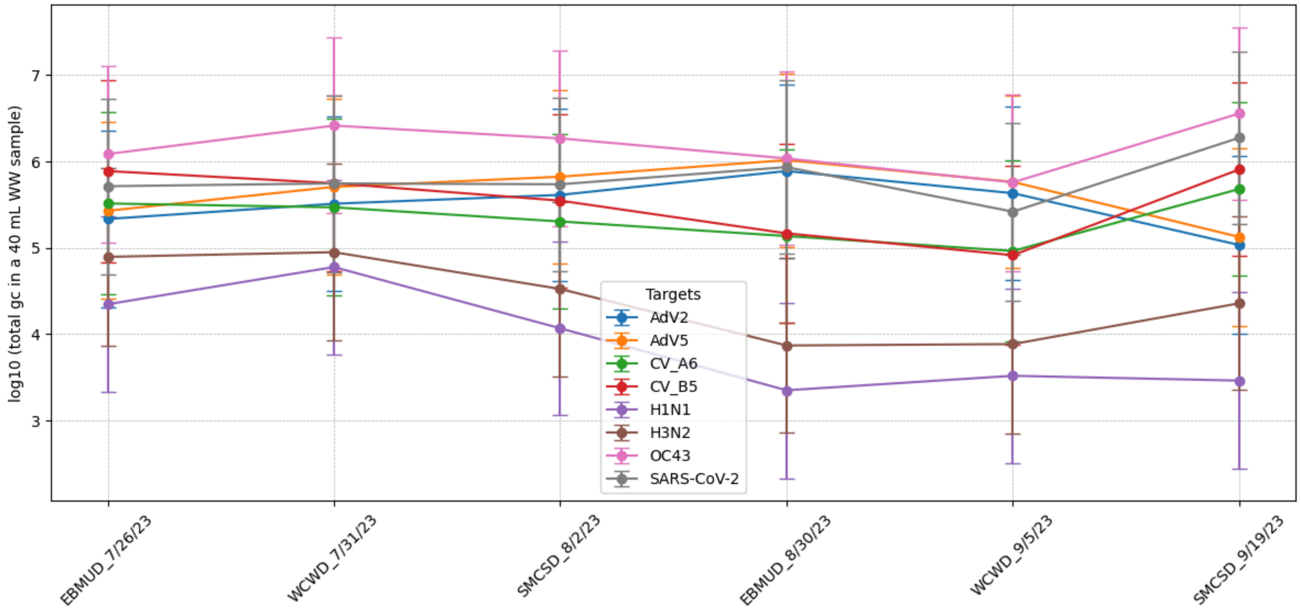

log10 (total gc in a 40 mL WW sample)
Targets
AdV2
AdV5
CV_A6
CV_B5
H1N1
H3N2
OC43
SARS-CoV-2
EBMUD_7/26/23
WCWD_7/31/23
SMCSD_8/2/23
EBMUD_8/30/23
WCWD_9/5/23
SMCSD_9/19/23


(B)

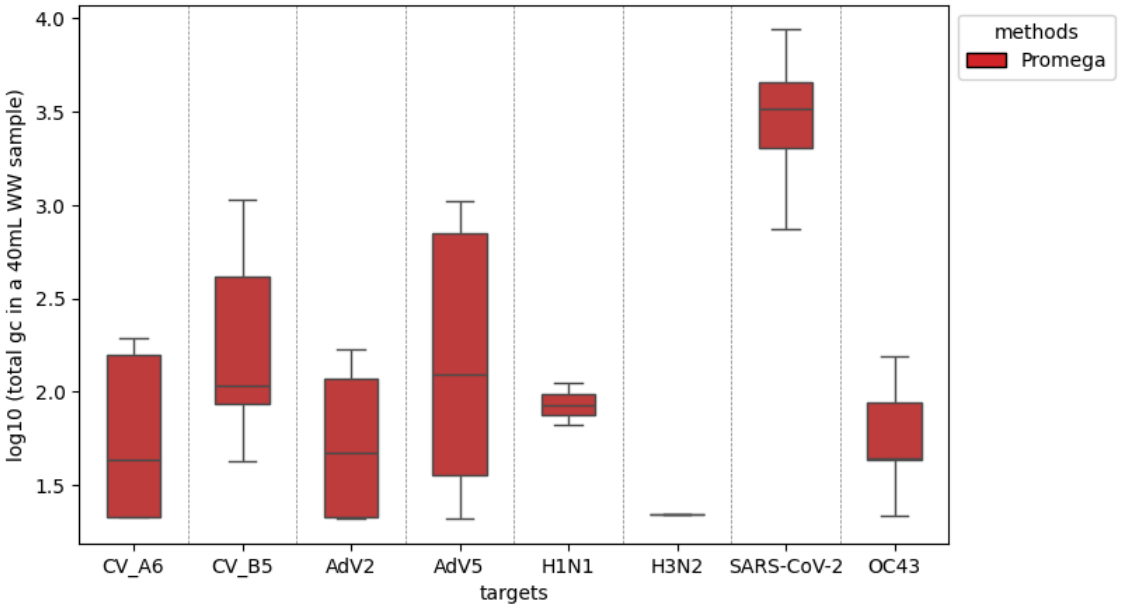

log10 (total gc in a 40mL WW sample)
methods
Promega
CV_A6
CV_B5
AdV2
AdV5
H1N1
H3N2
SARS-CoV-2
OC43
targets

**(C)**

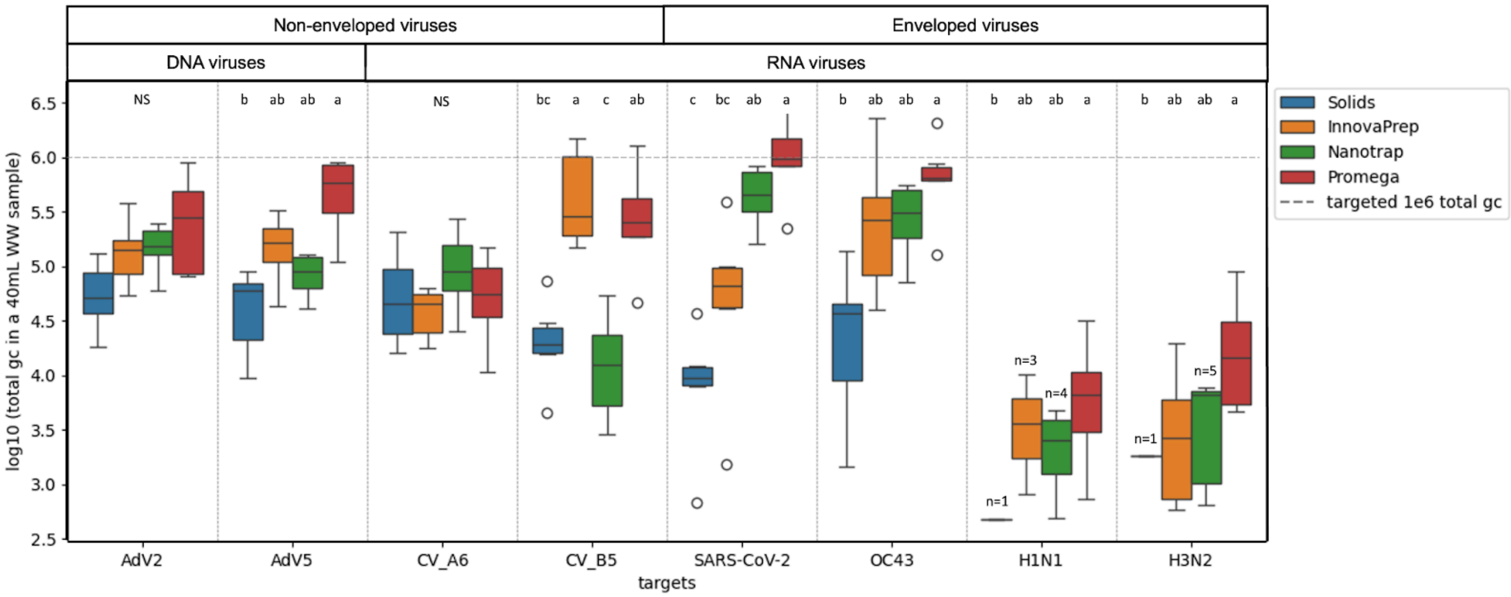


**Figure S4.** (**A**) Initial virus gene copies in 40 mL of wastewater based on the concentration of the pure virus cocktail. The targeted spiked-in gene copies were $10^6$. The pure virus cocktail was quantified after extraction with the Qiagen AllPrep PowerViral extraction kit with carrier RNA addition. Values shown are the geometric mean and standard deviation of triplicate extractions. (**B**) Endogenous virus gene copies in 40 mL of wastewater were quantified from the unspiked samples using Promega for extraction. **(C)** Total gene copies (gc) of spiked-in viruses recovered from a 40 mL of wastewater sample across four concentration / extraction methods (colors); note log10 scale of y-axis. Boxes and whiskers indicate the interquartile range (IQR) and minimum / maximum values within 1.5 times the IQR, respectively, across all samples (n=6 for two time points and three wastewater sources; three biological replicates were averaged first by calculating the geometric mean before plotting). Outliers are displayed as individual points. The initial spike-in was targeted to be $10^6$ gene copies per virus in each wastewater sample (gray dashed line). Significance letters are assigned based on the results of the post-hoc Dunn's test for pairwise comparisons of methods within each virus type. Any method sharing the same letter is not significantly different from each other.

**(A) EBMUD**

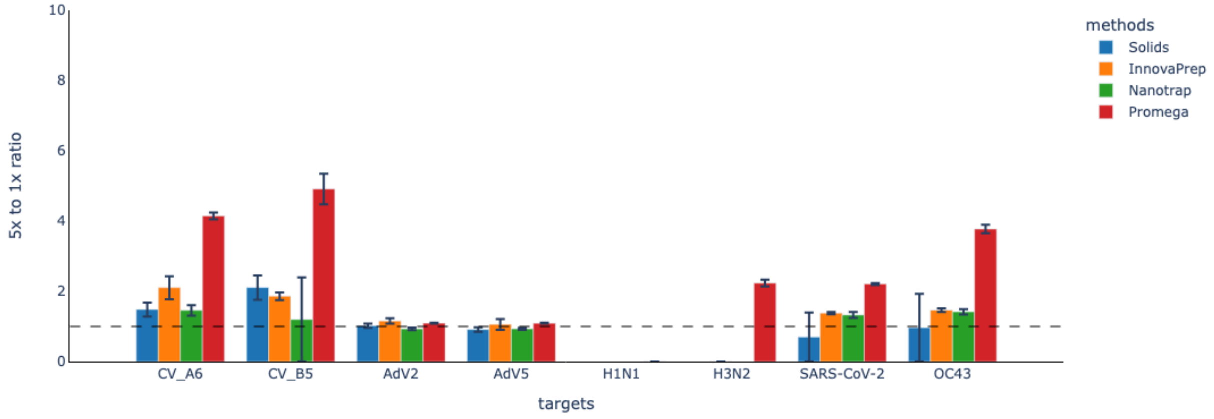


**(B) WCWD**

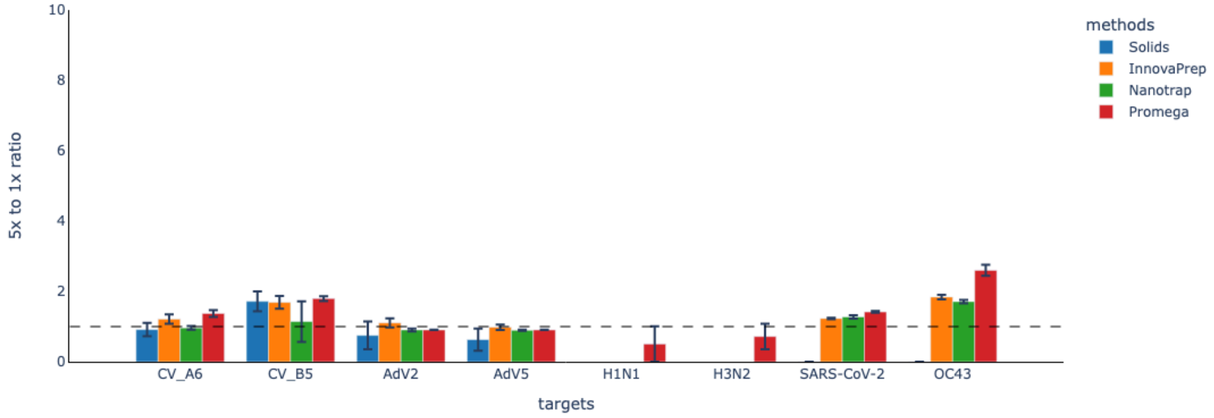


**(C) SMCSD**

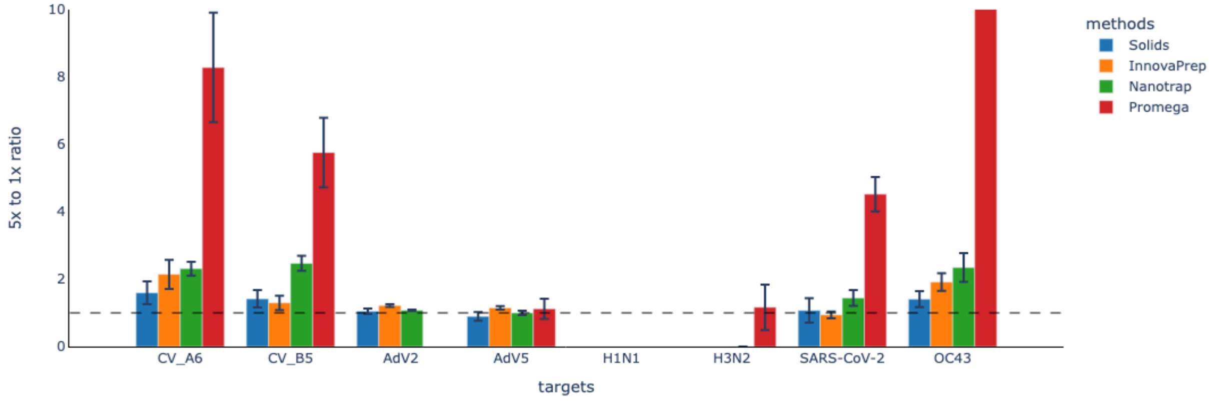


**Figure S5.** dPCR inhibition for each wastewater source **(A)** EBMUD; **(B)** WCWD; **(C)** SMCSD. The purified nucleic acids from each method from time point two were analyzed by dPCR with and without 5-fold dilution. If there are inhibitors present, diluting the sample to 5x reduces the inhibitor concentration, allowing the assay to detect more target TNA. After back-calculating (multiplying the diluted measurement by 5), we should observe a higher concentration if inhibition was present in the undiluted (1x) sample. Therefore, a ratio of one indicates no inhibition, while higher ratios indicate inhibition. When concentrations were undetectable for either 5x dilution or both 1x and 5x dilutions, the bars are not shown in the figure. Note that the undiluted concentrations of several samples were already low (Figure 2) so that the five-fold dilution could result in undetectable signal.

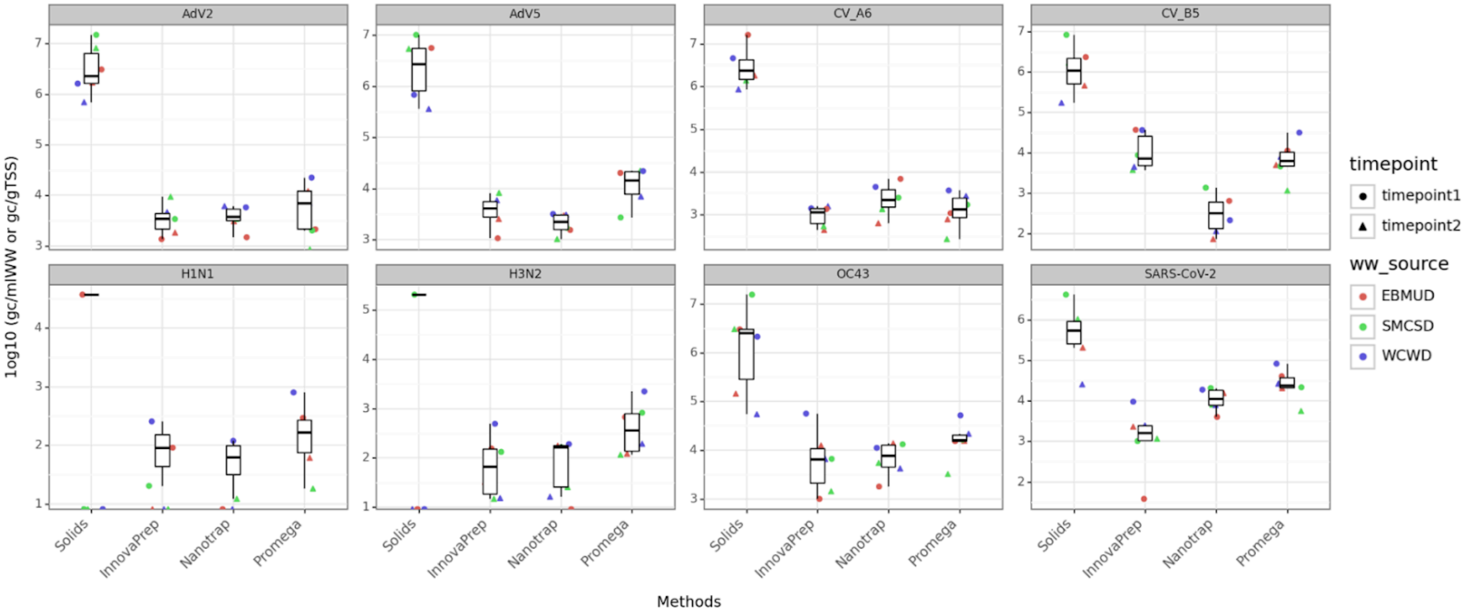


**Figure S6.** Virus concentration reported per mass across four methods with Solids reported in gc/gTSS and the rest in gc/mLWW (note different y-axes for each virus). To calculate the Solids concentration per mass, the TSS concentration (mg/L) provided by the wastewater treatment plant for each sampling date was used. While this mass may not exactly equal that of the solids pellet produced in the laboratory by centrifuging each 40-mL wastewater sample at 20,000 x g, it is likely a good approximation.

**Method A. Acquisition and culturing of viruses for spiking:**
Adenoviruses Type 2 and 5 were propagated using A549 cells (ATCC CCL-185) at 37°C using Dulbecco's Modified Eagle's Medium (DMEM, Thermo Fisher) with 5% fetal bovine serum (FBS) and penicillin/streptomycin (P/S, 100 U/mL Penicillin & 100 µg/mL Streptomycin; Millipore Sigma). Viral titer was determined via median tissue culture infectious dose ($TCID_{50}$) using A549 cells and calculated according to the Spearman–Kärber method. Human coronavirus OC43 (ATCC NR-56241) was propagated on MRC-5 cells (ATCC CCL-171) at 33°C, using DMEM with 5% FBS and P/S according to a previous study[2]. The OC43 titer was determined via $TCID_{50}$ using Vero/TMPRSS2 cells at 37°C using DMEM with 5% FBS and P/S. Coxsackie A6 virus (ATCC VR-1801) was grown on Rhabdomyosarcoma cells (RD; ATCC CCL-136) at 37°C using DMEM with 5% FBS and P/S. The $TCID_{50}$ of CV-A6 was obtained using RD cells. Coxsackie B5 virus (ATCC VR-185) was grown on Vero E6 cells (ATCC CRL-1586) at 37°C using DMEM with 5% FBS and P/S. The $TCID_{50}$ of CV-B5 was obtained using Vero E6 cells. Influenza A virus A/California/04/2009 (H1N1pdm09) and A/Netherlands/823/1992 (H3N2) were grown on MDCK cells (ATCC Cat. No. CCL-34) at 37°C using DMEM with 0.2% BSA, 25 mM HEPES, and 50 U/mL penicillin/50 ug/mL streptomycin (1X). MDCK cells were washed with PBS three times prior to viral infection to remove FBS which would infer with viral production. A concentration of 0.5-1 ug/mL of TPCK-treated trypsin (Sigma Cat. No. T-8642) was added to enhance the infection of influenza viruses in MDCK cells, by cleaving the hemagglutinin precursor protein enabling viral entry. The $TCID_{50}$ of both H1N1 and H3N2 were obtained using MDCK cells.

For all viruses after sufficient time for viral infection of cells (significant cell lysis), the flask contents were removed and centrifuged at 1,000 to 2,800 x g for 5-10 min at 4°C to pellet cell debris. The supernatants were then aliquoted into 1-mL cryotubes and stored at -80C. In addition, coxsackieviruses and adenoviruses were put through freeze/thaw cycles to release virions from cells prior to centrifugation.Finally, heat-inactivated SARS-CoV-2 (isolate: USA-WA1/2020, NR-52286) was obtained from Biodefense and Emerging Infections (BEI) Resources.

**Method B. Measurement of the forms of virus in the stocks used for spiking:**
The DNase pretreatment was performed using a previously established method[3,4]. Enzyme storage buffer consisted of 10 mM Tris-HCl (pH 8.0) and 2 mM $CaCl_2$ in 50% glycerol, and the 10X reaction buffer contained 100 mM Tris-HCl (pH 8.0), 25 mM $MgCl_2$, and 5 mM $CaCl_2$ in Milli-Q water. DNase I, grade II from bovine pancreas (Roche) was resuspended into 5 mL of the storage buffer at a concentration of 40,000 U $mL^{-1}$ and stored at -20 ℃. Immediately before DNase pretreatment of virus stocks, DNase I in storage buffer (40,000 U $mL^{-1}$) was diluted 1:40 in the 10X reaction buffer and gently mixed to obtain a 1,000 U $mL^{-1}$ working DNase solution. The DNase working solution (1000 U/mL) was spiked into virus stock samples to reach a concentration of 200 U/mL and incubated for 30 minutes at 37 °C without shaking. Untreated samples were incubated without DNase I at 37°C for 30 minutes without shaking. The DNase reaction was inactivated by adding 5 µL of 100 mM EDTA and 5 µL of 100 mM EGTA (per 200 µL of sample) and incubating samples for 5-10 minutes at 65-75 °C. The RNase pretreatment was performed according to an established procedure[3,5]. 220 µL of each virus stock were treated with 20 units of RNase ONE Ribonuclease (10 U $\mu L^{-1}$) (Promega). All samples were incubated at 37 °C for 15 min with shaking.

[1] Jiang M, Wang ALW, Be NA, Mulakken N, Nelson KL, Kantor RS. Evaluation of the Impact of Concentration and Extraction Methods on the Targeted Sequencing of Human Viruses from Wastewater. Environ Sci Technol. 2024;58(19):8239-8250. doi:10.1021/acs.est.4c00580

[2] Shafagati, N., Fite, K., Patanarut, A., Baer, A., Pinkham, C., An, S., ... & Kehn-Hall, K. (2016). Enhanced detection of respiratory pathogens with nanotrap particles. Virulence, 7(7), 756-769. http://dx.doi.org/10.1080/21505594.2016.1185585

[3] Harrison, K. R.; Snead, D.; Kilts, A.; Ammerman, M. L.; Wigginton, K. R. The Protective Effect of Virus Capsids on RNA and DNA Virus Genomes in Wastewater. medRxiv May 21, 2023, p 2023.05.19.23290245. https://doi.org/10.1101/2023.05.19.23290245.

[4] Thornton, J. E. DNase I Treatment. protocols.io 2015.

[5] Rockey, N.; Young, S.; Kohn, T.; Pecson, B.; Wobus, C.; Raskin, L.; Wigginton, K. UV Disinfection of Human Norovirus: Evaluating Infectivity Using a Genome-Wide PCR-Based Approach. ENVIRONMENTAL SCIENCE & TECHNOLOGY 2020, 54 (5), 2851–2858. https://doi.org/10.1021/acs.est.9b05747.
