## Supplementary_TableS3 for "Benchmarking concentration and direct extraction methods for wastewater-based surveillance of eight human respiratory viruses: implications for rapid application to novel pathogens"

### Supplementary Table S3: Sample metadata, effective volume, and dPCR concentration

| date_sampling | ww_source | timepoint | biological replicate | concentration_methods | sample_id | empty_tube_mass_g | full_tube_mass_g | concentrated_mass_g | extraction_vol_uL_or_mg | elution_vol_uL | cocktail_extr_vol_uL | Effective_vol_mL | notes |
|---|---|---|---|---|---|---|---|---|---|---|---|---|---|
| 7/26/23 | EBMUD | 1 | 1 | cocktail | EBMUD_cocktail_1_072623 | | | 0.2 | 200 | 100 | 40 | | |
| 7/26/23 | EBMUD | 1 | 2 | cocktail | EBMUD_cocktail_2_072623 | | | 0.2 | 200 | 100 | 40 | | |
| 7/31/23 | WCSD | 1 | 1 | cocktail | WCSD_cocktail_1_073123 | | | 0.2 | 200 | 100 | 200 | | |
| 7/31/23 | WCSD | 1 | 2 | cocktail | WCSD_cocktail_2_073123 | | | 0.2 | 200 | 100 | 200 | | |
| 8/2/23 | SMCSD | 1 | 1 | cocktail | SMCSD_cocktail_1_080223 | | | 0.2 | 200 | 100 | 40 | | |
| 8/2/23 | SMCSD | 1 | 2 | cocktail | SMCSD_cocktail_2_080223 | | | 0.2 | 200 | 100 | 40 | | |
| 8/30/23 | EBMUD | 2 | 1 | cocktail | EBMUD_cocktail_1_083023 | | | 0.2 | 200 | 100 | 40 | | |
| 8/30/23 | EBMUD | 2 | 2 | cocktail | EBMUD_cocktail_2_083023 | | | 0.2 | 200 | 100 | 20 | | |
| 9/5/23 | WCSD | 2 | 1 | cocktail | WCSD_cocktail_1_090523 | | | 0.2 | 200 | 100 | 40 | | |
| 9/5/23 | WCSD | 2 | 2 | cocktail | WCSD_cocktail_2_090523 | | | 0.2 | 200 | 100 | 20 | | |
| 9/19/23 | SMCSD | 2 | 1 | cocktail | SMCSD_cocktail_1_091923 | | | 0.2 | 200 | 100 | 40 | | |
| 9/19/23 | SMCSD | 2 | 2 | cocktail | SMCSD_cocktail_2_091923 | | | 0.2 | 200 | 100 | 40 | | |
| 8/30/23 | EBMUD | 2 | 1 | cocktail | EBMUD_cocktail_1_083023_INB | | | 0.2 | 200 | 100 | 40 | | |
| 8/30/23 | EBMUD | 2 | 2 | cocktail | EBMUD_cocktail_2_083023_INB | | | 0.2 | 200 | 100 | 20 | | |
| 9/5/23 | WCSD | 2 | 1 | cocktail | WCSD_cocktail_1_090523_INB | | | 0.2 | 200 | 100 | 40 | | |
| 9/5/23 | WCSD | 2 | 2 | cocktail | WCSD_cocktail_2_090523_INB | | | 0.2 | 200 | 100 | 20 | | |
| 9/19/23 | SMCSD | 2 | 1 | cocktail | SMCSD_cocktail_1_091923_INB | | | 0.2 | 200 | 100 | 40 | | |
| 9/19/23 | SMCSD | 2 | 2 | cocktail | SMCSD_cocktail_2_091923_INB | | | 0.2 | 200 | 100 | 40 | | |
| 9/19/23 | SMCSD | 2 | 1 | cocktail | SMCSD_cocktail_1_091923 | | | 0.2 | 200 | 100 | 40 | | |
| 9/19/23 | SMCSD | 2 | 2 | cocktail | SMCSD_cocktail_2_091923 | | | 0.2 | 200 | 100 | 40 | | |
| 7/26/23 | EBMUD | 1 | 1 | InnovaPrep | EBMUD_InnovaPrep_1_072623 | 14.21 | 14.93 | 0.72 | 200 | 100 | | 11.11 | |
| 7/26/23 | EBMUD | 1 | 2 | InnovaPrep | EBMUD_InnovaPrep_2_072623 | 14.22 | 14.74 | 0.52 | 200 | 100 | | 15.38 | |
| 7/26/23 | EBMUD | 1 | 3 | InnovaPrep | EBMUD_InnovaPrep_3_072623 | 14.16 | 14.63 | 0.47 | 200 | 100 | | 17.02 | |
| 7/31/23 | WCSD | 1 | 1 | InnovaPrep | WCSD_InnovaPrep_1_073123 | 14.23 | 14.69 | 0.46 | 200 | 100 | | 17.39 | |
| 7/31/23 | WCSD | 1 | 2 | InnovaPrep | WCSD_InnovaPrep_2_073123 | 14.06 | 14.64 | 0.58 | 200 | 100 | | 13.79 | |
| 7/31/23 | WCSD | 1 | 3 | InnovaPrep | WCSD_InnovaPrep_3_073123 | 14.1 | 14.78 | 0.68 | 200 | 100 | | 11.76 | |
| 8/2/23 | SMCSD | 1 | 1 | InnovaPrep | SMCSD_InnovaPrep_1_080223 | 14.3 | 14.5 | 0.2 | 200 | 100 | | 40 | |
| 8/2/23 | SMCSD | 1 | 2 | InnovaPrep | SMCSD_InnovaPrep_2_080223 | 14.33 | 14.6 | 0.27 | 200 | 100 | | 29.63 | |
| 8/2/23 | SMCSD | 1 | 3 | InnovaPrep | SMCSD_InnovaPrep_3_080223 | 14.3 | 14.52 | 0.22 | 200 | 100 | | 36.36 | |
| 8/30/23 | EBMUD | 2 | 1 | InnovaPrep | EBMUD_InnovaPrep_1_083023 | 13.4 | 13.63 | 0.23 | 200 | 100 | | 34.78 | |
| 8/30/23 | EBMUD | 2 | 2 | InnovaPrep | EBMUD_InnovaPrep_2_083023 | 13.37 | 13.72 | 0.35 | 200 | 100 | | 22.86 | |
| 8/30/23 | EBMUD | 2 | 3 | InnovaPrep | EBMUD_InnovaPrep_3_083023 | 13.3 | 13.65 | 0.35 | 200 | 100 | | 22.86 | |
| 9/5/23 | WCSD | 2 | 1 | InnovaPrep | WCSD_InnovaPrep_1_090523 | 14.12 | 14.45 | 0.33 | 200 | 100 | | 24.24 | |
| 9/5/23 | WCSD | 2 | 2 | InnovaPrep | WCSD_InnovaPrep_2_090523 | 14.27 | 14.6 | 0.33 | 200 | 100 | | 24.24 | |
| 9/5/23 | WCSD | 2 | 3 | InnovaPrep | WCSD_InnovaPrep_3_090523 | 14.15 | 14.4 | 0.25 | 200 | 100 | | 32 | |
| 9/19/23 | SMCSD | 2 | 1 | InnovaPrep | SMCSD_InnovaPrep_1_091923 | 13.23 | 13.43 | 0.2 | 200 | 100 | | 40 | |
| 9/19/23 | SMCSD | 2 | 2 | InnovaPrep | SMCSD_InnovaPrep_2_091923 | 13.19 | 13.53 | 0.34 | 200 | 100 | | 23.53 | |
| 9/19/23 | SMCSD | 2 | 3 | InnovaPrep | SMCSD_InnovaPrep_3_091923 | 13.13 | 13.3 | 0.17 | 200 | 100 | | 47.06 | |
| 8/30/23 | EBMUD | 2 | 1 | InnovaPrep | EBMUD_InnovaPrep_1_083023_INB | | | 0.23 | 200 | 100 | | 34.78 | |
| 8/30/23 | EBMUD | 2 | 2 | InnovaPrep | EBMUD_InnovaPrep_2_083023_INB | | | 0.35 | 200 | 100 | | 22.86 | |
| 8/30/23 | EBMUD | 2 | 3 | InnovaPrep | EBMUD_InnovaPrep_3_083023_INB | | | 0.35 | 200 | 100 | | 22.86 | |
| 9/5/23 | WCSD | 2 | 1 | InnovaPrep | WCSD_InnovaPrep_1_090523_INB | | | 0.33 | 200 | 100 | | 24.24 | |
| 9/5/23 | WCSD | 2 | 2 | InnovaPrep | WCSD_InnovaPrep_2_090523_INB | | | 0.33 | 200 | 100 | | 24.24 | |
| 9/5/23 | WCSD | 2 | 3 | InnovaPrep | WCSD_InnovaPrep_3_090523_INB | | | 0 | 200 | 100 | | N.A. | sample dropped due to dF |
| 9/19/23 | SMCSD | 2 | 1 | InnovaPrep | SMCSD_InnovaPrep_1_091923_INB | | | 0.2 | 200 | 100 | | 40 | |
| 9/19/23 | SMCSD | 2 | 2 | InnovaPrep | SMCSD_InnovaPrep_2_091923_INB | | | 0.34 | 200 | 100 | | 23.53 | |
| 9/19/23 | SMCSD | 2 | 3 | InnovaPrep | SMCSD_InnovaPrep_3_091923_INB | | | 0.17 | 200 | 100 | | 47.06 | |
| 7/26/23 | EBMUD | 1 | 1 | Nanotrap | EBMUD_Nanotrap_1_072623 | | | 0.2 | 200 | 100 | | 40 | |
| 7/26/23 | EBMUD | 1 | 2 | Nanotrap | EBMUD_Nanotrap_2_072623 | | | 0.2 | 200 | 100 | | 40 | |
| 7/26/23 | EBMUD | 1 | 3 | Nanotrap | EBMUD_Nanotrap_3_072623 | | | 0.2 | 200 | 100 | | 40 | |
| 7/31/23 | WCSD | 1 | 1 | Nanotrap | WCSD_Nanotrap_1_073123 | | | 0.2 | 200 | 100 | | 40 | |
| 7/31/23 | WCSD | 1 | 2 | Nanotrap | WCSD_Nanotrap_2_073123 | | | 0.2 | 200 | 100 | | 40 | |
| 7/31/23 | WCSD | 1 | 3 | Nanotrap | WCSD_Nanotrap_3_073123 | | | 0.2 | 200 | 100 | | 40 | |

| date_sampling | ww_source | timepoint | biological replicate | concentration_methods | sample_id | empty_tube_mass_g | full_tube_mass_g | concentrated_mass_g | extraction_vol_uL_or_mg | elution_vol_uL | cocktail_extr_vol_uL | Effective_vol_mL | notes |
|---|---|---|---|---|---|---|---|---|---|---|---|---|---|
| 8/2/23 | SMCSD | 1 | 1 | Nanotrap | SMCSD_Nanotrap_1_080223 | | | 0.2 | 200 | 100 | | 40 | |
| 8/2/23 | SMCSD | 1 | 2 | Nanotrap | SMCSD_Nanotrap_2_080223 | | | 0.2 | 200 | 100 | | 40 | |
| 8/2/23 | SMCSD | 1 | 3 | Nanotrap | SMCSD_Nanotrap_3_080223 | | | 0.2 | 200 | 100 | | 40 | |
| 8/30/23 | EBMUD | 2 | 1 | Nanotrap | EBMUD_Nanotrap_1_083023 | | | 0.2 | 200 | 100 | | 40 | |
| 8/30/23 | EBMUD | 2 | 2 | Nanotrap | EBMUD_Nanotrap_2_083023 | | | 0.2 | 200 | 100 | | 40 | |
| 8/30/23 | EBMUD | 2 | 3 | Nanotrap | EBMUD_Nanotrap_3_083023 | | | 0.2 | 200 | 100 | | 40 | |
| 9/5/23 | WCSD | 2 | 1 | Nanotrap | WCSD_Nanotrap_1_090523 | | | 0.2 | 200 | 100 | | 40 | |
| 9/5/23 | WCSD | 2 | 2 | Nanotrap | WCSD_Nanotrap_2_090523 | | | 0.2 | 200 | 100 | | 40 | |
| 9/5/23 | WCSD | 2 | 3 | Nanotrap | WCSD_Nanotrap_3_090523 | | | 0.2 | 200 | 100 | | 40 | |
| 9/19/23 | SMCSD | 2 | 1 | Nanotrap | SMCSD_Nanotrap_1_091923 | | | 0.2 | 200 | 100 | | 40 | |
| 9/19/23 | SMCSD | 2 | 2 | Nanotrap | SMCSD_Nanotrap_2_091923 | | | 0.2 | 200 | 100 | | 40 | |
| 9/19/23 | SMCSD | 2 | 3 | Nanotrap | SMCSD_Nanotrap_3_091923 | | | 0.2 | 200 | 100 | | 40 | |
| 8/30/23 | EBMUD | 2 | 1 | Nanotrap | EBMUD_Nanotrap_1_083023_INB | | | 0.2 | 200 | 100 | | 40 | |
| 8/30/23 | EBMUD | 2 | 2 | Nanotrap | EBMUD_Nanotrap_2_083023_INB | | | 0.2 | 200 | 100 | | 40 | |
| 8/30/23 | EBMUD | 2 | 3 | Nanotrap | EBMUD_Nanotrap_3_083023_INB | | | 0.2 | 200 | 100 | | 40 | |
| 9/5/23 | WCSD | 2 | 1 | Nanotrap | WCSD_Nanotrap_1_090523_INB | | | 0.2 | 200 | 100 | | 40 | |
| 9/5/23 | WCSD | 2 | 2 | Nanotrap | WCSD_Nanotrap_2_090523_INB | | | 0.2 | 200 | 100 | | 40 | |
| 9/5/23 | WCSD | 2 | 3 | Nanotrap | WCSD_Nanotrap_3_090523_INB | | | 0.2 | 200 | 100 | | 40 | |
| 9/19/23 | SMCSD | 2 | 1 | Nanotrap | SMCSD_Nanotrap_1_091923_INB | | | 0.2 | 200 | 100 | | 40 | |
| 9/19/23 | SMCSD | 2 | 2 | Nanotrap | SMCSD_Nanotrap_2_091923_INB | | | 0.2 | 200 | 100 | | 40 | |
| 9/19/23 | SMCSD | 2 | 3 | Nanotrap | SMCSD_Nanotrap_3_091923_INB | | | 0.2 | 200 | 100 | | 40 | |
| 7/26/23 | EBMUD | 1 | 1 | Promega | EBMUD_Promega_1_072623 | | | 1 | 1000 | 100 | | 40 | |
| 7/26/23 | EBMUD | 1 | 2 | Promega | EBMUD_Promega_2_072623 | | | 1 | 1000 | 100 | | 40 | |
| 7/26/23 | EBMUD | 1 | 3 | Promega | EBMUD_Promega_3_072623 | | | 1 | 1000 | 100 | | 40 | |
| 7/26/23 | EBMUD | 1 | unspike1 | Promega | EBMUD_Promega_unspike1_072623 | | | 1 | 1000 | 100 | | 40 | |
| 7/26/23 | EBMUD | 1 | unspike2 | Promega | EBMUD_Promega_unspike2_072623 | | | 1 | 1000 | 100 | | 40 | |
| 7/26/23 | EBMUD | 1 | unspike3 | Promega | EBMUD_Promega_unspike3_072623 | | | 1 | 1000 | 100 | | 40 | |
| 7/31/23 | WCSD | 1 | 1 | Promega | WCSD_Promega_1_073123 | | | 1 | 1000 | 100 | | 40 | |
| 7/31/23 | WCSD | 1 | 2 | Promega | WCSD_Promega_2_073123 | | | 1 | 1000 | 100 | | 40 | |
| 7/31/23 | WCSD | 1 | 3 | Promega | WCSD_Promega_3_073123 | | | 1 | 1000 | 100 | | 40 | |
| 7/31/23 | WCSD | 1 | 1 | Promega | WCSD_Promega_1_073123 | | | 1 | 1000 | 100 | | 40 | |
| 7/31/23 | WCSD | 1 | unspike1 | Promega | WCSD_Promega_unspike1_073123 | | | 1 | 1000 | 100 | | 40 | |
| 7/31/23 | WCSD | 1 | unspike2 | Promega | WCSD_Promega_unspike2_073123 | | | 1 | 1000 | 100 | | 40 | |
| 8/2/23 | SMCSD | 1 | 1 | Promega | SMCSD_Promega_1_080223 | | | 1 | 1000 | 100 | | 40 | |
| 8/2/23 | SMCSD | 1 | 2 | Promega | SMCSD_Promega_2_080223 | | | 1 | 1000 | 100 | | 40 | |
| 8/2/23 | SMCSD | 1 | 3 | Promega | SMCSD_Promega_3_080223 | | | 1 | 1000 | 100 | | 40 | |
| 8/2/23 | SMCSD | 1 | unspike1 | Promega | SMCSD_Promega_unspike1_080223 | | | 1 | 1000 | 100 | | 40 | |
| 8/2/23 | SMCSD | 1 | unspike2 | Promega | SMCSD_Promega_unspike2_080223 | | | 1 | 1000 | 100 | | 40 | |
| 8/2/23 | SMCSD | 1 | unspike3 | Promega | SMCSD_Promega_unspike3_080223 | | | 1 | 1000 | 100 | | 40 | |
| 8/30/23 | EBMUD | 2 | 1 | Promega | EBMUD_Promega_1_083023 | | | 1 | 1000 | 100 | | 40 | |
| 8/30/23 | EBMUD | 2 | 2 | Promega | EBMUD_Promega_2_083023 | | | 1 | 1000 | 100 | | 40 | |
| 8/30/23 | EBMUD | 2 | 3 | Promega | EBMUD_Promega_3_083023 | | | 1 | 1000 | 100 | | 40 | |
| 8/30/23 | EBMUD | 2 | unspike1 | Promega | EBMUD_Promega_unspike1_083023 | | | 1 | 1000 | 100 | | 40 | |
| 8/30/23 | EBMUD | 2 | unspike2 | Promega | EBMUD_Promega_unspike2_083023 | | | 1 | 1000 | 100 | | 40 | |
| 8/30/23 | EBMUD | 2 | unspike3 | Promega | EBMUD_Promega_unspike3_083023 | | | 1 | 1000 | 100 | | 40 | |
| 9/5/23 | WCSD | 2 | 1 | Promega | WCSD_Promega_1_090523 | | | 1 | 1000 | 100 | | 40 | |
| 9/5/23 | WCSD | 2 | 2 | Promega | WCSD_Promega_2_090523 | | | 1 | 1000 | 100 | | 40 | |
| 9/5/23 | WCSD | 2 | 3 | Promega | WCSD_Promega_3_090523 | | | 1 | 1000 | 100 | | 40 | |
| 9/5/23 | WCSD | 2 | unspike1 | Promega | WCSD_Promega_unspike1_090523 | | | 1 | 1000 | 100 | | 40 | |
| 9/5/23 | WCSD | 2 | unspike2 | Promega | WCSD_Promega_unspike2_090523 | | | 1 | 1000 | 100 | | 40 | |
| 9/5/23 | WCSD | 2 | unspike3 | Promega | WCSD_Promega_unspike3_090523 | | | 1 | 1000 | 100 | | 40 | |
| 9/19/23 | SMCSD | 2 | 1 | Promega | SMCSD_Promega_1_091923 | | | 1 | 1000 | 100 | | 40 | |
| 9/19/23 | SMCSD | 2 | 2 | Promega | SMCSD_Promega_2_091923 | | | 1 | 1000 | 100 | | 40 | |

| date_sampling | ww_source | timepoint | biological replicate | concentration_methods | sample_id | empty_tube_mass_g | full_tube_mass_g | concentrated_mass_g | extraction_vol_uL_or_mg | elution_vol_uL | cocktail_extr_vol_uL | Effective_vol_mL | notes |
|---|---|---|---|---|---|---|---|---|---|---|---|---|---|
| 9/19/23 | SMCSD | 2 | 3 | Promega | SMCSD_Promega_3_091923 | | | 1 | 1000 | 100 | | 40 | |
| 9/19/23 | SMCSD | 2 | unspike1 | Promega | SMCSD_Promega_unspike1_091923 | | | 1 | 1000 | 100 | | 40 | |
| 9/19/23 | SMCSD | 2 | unspike2 | Promega | SMCSD_Promega_unspike2_091923 | | | 1 | 1000 | 100 | | 40 | |
| 8/30/23 | EBMUD | 2 | 1 | Promega | EBMUD_Promega_1_083023_INB | | | 1 | 1000 | 100 | | 40 | |
| 8/30/23 | EBMUD | 2 | 2 | Promega | EBMUD_Promega_2_083023_INB | | | 1 | 1000 | 100 | | 40 | |
| 8/30/23 | EBMUD | 2 | 3 | Promega | EBMUD_Promega_3_083023_INB | | | 1 | 1000 | 100 | | 40 | |
| 8/30/23 | EBMUD | 2 | unspike1 | Promega | EBMUD_Promega_unspike1_083023_INB | | | 1 | 1000 | 100 | | 40 | |
| 8/30/23 | EBMUD | 2 | unspike2 | Promega | EBMUD_Promega_unspike2_083023_INB | | | 1 | 1000 | 100 | | 40 | |
| 8/30/23 | EBMUD | 2 | unspike3 | Promega | EBMUD_Promega_unspike3_083023_INB | | | 1 | 1000 | 100 | | 40 | |
| 9/5/23 | WCSD | 2 | 1 | Promega | WCSD_Promega_1_090523_INB | | | 1 | 1000 | 100 | | 40 | |
| 9/5/23 | WCSD | 2 | 2 | Promega | WCSD_Promega_2_090523_INB | | | 1 | 1000 | 100 | | 40 | |
| 9/5/23 | WCSD | 2 | 3 | Promega | WCSD_Promega_3_090523_INB | | | 1 | 1000 | 100 | | 40 | |
| 9/5/23 | WCSD | 2 | unspike1 | Promega | WCSD_Promega_unspike1_090523_INB | | | 1 | 1000 | 100 | | 40 | |
| 9/5/23 | WCSD | 2 | unspike2 | Promega | WCSD_Promega_unspike2_090523_INB | | | 1 | 1000 | 100 | | 40 | |
| 9/5/23 | WCSD | 2 | unspike3 | Promega | WCSD_Promega_unspike3_090523_INB | | | 1 | 1000 | 100 | | 40 | |
| 9/19/23 | SMCSD | 2 | 1 | Promega | SMCSD_Promega_1_091923_INB | | | 1 | 1000 | 100 | | 40 | |
| 9/19/23 | SMCSD | 2 | 2 | Promega | SMCSD_Promega_2_091923_INB | | | 1 | 1000 | 100 | | 40 | |
| 9/19/23 | SMCSD | 2 | 3 | Promega | SMCSD_Promega_3_091923_INB | | | 1 | 1000 | 100 | | 40 | |
| 9/19/23 | SMCSD | 2 | unspike1 | Promega | SMCSD_Promega_unspike1_091923_INB | | | 1 | 1000 | 100 | | 40 | |
| 9/19/23 | SMCSD | 2 | unspike2 | Promega | SMCSD_Promega_unspike2_091923_INB | | | 1 | 1000 | 100 | | 40 | |
| 9/19/23 | SMCSD | 2 | unspike3 | Promega | SMCSD_Promega_unspike3_091923_INB | | | 1 | 1000 | 100 | | 40 | |
| 7/26/23 | EBMUD | 1 | 1 | Solids | EBMUD_Solids_1_072623 | 14.14 | 14.77 | 0.63 | 250 | 100 | | 15.87 | |
| 7/26/23 | EBMUD | 1 | 2 | Solids | EBMUD_Solids_2_072623 | 14.07 | 14.65 | 0.58 | 250 | 100 | | 17.24 | |
| 7/26/23 | EBMUD | 1 | 3 | Solids | EBMUD_Solids_3_072623 | 14.23 | 14.83 | 0.6 | 250 | 100 | | 16.67 | |
| 7/31/23 | WCSD | 1 | 1 | Solids | WCSD_Solids_1_073123 | 14.15 | 14.46 | 0.31 | 250 | 100 | | 32.26 | |
| 7/31/23 | WCSD | 1 | 2 | Solids | WCSD_Solids_2_073123 | 14.1 | 14.33 | 0.23 | 250 | 100 | | 43.48 | |
| 7/31/23 | WCSD | 1 | 3 | Solids | WCSD_Solids_3_073123 | 14.15 | 14.43 | 0.28 | 250 | 100 | | 35.71 | |
| 8/2/23 | SMCSD | 1 | 1 | Solids | SMCSD_Solids_1_080223 | 14.03 | 14.34 | 0.31 | 250 | 100 | | 32.26 | |
| 8/2/23 | SMCSD | 1 | 2 | Solids | SMCSD_Solids_2_080223 | 14.15 | 14.8 | 0.65 | 250 | 100 | | 15.38 | |
| 8/2/23 | SMCSD | 1 | 3 | Solids | SMCSD_Solids_3_080223 | 14.15 | 14.97 | 0.82 | 250 | 100 | | 12.2 | |
| 8/30/23 | EBMUD | 2 | 1 | Solids | EBMUD_Solids_1_083023 | 13.33 | 13.84 | 0.51 | 250 | 100 | | 19.61 | |
| 8/30/23 | EBMUD | 2 | 2 | Solids | EBMUD_Solids_2_083023 | 13.41 | 13.78 | 0.37 | 250 | 100 | | 27.03 | |
| 8/30/23 | EBMUD | 2 | 3 | Solids | EBMUD_Solids_3_083023 | 13.43 | 13.81 | 0.38 | 250 | 100 | | 26.32 | |
| 9/5/23 | WCSD | 2 | 1 | Solids | WCSD_Solids_1_090523 | 14.4 | 14.62 | 0.22 | 250 | 100 | | 45.45 | |
| 9/5/23 | WCSD | 2 | 2 | Solids | WCSD_Solids_2_090523 | 14.08 | 14.38 | 0.3 | 250 | 100 | | 33.33 | |
| 9/5/23 | WCSD | 2 | 3 | Solids | WCSD_Solids_3_090523 | 14.22 | 14.5 | 0.28 | 250 | 100 | | 35.71 | |
| 9/19/23 | SMCSD | 2 | 1 | Solids | SMCSD_Solids_1_091923 | 13.02 | 13.34 | 0.32 | 250 | 100 | | 31.25 | |
| 9/19/23 | SMCSD | 2 | 2 | Solids | SMCSD_Solids_2_091923 | 13.17 | 13.46 | 0.29 | 250 | 100 | | 34.48 | |
| 9/19/23 | SMCSD | 2 | 3 | Solids | SMCSD_Solids_3_091923 | 13.23 | 13.49 | 0.26 | 250 | 100 | | 38.46 | |
| 8/30/23 | EBMUD | 2 | 1 | Solids | EBMUD_Solids_1_083023_INB | | | 0.51 | 250 | 100 | | 19.61 | |
| 8/30/23 | EBMUD | 2 | 2 | Solids | EBMUD_Solids_2_083023_INB | | | 0.37 | 250 | 100 | | 27.03 | |
| 8/30/23 | EBMUD | 2 | 3 | Solids | EBMUD_Solids_3_083023_INB | | | 0.38 | 250 | 100 | | 26.32 | |
| 9/5/23 | WCSD | 2 | 1 | Solids | WCSD_Solids_1_090523_INB | | | 0.22 | 250 | 100 | | 45.45 | |
| 9/5/23 | WCSD | 2 | 2 | Solids | WCSD_Solids_2_090523_INB | | | 0.3 | 250 | 100 | | 33.33 | |
| 9/5/23 | WCSD | 2 | 3 | Solids | WCSD_Solids_3_090523_INB | | | 0.28 | 250 | 100 | | 35.71 | |
| 9/19/23 | SMCSD | 2 | 1 | Solids | SMCSD_Solids_1_091923_INB | | | 0.32 | 250 | 100 | | 31.25 | |
| 9/19/23 | SMCSD | 2 | 2 | Solids | SMCSD_Solids_2_091923_INB | | | 0.29 | 250 | 100 | | 34.48 | |
| 9/19/23 | SMCSD | 2 | 3 | Solids | SMCSD_Solids_3_091923_INB | | | 0.26 | 250 | 100 | | 38.46 | |

| well | sample_id | timepoint | ww_source | methods | targets | conc | valid_part | pos_part | neg_part | plate_id | dilution_factor |
|---|---|---|---|---|---|---|---|---|---|---|---|
| B2 | EBMUD_IP_1_072623 | 1 | EBMUD | IP | A6 | 37.01 | 7491 | 88 | 7403 | 2029 | 1 |
| C2 | EBMUD_IP_2_072623 | 1 | EBMUD | IP | A6 | 53.98 | 8102 | 136 | 7966 | 2029 | 1 |
| B2 | EBMUD_IP_1_083023 | 2 | EBMUD | IP | A6 | 24.88 | 8216 | 65 | 8151 | 2034 | 1 |
| C2 | EBMUD_IP_2_083023 | 2 | EBMUD | IP | A6 | 22.98 | 8215 | 59 | 8156 | 2034 | 1 |
| D2 | EBMUD_IP_3_083023 | 2 | EBMUD | IP | A6 | 35.81 | 8122 | 90 | 8032 | 2034 | 1 |
| B4 | EBMUD_IP_1_072623 | 1 | EBMUD | IP | AdV2 | 33.06 | 8019 | 82 | 7937 | 2029 | 1 |
| C4 | EBMUD_IP_2_072623 | 1 | EBMUD | IP | AdV2 | 59.2 | 8288 | 151 | 8137 | 2029 | 1 |
| B4 | EBMUD_IP_1_083023 | 2 | EBMUD | IP | AdV2 | 361.1 | 8230 | 874 | 7356 | 2034 | 1 |
| C4 | EBMUD_IP_2_083023 | 2 | EBMUD | IP | AdV2 | 50.58 | 8277 | 129 | 8148 | 2034 | 1 |
| D4 | EBMUD_IP_3_083023 | 2 | EBMUD | IP | AdV2 | 25.4 | 8267 | 64 | 8203 | 2034 | 1 |
| B4 | EBMUD_IP_1_072623 | 1 | EBMUD | IP | AdV5 | 26.98 | 8019 | 67 | 7952 | 2029 | 1 |
| C4 | EBMUD_IP_2_072623 | 1 | EBMUD | IP | AdV5 | 44.6 | 8288 | 114 | 8174 | 2029 | 1 |
| B4 | EBMUD_IP_1_083023 | 2 | EBMUD | IP | AdV5 | 525.6 | 8224 | 1240 | 6984 | 2034 | 1 |
| C4 | EBMUD_IP_2_083023 | 2 | EBMUD | IP | AdV5 | 53.34 | 8277 | 136 | 8141 | 2034 | 1 |
| D4 | EBMUD_IP_3_083023 | 2 | EBMUD | IP | AdV5 | 34.18 | 8265 | 86 | 8179 | 2034 | 1 |
| B2 | EBMUD_IP_1_072623 | 1 | EBMUD | IP | B5 | 1008.6 | 7446 | 2050 | 5396 | 2029 | 1 |
| C2 | EBMUD_IP_2_072623 | 1 | EBMUD | IP | B5 | 1446.1 | 8085 | 2948 | 5137 | 2029 | 1 |
| B2 | EBMUD_IP_1_083023 | 2 | EBMUD | IP | B5 | 279.7 | 8216 | 702 | 7514 | 2034 | 1 |
| C2 | EBMUD_IP_2_083023 | 2 | EBMUD | IP | B5 | 407.1 | 8209 | 984 | 7225 | 2034 | 1 |
| D2 | EBMUD_IP_3_083023 | 2 | EBMUD | IP | B5 | 431.9 | 8119 | 1021 | 7098 | 2034 | 1 |
| B6 | EBMUD_IP_1_072623 | 1 | EBMUD | IP | H1N1 | 2.079 | 7757 | 5 | 7752 | 2029 | 1 |
| C6 | EBMUD_IP_2_072623 | 1 | EBMUD | IP | H1N1 | 4.138 | 8025 | 10 | 8015 | 2029 | 1 |
| B6 | EBMUD_IP_1_083023 | 2 | EBMUD | IP | H1N1 | 0.793 | 8136 | 2 | 8134 | 2034 | 1 |
| C6 | EBMUD_IP_2_083023 | 2 | EBMUD | IP | H1N1 | 0.401 | 8279 | 1 | 8278 | 2034 | 1 |
| D6 | EBMUD_IP_3_083023 | 2 | EBMUD | IP | H1N1 | 0.797 | 8251 | 2 | 8249 | 2034 | 1 |
| B6 | EBMUD_IP_1_072623 | 1 | EBMUD | IP | H3N2 | 4.992 | 7757 | 12 | 7745 | 2029 | 1 |
| C6 | EBMUD_IP_2_072623 | 1 | EBMUD | IP | H3N2 | 4.967 | 8025 | 12 | 8013 | 2029 | 1 |
| B6 | EBMUD_IP_1_083023 | 2 | EBMUD | IP | H3N2 | 1.189 | 8136 | 3 | 8133 | 2034 | 1 |
| C6 | EBMUD_IP_2_083023 | 2 | EBMUD | IP | H3N2 | 4.011 | 8279 | 10 | 8269 | 2034 | 1 |
| D6 | EBMUD_IP_3_083023 | 2 | EBMUD | IP | H3N2 | 1.594 | 8251 | 4 | 8247 | 2034 | 1 |
| B8 | EBMUD_IP_1_072623 | 1 | EBMUD | IP | N1 | 2.129 | 7676 | 5 | 7671 | 2029 | 1 |
| C8 | EBMUD_IP_2_072623 | 1 | EBMUD | IP | N1 | 0.862 | 7649 | 2 | 7647 | 2029 | 1 |
| B8 | EBMUD_IP_1_083023 | 2 | EBMUD | IP | N1 | 195.8 | 8079 | 470 | 7609 | 2034 | 1 |
| C8 | EBMUD_IP_2_083023 | 2 | EBMUD | IP | N1 | 120.7 | 8231 | 296 | 7935 | 2034 | 1 |
| D8 | EBMUD_IP_3_083023 | 2 | EBMUD | IP | N1 | 145.8 | 8267 | 356 | 7911 | 2034 | 1 |
| B8 | EBMUD_IP_1_072623 | 1 | EBMUD | IP | OC43 | 41.13 | 7673 | 96 | 7577 | 2029 | 1 |
| C8 | EBMUD_IP_2_072623 | 1 | EBMUD | IP | OC43 | 19.87 | 7649 | 46 | 7603 | 2029 | 1 |
| B8 | EBMUD_IP_1_083023 | 2 | EBMUD | IP | OC43 | 646.6 | 8064 | 1448 | 6616 | 2034 | 1 |
| C8 | EBMUD_IP_2_083023 | 2 | EBMUD | IP | OC43 | 755.3 | 8190 | 1678 | 6512 | 2034 | 1 |
| D8 | EBMUD_IP_3_083023 | 2 | EBMUD | IP | OC43 | 955.5 | 8262 | 2071 | 6191 | 2034 | 1 |
| D1 | EBMUD_NT_1_072623 | 1 | EBMUD | NT | A6 | 737.3 | 8204 | 1727 | 6477 | 2029 | 1 |
| E1 | EBMUD_NT_2_072623 | 1 | EBMUD | NT | A6 | 598.9 | 8146 | 1406 | 6740 | 2029 | 1 |
| F1 | EBMUD_NT_3_072623 | 1 | EBMUD | NT | A6 | 730.6 | 8169 | 1679 | 6490 | 2029 | 1 |
| D1 | EBMUD_NT_1_083023 | 2 | EBMUD | NT | A6 | 133.5 | 7971 | 334 | 7637 | 2034 | 1 |

| | | | | | | | | | | | |
|---|---|---|---|---|---|---|---|---|---|---|---|
| E1 | EBMUD_NT_2_083023 | 2 | EBMUD | NT | A6 | 20.25 | 7987 | 51 | 7936 | 2034 | 1 |
| F1 | EBMUD_NT_3_083023 | 2 | EBMUD | NT | A6 | 35.58 | 7719 | 86 | 7633 | 2034 | 1 |
| D3 | EBMUD_NT_1_072623 | 1 | EBMUD | NT | AdV2 | 132 | 8285 | 330 | 7955 | 2029 | 1 |
| E3 | EBMUD_NT_2_072623 | 1 | EBMUD | NT | AdV2 | 113.4 | 8237 | 283 | 7954 | 2029 | 1 |
| F3 | EBMUD_NT_3_072623 | 1 | EBMUD | NT | AdV2 | 200.2 | 8247 | 491 | 7756 | 2029 | 1 |
| D3 | EBMUD_NT_1_083023 | 2 | EBMUD | NT | AdV2 | 362.1 | 8265 | 872 | 7393 | 2034 | 1 |
| E3 | EBMUD_NT_2_083023 | 2 | EBMUD | NT | AdV2 | 283.7 | 8285 | 694 | 7591 | 2034 | 1 |
| F3 | EBMUD_NT_3_083023 | 2 | EBMUD | NT | AdV2 | 277.1 | 8262 | 673 | 7589 | 2034 | 1 |
| D3 | EBMUD_NT_1_072623 | 1 | EBMUD | NT | AdV5 | 150.9 | 8285 | 376 | 7909 | 2029 | 1 |
| E3 | EBMUD_NT_2_072623 | 1 | EBMUD | NT | AdV5 | 120.3 | 8237 | 300 | 7937 | 2029 | 1 |
| F3 | EBMUD_NT_3_072623 | 1 | EBMUD | NT | AdV5 | 192.2 | 8250 | 472 | 7778 | 2029 | 1 |
| D3 | EBMUD_NT_1_083023 | 2 | EBMUD | NT | AdV5 | 320.7 | 8264 | 777 | 7487 | 2034 | 1 |
| E3 | EBMUD_NT_2_083023 | 2 | EBMUD | NT | AdV5 | 283 | 8282 | 692 | 7590 | 2034 | 1 |
| F3 | EBMUD_NT_3_083023 | 2 | EBMUD | NT | AdV5 | 287 | 8262 | 696 | 7566 | 2034 | 1 |
| D1 | EBMUD_NT_1_072623 | 1 | EBMUD | NT | B5 | 64.51 | 8257 | 169 | 8088 | 2029 | 1 |
| E1 | EBMUD_NT_2_072623 | 1 | EBMUD | NT | B5 | 59.91 | 8150 | 153 | 7997 | 2029 | 1 |
| F1 | EBMUD_NT_3_072623 | 1 | EBMUD | NT | B5 | 68.12 | 8198 | 174 | 8024 | 2029 | 1 |
| D1 | EBMUD_NT_1_083023 | 2 | EBMUD | NT | B5 | 15.29 | 7975 | 39 | 7936 | 2034 | 1 |
| E1 | EBMUD_NT_2_083023 | 2 | EBMUD | NT | B5 | 4.357 | 7987 | 11 | 7976 | 2034 | 1 |
| F1 | EBMUD_NT_3_083023 | 2 | EBMUD | NT | B5 | 2.058 | 7719 | 5 | 7714 | 2034 | 1 |
| D5 | EBMUD_NT_1_083023 | 1 | EBMUD | NT | H1N1 | 1.189 | 8245 | 3 | 8242 | 2034 | 1 |
| E5 | EBMUD_NT_2_083023 | 1 | EBMUD | NT | H1N1 | 0 | 8273 | 0 | 8273 | 2034 | 1 |
| F5 | EBMUD_NT_3_083023 | 1 | EBMUD | NT | H1N1 | 0.813 | 8272 | 2 | 8270 | 2034 | 1 |
| D5 | EBMUD_NT_1_072623 | 2 | EBMUD | NT | H1N1 | 13.33 | 8106 | 33 | 8073 | 2029 | 1 |
| E5 | EBMUD_NT_2_072623 | 2 | EBMUD | NT | H1N1 | 4.902 | 8207 | 12 | 8195 | 2029 | 1 |
| F5 | EBMUD_NT_3_072623 | 2 | EBMUD | NT | H1N1 | 9.425 | 8213 | 23 | 8190 | 2029 | 1 |
| D5 | EBMUD_NT_1_083023 | 1 | EBMUD | NT | H3N2 | 0.793 | 8245 | 2 | 8243 | 2034 | 1 |
| E5 | EBMUD_NT_2_083023 | 1 | EBMUD | NT | H3N2 | 0 | 8273 | 0 | 8273 | 2034 | 1 |
| F5 | EBMUD_NT_3_083023 | 1 | EBMUD | NT | H3N2 | 0 | 8272 | 0 | 8272 | 2034 | 1 |
| D5 | EBMUD_NT_1_072623 | 2 | EBMUD | NT | H3N2 | 20.22 | 8106 | 50 | 8056 | 2029 | 1 |
| E5 | EBMUD_NT_2_072623 | 2 | EBMUD | NT | H3N2 | 15.14 | 8208 | 37 | 8171 | 2029 | 1 |
| F5 | EBMUD_NT_3_072623 | 2 | EBMUD | NT | H3N2 | 17.64 | 8213 | 43 | 8170 | 2029 | 1 |
| D7 | EBMUD_NT_1_083023 | 1 | EBMUD | NT | N1 | 758 | 8247 | 1675 | 6572 | 2034 | 1 |
| E7 | EBMUD_NT_2_083023 | 1 | EBMUD | NT | N1 | 211.7 | 8209 | 499 | 7710 | 2034 | 1 |
| F7 | EBMUD_NT_3_083023 | 1 | EBMUD | NT | N1 | 235.4 | 8286 | 563 | 7723 | 2034 | 1 |
| D7 | EBMUD_NT_1_072623 | 2 | EBMUD | NT | N1 | 1772.4 | 7419 | 3056 | 4363 | 2029 | 1 |
| E7 | EBMUD_NT_2_072623 | 2 | EBMUD | NT | N1 | 1431.5 | 8113 | 2804 | 5309 | 2029 | 1 |
| F7 | EBMUD_NT_3_072623 | 2 | EBMUD | NT | N1 | 1483.6 | 7685 | 2753 | 4932 | 2029 | 1 |
| D7 | EBMUD_NT_1_083023 | 1 | EBMUD | NT | OC43 | 338 | 8288 | 798 | 7490 | 2034 | 1 |
| E7 | EBMUD_NT_2_083023 | 1 | EBMUD | NT | OC43 | 88.24 | 8217 | 212 | 8005 | 2034 | 1 |
| F7 | EBMUD_NT_3_083023 | 1 | EBMUD | NT | OC43 | 110.4 | 8286 | 269 | 8017 | 2034 | 1 |
| D7 | EBMUD_NT_1_072623 | 2 | EBMUD | NT | OC43 | 1586.6 | 7339 | 2776 | 4563 | 2029 | 1 |
| E7 | EBMUD_NT_2_072623 | 2 | EBMUD | NT | OC43 | 1212.5 | 8149 | 2459 | 5690 | 2029 | 1 |
| F7 | EBMUD_NT_3_072623 | 2 | EBMUD | NT | OC43 | 1335.9 | 7669 | 2525 | 5144 | 2029 | 1 |
| A1 | EBMUD_PMG_1_072623 | 1 | EBMUD | PMG | A6 | 151.3 | 8273 | 401 | 7872 | 2029 | 1 |

| | | | | | | | | | | | |
|---|---|---|---|---|---|---|---|---|---|---|---|
| B1 | EBMUD_PMG_2_072623 | 1 | EBMUD | PMG | A6 | 77.5 | 7420 | 182 | 7238 | 2029 | 1 |
| C1 | EBMUD_PMG_3_072623 | 1 | EBMUD | PMG | A6 | 97.69 | 7455 | 230 | 7225 | 2029 | 1 |
| A1 | EBMUD_PMG_1_083023 | 2 | EBMUD | PMG | A6 | 88.5 | 8241 | 236 | 8005 | 2034 | 1 |
| B1 | EBMUD_PMG_2_083023 | 2 | EBMUD | PMG | A6 | 74.52 | 8264 | 195 | 8069 | 2034 | 1 |
| C1 | EBMUD_PMG_3_083023 | 2 | EBMUD | PMG | A6 | 71.37 | 7643 | 173 | 7470 | 2034 | 1 |
| A3 | EBMUD_PMG_1_072623 | 1 | EBMUD | PMG | AdV2 | 254.1 | 2426 | 190 | 2236 | 2029 | 1 |
| B3 | EBMUD_PMG_2_072623 | 1 | EBMUD | PMG | AdV2 | 0 | 8087 | 0 | 8087 | 2029 | 1 |
| C3 | EBMUD_PMG_3_072623 | 1 | EBMUD | PMG | AdV2 | 388.6 | 1412 | 159 | 1253 | 2029 | 1 |
| A3 | EBMUD_PMG_1_083023 | 2 | EBMUD | PMG | AdV2 | 1412.2 | 8098 | 2951 | 5147 | 2034 | 1 |
| B3 | EBMUD_PMG_2_083023 | 2 | EBMUD | PMG | AdV2 | 1208.1 | 8123 | 2558 | 5565 | 2034 | 1 |
| C3 | EBMUD_PMG_3_083023 | 2 | EBMUD | PMG | AdV2 | 1024.3 | 8226 | 2222 | 6004 | 2034 | 1 |
| A3 | EBMUD_PMG_1_072623 | 1 | EBMUD | PMG | AdV5 | 2635.4 | 7984 | 4557 | 3427 | 2029 | 1 |
| B3 | EBMUD_PMG_2_072623 | 1 | EBMUD | PMG | AdV5 | 1812.3 | 843 | 394 | 449 | 2042 | 1 |
| C3 | EBMUD_PMG_3_072623 | 1 | EBMUD | PMG | AdV5 | 1591.8 | 8148 | 3153 | 4995 | 2029 | 1 |
| A3 | EBMUD_PMG_1_083023 | 2 | EBMUD | PMG | AdV5 | 1135.3 | 8086 | 2469 | 5617 | 2034 | 1 |
| B3 | EBMUD_PMG_2_083023 | 2 | EBMUD | PMG | AdV5 | 1097 | 8175 | 2376 | 5799 | 2034 | 1 |
| C3 | EBMUD_PMG_3_083023 | 2 | EBMUD | PMG | AdV5 | 852 | 8211 | 1892 | 6319 | 2034 | 1 |
| A1 | EBMUD_PMG_1_072623 | 1 | EBMUD | PMG | B5 | 1637.2 | 8273 | 3440 | 4833 | 2029 | 1 |
| B1 | EBMUD_PMG_2_072623 | 1 | EBMUD | PMG | B5 | 741.9 | 7382 | 1562 | 5820 | 2029 | 1 |
| C1 | EBMUD_PMG_3_072623 | 1 | EBMUD | PMG | B5 | 1004.2 | 7411 | 2041 | 5370 | 2029 | 1 |
| A1 | EBMUD_PMG_1_083023 | 2 | EBMUD | PMG | B5 | 612.5 | 8207 | 1495 | 6712 | 2034 | 1 |
| B1 | EBMUD_PMG_2_083023 | 2 | EBMUD | PMG | B5 | 432.2 | 8257 | 1068 | 7189 | 2034 | 1 |
| C1 | EBMUD_PMG_3_083023 | 2 | EBMUD | PMG | B5 | 444.7 | 7634 | 1015 | 6619 | 2034 | 1 |
| A5 | EBMUD_PMG_1_072623 | 1 | EBMUD | PMG | H1N1 | 40.98 | 8049 | 105 | 7944 | 2029 | 1 |
| B5 | EBMUD_PMG_2_072623 | 1 | EBMUD | PMG | H1N1 | 22.69 | 7719 | 54 | 7665 | 2029 | 1 |
| C5 | EBMUD_PMG_3_072623 | 1 | EBMUD | PMG | H1N1 | 24.71 | 7431 | 56 | 7375 | 2029 | 1 |
| A5 | EBMUD_PMG_1_083023 | 2 | EBMUD | PMG | H1N1 | 6.054 | 8256 | 16 | 8240 | 2034 | 1 |
| B5 | EBMUD_PMG_2_083023 | 2 | EBMUD | PMG | H1N1 | 6.357 | 8143 | 16 | 8127 | 2034 | 1 |
| C5 | EBMUD_PMG_3_083023 | 2 | EBMUD | PMG | H1N1 | 5.932 | 8268 | 15 | 8253 | 2034 | 1 |
| A5 | EBMUD_PMG_1_072623 | 1 | EBMUD | PMG | H3N2 | 94.86 | 8049 | 241 | 7808 | 2029 | 1 |
| B5 | EBMUD_PMG_2_072623 | 1 | EBMUD | PMG | H3N2 | 41.72 | 7719 | 99 | 7620 | 2029 | 1 |
| C5 | EBMUD_PMG_3_072623 | 1 | EBMUD | PMG | H3N2 | 64.82 | 7431 | 146 | 7285 | 2029 | 1 |
| A5 | EBMUD_PMG_1_083023 | 2 | EBMUD | PMG | H3N2 | 12.88 | 8256 | 34 | 8222 | 2034 | 1 |
| B5 | EBMUD_PMG_2_083023 | 2 | EBMUD | PMG | H3N2 | 10.73 | 8143 | 27 | 8116 | 2034 | 1 |
| C5 | EBMUD_PMG_3_083023 | 2 | EBMUD | PMG | H3N2 | 12.67 | 8267 | 32 | 8235 | 2034 | 1 |
| A7 | EBMUD_PMG_1_072623 | 1 | EBMUD | PMG | N1 | 4853.1 | 7440 | 5830 | 1610 | 2029 | 1 |
| B7 | EBMUD_PMG_2_072623 | 1 | EBMUD | PMG | N1 | 4744.4 | 8022 | 6186 | 1836 | 2029 | 1 |
| C7 | EBMUD_PMG_3_072623 | 1 | EBMUD | PMG | N1 | 2605.5 | 7516 | 4115 | 3401 | 2029 | 1 |
| A7 | EBMUD_PMG_1_083023 | 2 | EBMUD | PMG | N1 | 2331.9 | 8275 | 4309 | 3966 | 2034 | 1 |
| B7 | EBMUD_PMG_2_083023 | 2 | EBMUD | PMG | N1 | 1977.9 | 8229 | 3779 | 4450 | 2034 | 1 |
| C7 | EBMUD_PMG_3_083023 | 2 | EBMUD | PMG | N1 | 1890.3 | 8133 | 3558 | 4575 | 2034 | 1 |
| A7 | EBMUD_PMG_1_072623 | 1 | EBMUD | PMG | OC43 | 2003.5 | 7440 | 3485 | 3955 | 2029 | 1 |
| B7 | EBMUD_PMG_2_072623 | 1 | EBMUD | PMG | OC43 | 714.3 | 7971 | 1587 | 6384 | 2029 | 1 |
| C7 | EBMUD_PMG_3_072623 | 1 | EBMUD | PMG | OC43 | 1880.2 | 7516 | 3275 | 4241 | 2029 | 1 |
| A7 | EBMUD_PMG_1_083023 | 2 | EBMUD | PMG | OC43 | 1644 | 8275 | 3348 | 4927 | 2034 | 1 |

| | | | | | | | | | | | |
|---|---|---|---|---|---|---|---|---|---|---|---|
| B7 | EBMUD_PMG_2_083023 | 2 | EBMUD | PMG | OC43 | 1468.5 | 8143 | 2984 | 5159 | 2034 | 1 |
| C7 | EBMUD_PMG_3_083023 | 2 | EBMUD | PMG | OC43 | 1526.9 | 8077 | 3002 | 5075 | 2034 | 1 |
| G1 | EBMUD_S_1_072623 | 1 | EBMUD | S | A6 | 254.9 | 8223 | 642 | 7581 | 2029 | 1 |
| H1 | EBMUD_S_2_072623 | 1 | EBMUD | S | A6 | 204.9 | 8081 | 524 | 7557 | 2029 | 1 |
| A2 | EBMUD_S_3_072623 | 1 | EBMUD | S | A6 | 180.9 | 7728 | 437 | 7291 | 2029 | 1 |
| G1 | EBMUD_S_1_083023 | 2 | EBMUD | S | A6 | 87.96 | 7989 | 221 | 7768 | 2034 | 1 |
| H1 | EBMUD_S_2_083023 | 2 | EBMUD | S | A6 | 89.45 | 8183 | 236 | 7947 | 2034 | 1 |
| A2 | EBMUD_S_3_083023 | 2 | EBMUD | S | A6 | 150.7 | 8172 | 387 | 7785 | 2034 | 1 |
| G3 | EBMUD_S_1_072623 | 1 | EBMUD | S | AdV2 | 53.39 | 8176 | 134 | 8042 | 2029 | 1 |
| H3 | EBMUD_S_2_072623 | 1 | EBMUD | S | AdV2 | 42.95 | 8248 | 112 | 8136 | 2029 | 1 |
| A4 | EBMUD_S_3_072623 | 1 | EBMUD | S | AdV2 | 25.8 | 8143 | 67 | 8076 | 2029 | 1 |
| G3 | EBMUD_S_1_083023 | 2 | EBMUD | S | AdV2 | 111.7 | 8150 | 277 | 7873 | 2034 | 1 |
| H3 | EBMUD_S_2_083023 | 2 | EBMUD | S | AdV2 | 96.17 | 8259 | 249 | 8010 | 2034 | 1 |
| A4 | EBMUD_S_3_083023 | 2 | EBMUD | S | AdV2 | 91.27 | 8263 | 238 | 8025 | 2034 | 1 |
| G3 | EBMUD_S_1_072623 | 1 | EBMUD | S | AdV5 | 111.9 | 7725 | 263 | 7462 | 2029 | 1 |
| H3 | EBMUD_S_2_072623 | 1 | EBMUD | S | AdV5 | 79.89 | 8243 | 207 | 8036 | 2029 | 1 |
| A4 | EBMUD_S_3_072623 | 1 | EBMUD | S | AdV5 | 28.12 | 8143 | 73 | 8070 | 2029 | 1 |
| G3 | EBMUD_S_1_083023 | 2 | EBMUD | S | AdV5 | 97.39 | 8150 | 242 | 7908 | 2034 | 1 |
| H3 | EBMUD_S_2_083023 | 2 | EBMUD | S | AdV5 | 73.89 | 8259 | 192 | 8067 | 2034 | 1 |
| A4 | EBMUD_S_3_083023 | 2 | EBMUD | S | AdV5 | 75.36 | 8263 | 197 | 8066 | 2034 | 1 |
| G1 | EBMUD_S_1_072623 | 1 | EBMUD | S | B5 | 34.51 | 8223 | 90 | 8133 | 2029 | 1 |
| H1 | EBMUD_S_2_072623 | 1 | EBMUD | S | B5 | 33.84 | 8084 | 89 | 7995 | 2029 | 1 |
| A2 | EBMUD_S_3_072623 | 1 | EBMUD | S | B5 | 25.03 | 7728 | 62 | 7666 | 2029 | 1 |
| G1 | EBMUD_S_1_083023 | 2 | EBMUD | S | B5 | 16.92 | 7989 | 43 | 7946 | 2034 | 1 |
| H1 | EBMUD_S_2_083023 | 2 | EBMUD | S | B5 | 21.37 | 8183 | 57 | 8126 | 2034 | 1 |
| A2 | EBMUD_S_3_083023 | 2 | EBMUD | S | B5 | 47.12 | 8172 | 123 | 8049 | 2034 | 1 |
| G5 | EBMUD_S_1_072623 | 1 | EBMUD | S | H1N1 | 0.397 | 8260 | 1 | 8259 | 2029 | 1 |
| H5 | EBMUD_S_2_072623 | 1 | EBMUD | S | H1N1 | 1.532 | 8181 | 4 | 8177 | 2029 | 1 |
| A6 | EBMUD_S_3_072623 | 1 | EBMUD | S | H1N1 | 0 | 7964 | 0 | 7964 | 2029 | 1 |
| G5 | EBMUD_S_1_083023 | 2 | EBMUD | S | H1N1 | 0 | 8191 | 0 | 8191 | 2034 | 1 |
| H5 | EBMUD_S_2_083023 | 2 | EBMUD | S | H1N1 | 0 | 8067 | 0 | 8067 | 2034 | 1 |
| A6 | EBMUD_S_3_083023 | 2 | EBMUD | S | H1N1 | 0 | 8095 | 0 | 8095 | 2034 | 1 |
| G5 | EBMUD_S_1_072623 | 1 | EBMUD | S | H3N2 | 0.397 | 8260 | 1 | 8259 | 2029 | 1 |
| H5 | EBMUD_S_2_072623 | 1 | EBMUD | S | H3N2 | 0.766 | 8181 | 2 | 8179 | 2029 | 1 |
| A6 | EBMUD_S_3_072623 | 1 | EBMUD | S | H3N2 | 0 | 7964 | 0 | 7964 | 2029 | 1 |
| G5 | EBMUD_S_1_083023 | 2 | EBMUD | S | H3N2 | 0 | 8191 | 0 | 8191 | 2034 | 1 |
| H5 | EBMUD_S_2_083023 | 2 | EBMUD | S | H3N2 | 0 | 8067 | 0 | 8067 | 2034 | 1 |
| A6 | EBMUD_S_3_083023 | 2 | EBMUD | S | H3N2 | 0 | 8095 | 0 | 8095 | 2034 | 1 |
| G7 | EBMUD_S_1_072623 | 1 | EBMUD | S | N1 | 12.72 | 7562 | 29 | 7533 | 2029 | 1 |
| H7 | EBMUD_S_2_072623 | 1 | EBMUD | S | N1 | 8.52 | 8108 | 22 | 8086 | 2029 | 1 |
| A8 | EBMUD_S_3_072623 | 1 | EBMUD | S | N1 | 3.265 | 7599 | 8 | 7591 | 2029 | 1 |
| G7 | EBMUD_S_1_083023 | 2 | EBMUD | S | N1 | 10.51 | 8204 | 26 | 8178 | 2034 | 1 |
| H7 | EBMUD_S_2_083023 | 2 | EBMUD | S | N1 | 10.29 | 8240 | 27 | 8213 | 2034 | 1 |
| A8 | EBMUD_S_3_083023 | 2 | EBMUD | S | N1 | 16.38 | 8162 | 43 | 8119 | 2034 | 1 |
| G7 | EBMUD_S_1_072623 | 1 | EBMUD | S | OC43 | 46.74 | 7562 | 106 | 7456 | 2029 | 1 |

| | | | | | | | | | | | |
|---|---|---|---|---|---|---|---|---|---|---|---|
| H7 | EBMUD_S_2_072623 | 1 | EBMUD | S | OC43 | 50.29 | 8108 | 129 | 7979 | 2029 | 1 |
| A8 | EBMUD_S_3_072623 | 1 | EBMUD | S | OC43 | 22.93 | 7599 | 56 | 7543 | 2029 | 1 |
| G7 | EBMUD_S_1_083023 | 2 | EBMUD | S | OC43 | 6.869 | 8204 | 17 | 8187 | 2034 | 1 |
| H7 | EBMUD_S_2_083023 | 2 | EBMUD | S | OC43 | 9.527 | 8241 | 25 | 8216 | 2034 | 1 |
| A8 | EBMUD_S_3_083023 | 2 | EBMUD | S | OC43 | 9.891 | 8162 | 26 | 8136 | 2034 | 1 |
| B2 | SMCSD_IP_1_080223 | 1 | SMCSD | IP | A6 | 106.3 | 8155 | 272 | 7883 | 2027 | 1 |
| C2 | SMCSD_IP_2_080223 | 1 | SMCSD | IP | A6 | 56.72 | 8281 | 146 | 8135 | 2027 | 1 |
| D2 | SMCSD_IP_3_080223 | 1 | SMCSD | IP | A6 | 90.51 | 8210 | 228 | 7982 | 2027 | 1 |
| B2 | SMCSD_IP_1_091923 | 2 | SMCSD | IP | A6 | 51.7 | 8186 | 134 | 8052 | 2038 | 1 |
| C2 | SMCSD_IP_2_091923 | 2 | SMCSD | IP | A6 | 40.58 | 8224 | 104 | 8120 | 2038 | 1 |
| D2 | SMCSD_IP_3_091923 | 2 | SMCSD | IP | A6 | 47.19 | 8233 | 120 | 8113 | 2038 | 1 |
| B4 | SMCSD_IP_1_080223 | 1 | SMCSD | IP | AdV2 | 226.9 | 8252 | 562 | 7690 | 2027 | 1 |
| C4 | SMCSD_IP_2_080223 | 1 | SMCSD | IP | AdV2 | 358 | 8278 | 871 | 7407 | 2027 | 1 |
| D4 | SMCSD_IP_3_080223 | 1 | SMCSD | IP | AdV2 | 283.5 | 8267 | 687 | 7580 | 2027 | 1 |
| B4 | SMCSD_IP_1_091923 | 2 | SMCSD | IP | AdV2 | 815.3 | 8168 | 1829 | 6339 | 2038 | 1 |
| C4 | SMCSD_IP_2_091923 | 2 | SMCSD | IP | AdV2 | 754.3 | 8216 | 1716 | 6500 | 2038 | 1 |
| D4 | SMCSD_IP_3_091923 | 2 | SMCSD | IP | AdV2 | 846.8 | 8205 | 1873 | 6332 | 2038 | 1 |
| B4 | SMCSD_IP_1_080223 | 1 | SMCSD | IP | AdV5 | 329.9 | 8246 | 804 | 7442 | 2027 | 1 |
| C4 | SMCSD_IP_2_080223 | 1 | SMCSD | IP | AdV5 | 521.2 | 8271 | 1236 | 7035 | 2027 | 1 |
| D4 | SMCSD_IP_3_080223 | 1 | SMCSD | IP | AdV5 | 363.6 | 8273 | 871 | 7402 | 2027 | 1 |
| B4 | SMCSD_IP_1_091923 | 2 | SMCSD | IP | AdV5 | 668.1 | 8210 | 1540 | 6670 | 2038 | 1 |
| C4 | SMCSD_IP_2_091923 | 2 | SMCSD | IP | AdV5 | 684.7 | 8243 | 1579 | 6664 | 2038 | 1 |
| D4 | SMCSD_IP_3_091923 | 2 | SMCSD | IP | AdV5 | 732.3 | 8234 | 1653 | 6581 | 2038 | 1 |
| B2 | SMCSD_IP_1_080223 | 1 | SMCSD | IP | B5 | 871.7 | 8072 | 1961 | 6111 | 2027 | 1 |
| C2 | SMCSD_IP_2_080223 | 1 | SMCSD | IP | B5 | 532.4 | 8272 | 1272 | 7000 | 2027 | 1 |
| D2 | SMCSD_IP_3_080223 | 1 | SMCSD | IP | B5 | 932.9 | 8192 | 2064 | 6128 | 2027 | 1 |
| B2 | SMCSD_IP_1_091923 | 2 | SMCSD | IP | B5 | 321.8 | 8183 | 799 | 7384 | 2038 | 1 |
| C2 | SMCSD_IP_2_091923 | 2 | SMCSD | IP | B5 | 255.1 | 8221 | 632 | 7589 | 2038 | 1 |
| D2 | SMCSD_IP_3_091923 | 2 | SMCSD | IP | B5 | 431.2 | 8227 | 1033 | 7194 | 2038 | 1 |
| B6 | SMCSD_IP_1_080223 | 1 | SMCSD | IP | H1N1 | 0 | 8267 | 0 | 8267 | 2027 | 1 |
| C6 | SMCSD_IP_2_080223 | 1 | SMCSD | IP | H1N1 | 1.608 | 8260 | 4 | 8256 | 2027 | 1 |
| D6 | SMCSD_IP_3_080223 | 1 | SMCSD | IP | H1N1 | 3.583 | 8260 | 9 | 8251 | 2027 | 1 |
| B6 | SMCSD_IP_1_091923 | 2 | SMCSD | IP | H1N1 | 0 | 8254 | 0 | 8254 | 2038 | 1 |
| C6 | SMCSD_IP_2_091923 | 2 | SMCSD | IP | H1N1 | 0 | 8260 | 0 | 8260 | 2038 | 1 |
| D6 | SMCSD_IP_3_091923 | 2 | SMCSD | IP | H1N1 | 0 | 8265 | 0 | 8265 | 2038 | 1 |
| B6 | SMCSD_IP_1_080223 | 1 | SMCSD | IP | H3N2 | 13.29 | 8267 | 34 | 8233 | 2027 | 1 |
| C6 | SMCSD_IP_2_080223 | 1 | SMCSD | IP | H3N2 | 10.46 | 8260 | 26 | 8234 | 2027 | 1 |
| D6 | SMCSD_IP_3_080223 | 1 | SMCSD | IP | H3N2 | 11.16 | 8259 | 28 | 8231 | 2027 | 1 |
| B6 | SMCSD_IP_1_091923 | 2 | SMCSD | IP | H3N2 | 0.781 | 8254 | 2 | 8252 | 2038 | 1 |
| C6 | SMCSD_IP_2_091923 | 2 | SMCSD | IP | H3N2 | 1.608 | 8260 | 4 | 8256 | 2038 | 1 |
| D6 | SMCSD_IP_3_091923 | 2 | SMCSD | IP | H3N2 | 1.989 | 8265 | 5 | 8260 | 2038 | 1 |
| B8 | SMCSD_IP_1_080223 | 1 | SMCSD | IP | N1 | 93.99 | 8215 | 233 | 7982 | 2027 | 1 |
| C8 | SMCSD_IP_2_080223 | 1 | SMCSD | IP | N1 | 94.2 | 8266 | 233 | 8033 | 2027 | 1 |
| D8 | SMCSD_IP_3_080223 | 1 | SMCSD | IP | N1 | 77.71 | 8278 | 192 | 8086 | 2027 | 1 |
| B8 | SMCSD_IP_1_091923 | 2 | SMCSD | IP | N1 | 115 | 8183 | 283 | 7900 | 2038 | 1 |

| | | | | | | | | | | | |
|---|---|---|---|---|---|---|---|---|---|---|---|
| C8 | SMCSD_IP_2_091923 | 2 | SMCSD | IP | N1 | 79.91 | 8221 | 197 | 8024 | 2038 | 1 |
| D8 | SMCSD_IP_3_091923 | 2 | SMCSD | IP | N1 | 117.4 | 8208 | 286 | 7922 | 2038 | 1 |
| B8 | SMCSD_IP_1_080223 | 1 | SMCSD | IP | OC43 | 514.8 | 8210 | 1197 | 7013 | 2027 | 1 |
| C8 | SMCSD_IP_2_080223 | 1 | SMCSD | IP | OC43 | 689.7 | 8243 | 1557 | 6686 | 2027 | 1 |
| D8 | SMCSD_IP_3_080223 | 1 | SMCSD | IP | OC43 | 489.2 | 8272 | 1136 | 7136 | 2027 | 1 |
| B8 | SMCSD_IP_1_091923 | 2 | SMCSD | IP | OC43 | 159.2 | 8177 | 389 | 7788 | 2038 | 1 |
| C8 | SMCSD_IP_2_091923 | 2 | SMCSD | IP | OC43 | 59.04 | 8221 | 146 | 8075 | 2038 | 1 |
| D8 | SMCSD_IP_3_091923 | 2 | SMCSD | IP | OC43 | 198.7 | 8208 | 478 | 7730 | 2038 | 1 |
| D1 | SMCSD_NT_1_080223 | 1 | SMCSD | NT | A6 | 277 | 8121 | 690 | 7431 | 2027 | 1 |
| E1 | SMCSD_NT_2_080223 | 1 | SMCSD | NT | A6 | 313.8 | 8276 | 782 | 7494 | 2027 | 1 |
| F1 | SMCSD_NT_3_080223 | 1 | SMCSD | NT | A6 | 161.4 | 8255 | 409 | 7846 | 2027 | 1 |
| D1 | SMCSD_NT_1_091923 | 2 | SMCSD | NT | A6 | 98.19 | 8165 | 253 | 7912 | 2038 | 1 |
| E1 | SMCSD_NT_2_091923 | 2 | SMCSD | NT | A6 | 156.4 | 8288 | 400 | 7888 | 2038 | 1 |
| F1 | SMCSD_NT_3_091923 | 2 | SMCSD | NT | A6 | 151.7 | 8208 | 383 | 7825 | 2038 | 1 |
| D3 | SMCSD_NT_1_080223 | 1 | SMCSD | NT | AdV2 | 477.3 | 8195 | 1120 | 7075 | 2027 | 1 |
| E3 | SMCSD_NT_2_080223 | 1 | SMCSD | NT | AdV2 | 430.1 | 8237 | 1023 | 7214 | 2027 | 1 |
| F3 | SMCSD_NT_3_080223 | 1 | SMCSD | NT | AdV2 | 338.5 | 8269 | 815 | 7454 | 2027 | 1 |
| D3 | SMCSD_NT_1_091923 | 2 | SMCSD | NT | AdV2 | 268.4 | 8284 | 657 | 7627 | 2038 | 1 |
| E3 | SMCSD_NT_2_091923 | 2 | SMCSD | NT | AdV2 | 454.5 | 8267 | 1081 | 7186 | 2038 | 1 |
| F3 | SMCSD_NT_3_091923 | 2 | SMCSD | NT | AdV2 | 310.5 | 8248 | 749 | 7499 | 2038 | 1 |
| D3 | SMCSD_NT_1_080223 | 1 | SMCSD | NT | AdV5 | 206.3 | 8205 | 505 | 7700 | 2027 | 1 |
| E3 | SMCSD_NT_2_080223 | 1 | SMCSD | NT | AdV5 | 182.8 | 8246 | 452 | 7794 | 2027 | 1 |
| F3 | SMCSD_NT_3_080223 | 1 | SMCSD | NT | AdV5 | 124.2 | 8273 | 309 | 7964 | 2027 | 1 |
| D3 | SMCSD_NT_1_091923 | 2 | SMCSD | NT | AdV5 | 93.07 | 8284 | 234 | 8050 | 2038 | 1 |
| E3 | SMCSD_NT_2_091923 | 2 | SMCSD | NT | AdV5 | 118.6 | 8273 | 297 | 7976 | 2038 | 1 |
| F3 | SMCSD_NT_3_091923 | 2 | SMCSD | NT | AdV5 | 95.47 | 8251 | 238 | 8013 | 2038 | 1 |
| D1 | SMCSD_NT_1_080223 | 1 | SMCSD | NT | B5 | 146.7 | 8121 | 373 | 7748 | 2027 | 1 |
| E1 | SMCSD_NT_2_080223 | 1 | SMCSD | NT | B5 | 156.2 | 8276 | 399 | 7877 | 2027 | 1 |
| F1 | SMCSD_NT_3_080223 | 1 | SMCSD | NT | B5 | 105.2 | 8253 | 269 | 7984 | 2027 | 1 |
| D1 | SMCSD_NT_1_091923 | 2 | SMCSD | NT | B5 | 43.47 | 8165 | 113 | 8052 | 2038 | 1 |
| E1 | SMCSD_NT_2_091923 | 2 | SMCSD | NT | B5 | 45.33 | 8288 | 118 | 8170 | 2038 | 1 |
| F1 | SMCSD_NT_3_091923 | 2 | SMCSD | NT | B5 | 50.69 | 8208 | 130 | 8078 | 2038 | 1 |
| D5 | SMCSD_NT_1_080223 | 1 | SMCSD | NT | H1N1 | 4.361 | 8248 | 11 | 8237 | 2027 | 1 |
| E5 | SMCSD_NT_2_080223 | 1 | SMCSD | NT | H1N1 | 5.28 | 8255 | 13 | 8242 | 2027 | 1 |
| F5 | SMCSD_NT_3_080223 | 1 | SMCSD | NT | H1N1 | 3.26 | 8252 | 8 | 8244 | 2027 | 1 |
| D5 | SMCSD_NT_1_091923 | 2 | SMCSD | NT | H1N1 | 1.185 | 8277 | 3 | 8274 | 2038 | 1 |
| E5 | SMCSD_NT_2_091923 | 2 | SMCSD | NT | H1N1 | 1.63 | 8225 | 4 | 8221 | 2038 | 1 |
| F5 | SMCSD_NT_3_091923 | 2 | SMCSD | NT | H1N1 | 2.039 | 8245 | 5 | 8240 | 2038 | 1 |
| D5 | SMCSD_NT_1_080223 | 1 | SMCSD | NT | H3N2 | 19.07 | 8248 | 48 | 8200 | 2027 | 1 |
| E5 | SMCSD_NT_2_080223 | 1 | SMCSD | NT | H3N2 | 17.91 | 8255 | 44 | 8211 | 2027 | 1 |
| F5 | SMCSD_NT_3_080223 | 1 | SMCSD | NT | H3N2 | 12.24 | 8252 | 30 | 8222 | 2027 | 1 |
| D5 | SMCSD_NT_1_091923 | 2 | SMCSD | NT | H3N2 | 2.37 | 8277 | 6 | 8271 | 2038 | 1 |
| E5 | SMCSD_NT_2_091923 | 2 | SMCSD | NT | H3N2 | 3.26 | 8225 | 8 | 8217 | 2038 | 1 |
| F5 | SMCSD_NT_3_091923 | 2 | SMCSD | NT | H3N2 | 2.039 | 8245 | 5 | 8240 | 2038 | 1 |
| D7 | SMCSD_NT_1_080223 | 1 | SMCSD | NT | N1 | 2457.2 | 8269 | 4308 | 3961 | 2027 | 1 |

| | | | | | | | | | | | |
|---|---|---|---|---|---|---|---|---|---|---|---|
| E7 | SMCSD_NT_2_080223 | 1 | SMCSD | NT | N1 | 2398.3 | 8262 | 4202 | 4060 | 2027 | 1 |
| F7 | SMCSD_NT_3_080223 | 1 | SMCSD | NT | N1 | 1360.4 | 8137 | 2719 | 5418 | 2027 | 1 |
| D7 | SMCSD_NT_1_091923 | 2 | SMCSD | NT | N1 | 833.7 | 8204 | 1813 | 6391 | 2038 | 1 |
| E7 | SMCSD_NT_2_091923 | 2 | SMCSD | NT | N1 | 883.3 | 8218 | 1892 | 6326 | 2038 | 1 |
| F7 | SMCSD_NT_3_091923 | 2 | SMCSD | NT | N1 | 695.3 | 8222 | 1543 | 6679 | 2038 | 1 |
| D7 | SMCSD_NT_1_080223 | 1 | SMCSD | NT | OC43 | 1562.7 | 8173 | 3055 | 5118 | 2027 | 1 |
| E7 | SMCSD_NT_2_080223 | 1 | SMCSD | NT | OC43 | 1532.4 | 8208 | 2995 | 5213 | 2027 | 1 |
| F7 | SMCSD_NT_3_080223 | 1 | SMCSD | NT | OC43 | 858.7 | 8251 | 1868 | 6383 | 2027 | 1 |
| D7 | SMCSD_NT_1_091923 | 2 | SMCSD | NT | OC43 | 555.7 | 8237 | 1263 | 6974 | 2038 | 1 |
| E7 | SMCSD_NT_2_091923 | 2 | SMCSD | NT | OC43 | 585.4 | 8228 | 1310 | 6918 | 2038 | 1 |
| F7 | SMCSD_NT_3_091923 | 2 | SMCSD | NT | OC43 | 491 | 8241 | 1125 | 7116 | 2038 | 1 |
| A1 | SMCSD_PMG_1_080223 | 1 | SMCSD | PMG | A6 | 155.8 | 8200 | 409 | 7791 | 2027 | 1 |
| B1 | SMCSD_PMG_2_080223 | 1 | SMCSD | PMG | A6 | 232.2 | 7418 | 532 | 6886 | 2027 | 1 |
| C1 | SMCSD_PMG_3_080223 | 1 | SMCSD | PMG | A6 | 126.8 | 7424 | 296 | 7128 | 2027 | 1 |
| A1 | SMCSD_PMG_1_091923 | 2 | SMCSD | PMG | A6 | 12.12 | 8055 | 32 | 8023 | 2038 | 1 |
| B1 | SMCSD_PMG_2_091923 | 2 | SMCSD | PMG | A6 | 33.59 | 8219 | 88 | 8131 | 2038 | 1 |
| C1 | SMCSD_PMG_3_091923 | 2 | SMCSD | PMG | A6 | 34.9 | 8263 | 92 | 8171 | 2038 | 1 |
| A3 | SMCSD_PMG_1_080223 | 1 | SMCSD | PMG | AdV2 | 114.9 | 8221 | 338 | 7883 | 2042 | 1 |
| B3 | SMCSD_PMG_2_080223 | 1 | SMCSD | PMG | AdV2 | 188.4 | 8181 | 521 | 7660 | 2042 | 1 |
| C3 | SMCSD_PMG_3_080223 | 1 | SMCSD | PMG | AdV2 | 302.5 | 8201 | 818 | 7383 | 2042 | 1 |
| A3 | SMCSD_PMG_1_091923 | 2 | SMCSD | PMG | AdV2 | 0 | 8178 | 0 | 8178 | 2038 | 1 |
| B3 | SMCSD_PMG_2_091923 | 2 | SMCSD | PMG | AdV2 | 0 | 8239 | 0 | 8239 | 2038 | 1 |
| C3 | SMCSD_PMG_3_091923 | 2 | SMCSD | PMG | AdV2 | 0.393 | 8274 | 1 | 8273 | 2038 | 1 |
| A3 | SMCSD_PMG_1_080223 | 1 | SMCSD | PMG | AdV5 | 93.44 | 8221 | 276 | 7945 | 2042 | 1 |
| B3 | SMCSD_PMG_2_080223 | 1 | SMCSD | PMG | AdV5 | 272 | 8178 | 741 | 7437 | 2042 | 1 |
| C3 | SMCSD_PMG_3_080223 | 1 | SMCSD | PMG | AdV5 | 447.2 | 8201 | 1180 | 7021 | 2042 | 1 |
| A3 | SMCSD_PMG_1_091923 | 2 | SMCSD | PMG | AdV5 | 10.17 | 7805 | 27 | 7778 | 15.95 | 1 |
| B3 | SMCSD_PMG_2_091923 | 2 | SMCSD | PMG | AdV5 | 6671.3 | 8246 | 7397 | 849 | 18.25 | 1 |
| C3 | SMCSD_PMG_3_091923 | 2 | SMCSD | PMG | AdV5 | 9.156 | 7921 | 25 | 7896 | 14.73 | 1 |
| A1 | SMCSD_PMG_1_080223 | 1 | SMCSD | PMG | B5 | 477 | 8195 | 1188 | 7007 | 2027 | 1 |
| B1 | SMCSD_PMG_2_080223 | 1 | SMCSD | PMG | B5 | 563.8 | 7405 | 1224 | 6181 | 2027 | 1 |
| C1 | SMCSD_PMG_3_080223 | 1 | SMCSD | PMG | B5 | 338.8 | 7418 | 764 | 6654 | 2027 | 1 |
| A1 | SMCSD_PMG_1_091923 | 2 | SMCSD | PMG | B5 | 52.25 | 8055 | 137 | 7918 | 2038 | 1 |
| B1 | SMCSD_PMG_2_091923 | 2 | SMCSD | PMG | B5 | 150.9 | 8219 | 388 | 7831 | 2038 | 1 |
| C1 | SMCSD_PMG_3_091923 | 2 | SMCSD | PMG | B5 | 142.1 | 8257 | 368 | 7889 | 2038 | 1 |
| A5 | SMCSD_PMG_1_080223 | 1 | SMCSD | PMG | H1N1 | 14.89 | 8191 | 39 | 8152 | 2027 | 1 |
| B5 | SMCSD_PMG_2_080223 | 1 | SMCSD | PMG | H1N1 | 27.93 | 8019 | 69 | 7950 | 2027 | 1 |
| C5 | SMCSD_PMG_3_080223 | 1 | SMCSD | PMG | H1N1 | 17.83 | 8268 | 45 | 8223 | 2027 | 1 |
| A5 | SMCSD_PMG_1_091923 | 2 | SMCSD | PMG | H1N1 | 1.533 | 8142 | 4 | 8138 | 2038 | 1 |
| B5 | SMCSD_PMG_2_091923 | 2 | SMCSD | PMG | H1N1 | 2.379 | 8155 | 6 | 8149 | 2038 | 1 |
| C5 | SMCSD_PMG_3_091923 | 2 | SMCSD | PMG | H1N1 | 1.58 | 8272 | 4 | 8268 | 2038 | 1 |
| A5 | SMCSD_PMG_1_080223 | 1 | SMCSD | PMG | H3N2 | 77.53 | 8191 | 201 | 7990 | 2027 | 1 |
| B5 | SMCSD_PMG_2_080223 | 1 | SMCSD | PMG | H3N2 | 106.1 | 8019 | 259 | 7760 | 2027 | 1 |
| C5 | SMCSD_PMG_3_080223 | 1 | SMCSD | PMG | H3N2 | 63.43 | 8268 | 159 | 8109 | 2027 | 1 |
| A5 | SMCSD_PMG_1_091923 | 2 | SMCSD | PMG | H3N2 | 8.828 | 8142 | 23 | 8119 | 2038 | 1 |

| | | | | | | | | | | | |
|---|---|---|---|---|---|---|---|---|---|---|---|
| B5 | SMCSD_PMG_2_091923 | 2 | SMCSD | PMG | H3N2 | 10.32 | 8155 | 26 | 8129 | 2038 | 1 |
| C5 | SMCSD_PMG_3_091923 | 2 | SMCSD | PMG | H3N2 | 15.44 | 8272 | 39 | 8233 | 2038 | 1 |
| A7 | SMCSD_PMG_1_080223 | 1 | SMCSD | PMG | N1 | 2000.6 | 8277 | 3873 | 4404 | 2027 | 1 |
| B7 | SMCSD_PMG_2_080223 | 1 | SMCSD | PMG | N1 | 2523.6 | 8214 | 4465 | 3749 | 2027 | 1 |
| C7 | SMCSD_PMG_3_080223 | 1 | SMCSD | PMG | N1 | 1970.6 | 8039 | 3626 | 4413 | 2027 | 1 |
| A7 | SMCSD_PMG_1_091923 | 2 | SMCSD | PMG | N1 | 523.8 | 8162 | 1243 | 6919 | 2038 | 1 |
| B7 | SMCSD_PMG_2_091923 | 2 | SMCSD | PMG | N1 | 671.5 | 8175 | 1540 | 6635 | 2038 | 1 |
| C7 | SMCSD_PMG_3_091923 | 2 | SMCSD | PMG | N1 | 491.6 | 8261 | 1148 | 7113 | 2038 | 1 |
| A7 | SMCSD_PMG_1_080223 | 1 | SMCSD | PMG | OC43 | 1195.6 | 8200 | 2576 | 5624 | 2027 | 1 |
| B7 | SMCSD_PMG_2_080223 | 1 | SMCSD | PMG | OC43 | 2120 | 8214 | 3964 | 4250 | 2027 | 1 |
| C7 | SMCSD_PMG_3_080223 | 1 | SMCSD | PMG | OC43 | 1610.8 | 7935 | 3075 | 4860 | 2027 | 1 |
| A7 | SMCSD_PMG_1_091923 | 2 | SMCSD | PMG | OC43 | 230.4 | 8162 | 572 | 7590 | 2038 | 1 |
| B7 | SMCSD_PMG_2_091923 | 2 | SMCSD | PMG | OC43 | 449.6 | 8204 | 1070 | 7134 | 2038 | 1 |
| C7 | SMCSD_PMG_3_091923 | 2 | SMCSD | PMG | OC43 | 287.8 | 8262 | 693 | 7569 | 2038 | 1 |
| G1 | SMCSD_S_1_080223 | 1 | SMCSD | S | A6 | 52.82 | 8262 | 138 | 8124 | 2027 | 1 |
| H1 | SMCSD_S_2_080223 | 1 | SMCSD | S | A6 | 17.84 | 8249 | 48 | 8201 | 2027 | 1 |
| A2 | SMCSD_S_3_080223 | 1 | SMCSD | S | A6 | 30.47 | 8094 | 79 | 8015 | 2027 | 1 |
| G1 | SMCSD_S_1_091923 | 2 | SMCSD | S | A6 | 38.51 | 8275 | 101 | 8174 | 2038 | 1 |
| H1 | SMCSD_S_2_091923 | 2 | SMCSD | S | A6 | 22.71 | 8242 | 61 | 8181 | 2038 | 1 |
| A2 | SMCSD_S_3_091923 | 2 | SMCSD | S | A6 | 42.45 | 8253 | 112 | 8141 | 2038 | 1 |
| G3 | SMCSD_S_1_080223 | 1 | SMCSD | S | AdV2 | 161.8 | 8169 | 399 | 7770 | 2027 | 1 |
| H3 | SMCSD_S_2_080223 | 1 | SMCSD | S | AdV2 | 130.7 | 8267 | 337 | 7930 | 2027 | 1 |
| A4 | SMCSD_S_3_080223 | 1 | SMCSD | S | AdV2 | 135.5 | 7750 | 329 | 7421 | 2027 | 1 |
| G3 | SMCSD_S_1_091923 | 2 | SMCSD | S | AdV2 | 192.8 | 8181 | 474 | 7707 | 2038 | 1 |
| H3 | SMCSD_S_2_091923 | 2 | SMCSD | S | AdV2 | 204.9 | 8265 | 522 | 7743 | 2038 | 1 |
| A4 | SMCSD_S_3_091923 | 2 | SMCSD | S | AdV2 | 217.3 | 7066 | 475 | 6591 | 2038 | 1 |
| G3 | SMCSD_S_1_080223 | 1 | SMCSD | S | AdV5 | 102.9 | 8169 | 256 | 7913 | 2027 | 1 |
| H3 | SMCSD_S_2_080223 | 1 | SMCSD | S | AdV5 | 92.94 | 8267 | 241 | 8026 | 2027 | 1 |
| A4 | SMCSD_S_3_080223 | 1 | SMCSD | S | AdV5 | 90.37 | 7748 | 221 | 7527 | 2027 | 1 |
| G3 | SMCSD_S_1_091923 | 2 | SMCSD | S | AdV5 | 118.2 | 8181 | 294 | 7887 | 2038 | 1 |
| H3 | SMCSD_S_2_091923 | 2 | SMCSD | S | AdV5 | 147.1 | 8265 | 378 | 7887 | 2038 | 1 |
| A4 | SMCSD_S_3_091923 | 2 | SMCSD | S | AdV5 | 138.6 | 8156 | 354 | 7802 | 2038 | 1 |
| G1 | SMCSD_S_1_080223 | 1 | SMCSD | S | B5 | 115.2 | 8264 | 298 | 7966 | 2027 | 1 |
| H1 | SMCSD_S_2_080223 | 1 | SMCSD | S | B5 | 54.59 | 8249 | 146 | 8103 | 2027 | 1 |
| A2 | SMCSD_S_3_080223 | 1 | SMCSD | S | B5 | 81.28 | 8094 | 209 | 7885 | 2027 | 1 |
| G1 | SMCSD_S_1_091923 | 2 | SMCSD | S | B5 | 48.88 | 8275 | 128 | 8147 | 2038 | 1 |
| H1 | SMCSD_S_2_091923 | 2 | SMCSD | S | B5 | 17.11 | 8242 | 46 | 8196 | 2038 | 1 |
| A2 | SMCSD_S_3_091923 | 2 | SMCSD | S | B5 | 63.9 | 8253 | 168 | 8085 | 2038 | 1 |
| G5 | SMCSD_S_1_080223 | 1 | SMCSD | S | H1N1 | 0.793 | 8272 | 2 | 8270 | 2027 | 1 |
| H5 | SMCSD_S_2_080223 | 1 | SMCSD | S | H1N1 | 0.758 | 8264 | 2 | 8262 | 2027 | 1 |
| A6 | SMCSD_S_3_080223 | 1 | SMCSD | S | H1N1 | 0.771 | 8048 | 2 | 8046 | 2027 | 1 |
| G5 | SMCSD_S_1_091923 | 2 | SMCSD | S | H1N1 | 0 | 8222 | 0 | 8222 | 2038 | 1 |
| H5 | SMCSD_S_2_091923 | 2 | SMCSD | S | H1N1 | 0 | 8201 | 0 | 8201 | 2038 | 1 |
| A6 | SMCSD_S_3_091923 | 2 | SMCSD | S | H1N1 | 0 | 8086 | 0 | 8086 | 2038 | 1 |
| G5 | SMCSD_S_1_080223 | 1 | SMCSD | S | H3N2 | 1.19 | 8272 | 3 | 8269 | 2027 | 1 |

| | | | | | | | | | | | |
|---|---|---|---|---|---|---|---|---|---|---|---|
| H5 | SMCSD_S_2_080223 | 1 | SMCSD | S | H3N2 | 2.275 | 8264 | 6 | 8258 | 2027 | 1 |
| A6 | SMCSD_S_3_080223 | 1 | SMCSD | S | H3N2 | 2.313 | 8048 | 6 | 8042 | 2027 | 1 |
| G5 | SMCSD_S_1_091923 | 2 | SMCSD | S | H3N2 | 0 | 8222 | 0 | 8222 | 2038 | 1 |
| H5 | SMCSD_S_2_091923 | 2 | SMCSD | S | H3N2 | 0 | 8201 | 0 | 8201 | 2038 | 1 |
| A6 | SMCSD_S_3_091923 | 2 | SMCSD | S | H3N2 | 0 | 8086 | 0 | 8086 | 2038 | 1 |
| G7 | SMCSD_S_1_080223 | 1 | SMCSD | S | N1 | 28.96 | 8269 | 72 | 8197 | 2027 | 1 |
| H7 | SMCSD_S_2_080223 | 1 | SMCSD | S | N1 | 28.6 | 8260 | 75 | 8185 | 2027 | 1 |
| A8 | SMCSD_S_3_080223 | 1 | SMCSD | S | N1 | 50.42 | 8182 | 132 | 8050 | 2027 | 1 |
| G7 | SMCSD_S_1_091923 | 2 | SMCSD | S | N1 | 40.08 | 8229 | 99 | 8130 | 2038 | 1 |
| H7 | SMCSD_S_2_091923 | 2 | SMCSD | S | N1 | 16.84 | 8216 | 44 | 8172 | 2038 | 1 |
| A8 | SMCSD_S_3_091923 | 2 | SMCSD | S | N1 | 19.04 | 8167 | 50 | 8117 | 2038 | 1 |
| G7 | SMCSD_S_1_080223 | 1 | SMCSD | S | OC43 | 152.4 | 8269 | 372 | 7897 | 2027 | 1 |
| H7 | SMCSD_S_2_080223 | 1 | SMCSD | S | OC43 | 131.4 | 8260 | 339 | 7921 | 2027 | 1 |
| A8 | SMCSD_S_3_080223 | 1 | SMCSD | S | OC43 | 151.4 | 8182 | 390 | 7792 | 2027 | 1 |
| G7 | SMCSD_S_1_091923 | 2 | SMCSD | S | OC43 | 114.7 | 8227 | 280 | 7947 | 2038 | 1 |
| H7 | SMCSD_S_2_091923 | 2 | SMCSD | S | OC43 | 49.62 | 8216 | 129 | 8087 | 2038 | 1 |
| A8 | SMCSD_S_3_091923 | 2 | SMCSD | S | OC43 | 57.86 | 8167 | 151 | 8016 | 2038 | 1 |
| B2 | WCSD_IP_1_073123 | 1 | WCSD | IP | A6 | 53.39 | 8165 | 138 | 8027 | 2025 | 1 |
| C2 | WCSD_IP_2_073123 | 1 | WCSD | IP | A6 | 61.72 | 7459 | 143 | 7316 | 2025 | 1 |
| D2 | WCSD_IP_3_073123 | 1 | WCSD | IP | A6 | 35.7 | 7604 | 84 | 7520 | 2025 | 1 |
| B2 | WCSD_IP_1_090523 | 2 | WCSD | IP | A6 | 76.41 | 8091 | 195 | 7896 | 2035 | 1 |
| C2 | WCSD_IP_2_090523 | 2 | WCSD | IP | A6 | 106.6 | 8214 | 270 | 7944 | 2035 | 1 |
| D2 | WCSD_IP_3_090523 | 2 | WCSD | IP | A6 | 137.5 | 8049 | 337 | 7712 | 2035 | 1 |
| B4 | WCSD_IP_1_073123 | 1 | WCSD | IP | AdV2 | 96.96 | 8216 | 244 | 7972 | 2025 | 1 |
| C4 | WCSD_IP_2_073123 | 1 | WCSD | IP | AdV2 | 141.3 | 8293 | 356 | 7937 | 2025 | 1 |
| D4 | WCSD_IP_3_073123 | 1 | WCSD | IP | AdV2 | 129.5 | 8287 | 322 | 7965 | 2025 | 1 |
| B4 | WCSD_IP_1_090523 | 2 | WCSD | IP | AdV2 | 323 | 8164 | 780 | 7384 | 2035 | 1 |
| C4 | WCSD_IP_2_090523 | 2 | WCSD | IP | AdV2 | 440.7 | 8240 | 1054 | 7186 | 2035 | 1 |
| D4 | WCSD_IP_3_090523 | 2 | WCSD | IP | AdV2 | 105.5 | 8276 | 263 | 8013 | 2035 | 1 |
| B4 | WCSD_IP_1_073123 | 1 | WCSD | IP | AdV5 | 108.7 | 8216 | 273 | 7943 | 2025 | 1 |
| C4 | WCSD_IP_2_073123 | 1 | WCSD | IP | AdV5 | 130.8 | 8290 | 330 | 7960 | 2025 | 1 |
| D4 | WCSD_IP_3_073123 | 1 | WCSD | IP | AdV5 | 128.6 | 8291 | 320 | 7971 | 2025 | 1 |
| B4 | WCSD_IP_1_090523 | 2 | WCSD | IP | AdV5 | 407.1 | 8158 | 970 | 7188 | 2035 | 1 |
| C4 | WCSD_IP_2_090523 | 2 | WCSD | IP | AdV5 | 565.1 | 8219 | 1323 | 6896 | 2035 | 1 |
| D4 | WCSD_IP_3_090523 | 2 | WCSD | IP | AdV5 | 129.3 | 8272 | 321 | 7951 | 2035 | 1 |
| B2 | WCSD_IP_1_073123 | 1 | WCSD | IP | B5 | 1408.4 | 8085 | 2928 | 5157 | 2025 | 1 |
| C2 | WCSD_IP_2_073123 | 1 | WCSD | IP | B5 | 1484.3 | 7343 | 2733 | 4610 | 2025 | 1 |
| D2 | WCSD_IP_3_073123 | 1 | WCSD | IP | B5 | 1001.9 | 7572 | 2028 | 5544 | 2025 | 1 |
| B2 | WCSD_IP_1_090523 | 2 | WCSD | IP | B5 | 252.2 | 8093 | 626 | 7467 | 2035 | 1 |
| C2 | WCSD_IP_2_090523 | 2 | WCSD | IP | B5 | 277.3 | 8211 | 684 | 7527 | 2035 | 1 |
| D2 | WCSD_IP_3_090523 | 2 | WCSD | IP | B5 | 371.8 | 8055 | 880 | 7175 | 2035 | 1 |
| B6 | WCSD_IP_1_073123 | 1 | WCSD | IP | H1N1 | 8.699 | 7424 | 20 | 7404 | 2025 | 1 |
| C6 | WCSD_IP_2_073123 | 1 | WCSD | IP | H1N1 | 11.15 | 8050 | 27 | 8023 | 2025 | 1 |
| D6 | WCSD_IP_3_073123 | 1 | WCSD | IP | H1N1 | 7.187 | 8241 | 18 | 8223 | 2025 | 1 |
| B6 | WCSD_IP_1_090523 | 2 | WCSD | IP | H1N1 | 0.782 | 8245 | 2 | 8243 | 2035 | 1 |

| | | | | | | | | | | | |
|---|---|---|---|---|---|---|---|---|---|---|---|
| C6 | WCSD_IP_2_090523 | 2 | WCSD | IP | H1N1 | 0.403 | 8227 | 1 | 8226 | 2035 | 1 |
| D6 | WCSD_IP_3_090523 | 2 | WCSD | IP | H1N1 | 0.804 | 8173 | 2 | 8171 | 2035 | 1 |
| B6 | WCSD_IP_1_073123 | 1 | WCSD | IP | H3N2 | 17.42 | 7424 | 40 | 7384 | 2025 | 1 |
| C6 | WCSD_IP_2_073123 | 1 | WCSD | IP | H3N2 | 16.95 | 8050 | 41 | 8009 | 2025 | 1 |
| D6 | WCSD_IP_3_073123 | 1 | WCSD | IP | H3N2 | 17.2 | 8241 | 43 | 8198 | 2025 | 1 |
| B6 | WCSD_IP_1_090523 | 2 | WCSD | IP | H3N2 | 1.565 | 8245 | 4 | 8241 | 2035 | 1 |
| C6 | WCSD_IP_2_090523 | 2 | WCSD | IP | H3N2 | 0 | 8227 | 0 | 8227 | 2035 | 1 |
| D6 | WCSD_IP_3_090523 | 2 | WCSD | IP | H3N2 | 1.609 | 8173 | 4 | 8169 | 2035 | 1 |
| B8 | WCSD_IP_1_073123 | 1 | WCSD | IP | N1 | 534 | 8037 | 1212 | 6825 | 2025 | 1 |
| C8 | WCSD_IP_2_073123 | 1 | WCSD | IP | N1 | 363.8 | 8227 | 860 | 7367 | 2025 | 1 |
| D8 | WCSD_IP_3_073123 | 1 | WCSD | IP | N1 | 173.2 | 8262 | 421 | 7841 | 2025 | 1 |
| B8 | WCSD_IP_1_090523 | 2 | WCSD | IP | N1 | 189.5 | 8252 | 465 | 7787 | 2035 | 1 |
| C8 | WCSD_IP_2_090523 | 2 | WCSD | IP | N1 | 185.5 | 8275 | 453 | 7822 | 2035 | 1 |
| D8 | WCSD_IP_3_090523 | 2 | WCSD | IP | N1 | 101.5 | 8278 | 250 | 8028 | 2035 | 1 |
| B8 | WCSD_IP_1_073123 | 1 | WCSD | IP | OC43 | 2362.5 | 8046 | 4142 | 3904 | 2025 | 1 |
| C8 | WCSD_IP_2_073123 | 1 | WCSD | IP | OC43 | 2208 | 8227 | 4018 | 4209 | 2025 | 1 |
| D8 | WCSD_IP_3_073123 | 1 | WCSD | IP | OC43 | 1483.8 | 8229 | 2972 | 5257 | 2025 | 1 |
| B8 | WCSD_IP_1_090523 | 2 | WCSD | IP | OC43 | 508.7 | 8239 | 1188 | 7051 | 2035 | 1 |
| C8 | WCSD_IP_2_090523 | 2 | WCSD | IP | OC43 | 343.5 | 8272 | 819 | 7453 | 2035 | 1 |
| D8 | WCSD_IP_3_090523 | 2 | WCSD | IP | OC43 | 452.2 | 8265 | 1055 | 7210 | 2035 | 1 |
| D1 | WCSD_NT_1_073123 | 1 | WCSD | NT | A6 | 426.5 | 7504 | 959 | 6545 | 2025 | 1 |
| E1 | WCSD_NT_2_073123 | 1 | WCSD | NT | A6 | 467.4 | 8265 | 1136 | 7129 | 2025 | 1 |
| F1 | WCSD_NT_3_073123 | 1 | WCSD | NT | A6 | 448.4 | 8247 | 1086 | 7161 | 2025 | 1 |
| D1 | WCSD_NT_1_090523 | 2 | WCSD | NT | A6 | 151.4 | 8040 | 381 | 7659 | 2035 | 1 |
| E1 | WCSD_NT_2_090523 | 2 | WCSD | NT | A6 | 255.4 | 8079 | 627 | 7452 | 2035 | 1 |
| F1 | WCSD_NT_3_090523 | 2 | WCSD | NT | A6 | 186.4 | 8104 | 462 | 7642 | 2035 | 1 |
| D3 | WCSD_NT_1_073123 | 1 | WCSD | NT | AdV2 | 625.7 | 8241 | 1444 | 6797 | 2025 | 1 |
| E3 | WCSD_NT_2_073123 | 1 | WCSD | NT | AdV2 | 507.4 | 8272 | 1198 | 7074 | 2025 | 1 |
| F3 | WCSD_NT_3_073123 | 1 | WCSD | NT | AdV2 | 605.1 | 8263 | 1399 | 6864 | 2025 | 1 |
| D3 | WCSD_NT_1_090523 | 2 | WCSD | NT | AdV2 | 438 | 8252 | 1041 | 7211 | 2035 | 1 |
| E3 | WCSD_NT_2_090523 | 2 | WCSD | NT | AdV2 | 785.8 | 8244 | 1774 | 6470 | 2035 | 1 |
| F3 | WCSD_NT_3_090523 | 2 | WCSD | NT | AdV2 | 608.9 | 8227 | 1401 | 6826 | 2035 | 1 |
| D3 | WCSD_NT_1_073123 | 1 | WCSD | NT | AdV5 | 338.2 | 8241 | 815 | 7426 | 2025 | 1 |
| E3 | WCSD_NT_2_073123 | 1 | WCSD | NT | AdV5 | 271.2 | 8290 | 665 | 7625 | 2025 | 1 |
| F3 | WCSD_NT_3_073123 | 1 | WCSD | NT | AdV5 | 339.9 | 8276 | 819 | 7457 | 2025 | 1 |
| D3 | WCSD_NT_1_090523 | 2 | WCSD | NT | AdV5 | 236.5 | 8259 | 580 | 7679 | 2035 | 1 |
| E3 | WCSD_NT_2_090523 | 2 | WCSD | NT | AdV5 | 411.5 | 8275 | 986 | 7289 | 2035 | 1 |
| F3 | WCSD_NT_3_090523 | 2 | WCSD | NT | AdV5 | 272.4 | 8239 | 660 | 7579 | 2035 | 1 |
| D1 | WCSD_NT_1_073123 | 1 | WCSD | NT | B5 | 18.35 | 7504 | 44 | 7460 | 2025 | 1 |
| E1 | WCSD_NT_2_073123 | 1 | WCSD | NT | B5 | 19.17 | 8271 | 50 | 8221 | 2025 | 1 |
| F1 | WCSD_NT_3_073123 | 1 | WCSD | NT | B5 | 26.25 | 8259 | 68 | 8191 | 2025 | 1 |
| D1 | WCSD_NT_1_090523 | 2 | WCSD | NT | B5 | 6.992 | 8040 | 18 | 8022 | 2035 | 1 |
| E1 | WCSD_NT_2_090523 | 2 | WCSD | NT | B5 | 15.3 | 8079 | 39 | 8040 | 2035 | 1 |
| F1 | WCSD_NT_3_090523 | 2 | WCSD | NT | B5 | 11.78 | 8104 | 30 | 8074 | 2035 | 1 |
| D5 | WCSD_NT_1_073123 | 1 | WCSD | NT | H1N1 | 10.09 | 7782 | 24 | 7758 | 2025 | 1 |

| | | | | | | | | | | | |
|---|---|---|---|---|---|---|---|---|---|---|---|
| E5 | WCSD_NT_2_073123 | 1 | WCSD | NT | H1N1 | 15 | 8060 | 36 | 8024 | 2025 | 1 |
| F5 | WCSD_NT_3_073123 | 1 | WCSD | NT | H1N1 | 10.76 | 8134 | 26 | 8108 | 2025 | 1 |
| D5 | WCSD_NT_1_090523 | 2 | WCSD | NT | H1N1 | 0 | 4704 | 0 | 4704 | 2035 | 1 |
| E5 | WCSD_NT_2_090523 | 2 | WCSD | NT | H1N1 | 0.406 | 8251 | 1 | 8250 | 2035 | 1 |
| F5 | WCSD_NT_3_090523 | 2 | WCSD | NT | H1N1 | 1.228 | 8214 | 3 | 8211 | 2035 | 1 |
| D5 | WCSD_NT_1_073123 | 1 | WCSD | NT | H3N2 | 17.26 | 7782 | 41 | 7741 | 2025 | 1 |
| E5 | WCSD_NT_2_073123 | 1 | WCSD | NT | H3N2 | 22.1 | 8060 | 53 | 8007 | 2025 | 1 |
| F5 | WCSD_NT_3_073123 | 1 | WCSD | NT | H3N2 | 17.81 | 8134 | 43 | 8091 | 2025 | 1 |
| D5 | WCSD_NT_1_090523 | 2 | WCSD | NT | H3N2 | 2.085 | 4704 | 3 | 4701 | 2035 | 1 |
| E5 | WCSD_NT_2_090523 | 2 | WCSD | NT | H3N2 | 2.437 | 8251 | 6 | 8245 | 2035 | 1 |
| F5 | WCSD_NT_3_090523 | 2 | WCSD | NT | H3N2 | 2.456 | 8214 | 6 | 8208 | 2035 | 1 |
| D7 | WCSD_NT_1_073123 | 1 | WCSD | NT | N1 | 1938.9 | 8013 | 3530 | 4483 | 2025 | 1 |
| E7 | WCSD_NT_2_073123 | 1 | WCSD | NT | N1 | 1788.3 | 8260 | 3397 | 4863 | 2025 | 1 |
| F7 | WCSD_NT_3_073123 | 1 | WCSD | NT | N1 | 1944.1 | 8247 | 3635 | 4612 | 2025 | 1 |
| D7 | WCSD_NT_1_090523 | 2 | WCSD | NT | N1 | 617.9 | 8233 | 1391 | 6842 | 2035 | 1 |
| E7 | WCSD_NT_2_090523 | 2 | WCSD | NT | N1 | 939.7 | 8223 | 1998 | 6225 | 2035 | 1 |
| F7 | WCSD_NT_3_090523 | 2 | WCSD | NT | N1 | 802.7 | 8268 | 1764 | 6504 | 2035 | 1 |
| D7 | WCSD_NT_1_073123 | 1 | WCSD | NT | OC43 | 1144.4 | 7853 | 2279 | 5574 | 2025 | 1 |
| E7 | WCSD_NT_2_073123 | 1 | WCSD | NT | OC43 | 1180.6 | 8210 | 2423 | 5787 | 2025 | 1 |
| F7 | WCSD_NT_3_073123 | 1 | WCSD | NT | OC43 | 1015.1 | 8203 | 2147 | 6056 | 2025 | 1 |
| D7 | WCSD_NT_1_090523 | 2 | WCSD | NT | OC43 | 339 | 8276 | 799 | 7477 | 2035 | 1 |
| E7 | WCSD_NT_2_090523 | 2 | WCSD | NT | OC43 | 481.5 | 8283 | 1101 | 7182 | 2035 | 1 |
| F7 | WCSD_NT_3_090523 | 2 | WCSD | NT | OC43 | 442.2 | 8285 | 1026 | 7259 | 2035 | 1 |
| A1 | WCSD_PMG_1_073123 | 1 | WCSD | PMG | A6 | 447.4 | 7430 | 1015 | 6415 | 2025 | 1 |
| B1 | WCSD_PMG_2_073123 | 1 | WCSD | PMG | A6 | 220.3 | 8259 | 563 | 7696 | 2025 | 1 |
| C1 | WCSD_PMG_3_073123 | 1 | WCSD | PMG | A6 | 365.9 | 7421 | 822 | 6599 | 2025 | 1 |
| A1 | WCSD_PMG_1_090523 | 2 | WCSD | PMG | A6 | 223.3 | 8022 | 567 | 7455 | 2035 | 1 |
| B1 | WCSD_PMG_2_090523 | 2 | WCSD | PMG | A6 | 269.3 | 7898 | 653 | 7245 | 2035 | 1 |
| C1 | WCSD_PMG_3_090523 | 2 | WCSD | PMG | A6 | 329.3 | 8020 | 804 | 7216 | 2035 | 1 |
| A3 | WCSD_PMG_1_073123 | 1 | WCSD | PMG | AdV2 | 3109.2 | 8137 | 5137 | 3000 | 2025 | 1 |
| B3 | WCSD_PMG_2_073123 | 1 | WCSD | PMG | AdV2 | 1221.6 | 7810 | 2482 | 5328 | 2025 | 1 |
| C3 | WCSD_PMG_3_073123 | 1 | WCSD | PMG | AdV2 | 1544.9 | 7901 | 2987 | 4914 | 2025 | 1 |
| A3 | WCSD_PMG_1_090523 | 2 | WCSD | PMG | AdV2 | 463.1 | 8139 | 1124 | 7015 | 2035 | 1 |
| B3 | WCSD_PMG_2_090523 | 2 | WCSD | PMG | AdV2 | 545.9 | 8218 | 1291 | 6927 | 2035 | 1 |
| C3 | WCSD_PMG_3_090523 | 2 | WCSD | PMG | AdV2 | 1101.9 | 8210 | 2359 | 5851 | 2035 | 1 |
| A3 | WCSD_PMG_1_073123 | 1 | WCSD | PMG | AdV5 | 2850.3 | 8137 | 4877 | 3260 | 2025 | 1 |
| B3 | WCSD_PMG_2_073123 | 1 | WCSD | PMG | AdV5 | 1518 | 8032 | 3038 | 4994 | 2025 | 1 |
| C3 | WCSD_PMG_3_073123 | 1 | WCSD | PMG | AdV5 | 1551.8 | 8195 | 3109 | 5086 | 2025 | 1 |
| A3 | WCSD_PMG_1_090523 | 2 | WCSD | PMG | AdV5 | 476 | 8139 | 1153 | 6986 | 2035 | 1 |
| B3 | WCSD_PMG_2_090523 | 2 | WCSD | PMG | AdV5 | 537.5 | 8207 | 1271 | 6936 | 2035 | 1 |
| C3 | WCSD_PMG_3_090523 | 2 | WCSD | PMG | AdV5 | 1072.3 | 8169 | 2294 | 5875 | 2035 | 1 |
| A1 | WCSD_PMG_1_073123 | 1 | WCSD | PMG | B5 | 3816.9 | 7430 | 5308 | 2122 | 2025 | 1 |
| B1 | WCSD_PMG_2_073123 | 1 | WCSD | PMG | B5 | 1916.5 | 8259 | 3790 | 4469 | 2025 | 1 |
| C1 | WCSD_PMG_3_073123 | 1 | WCSD | PMG | B5 | 3067.4 | 7421 | 4647 | 2774 | 2025 | 1 |
| A1 | WCSD_PMG_1_090523 | 2 | WCSD | PMG | B5 | 760.1 | 8014 | 1770 | 6244 | 2035 | 1 |

| | | | | | | | | | | | |
|---|---|---|---|---|---|---|---|---|---|---|---|
| B1 | WCSD_PMG_2_090523 | 2 | WCSD | PMG | B5 | 771.3 | 7841 | 1717 | 6124 | 2035 | 1 |
| C1 | WCSD_PMG_3_090523 | 2 | WCSD | PMG | B5 | 858.7 | 7928 | 1909 | 6019 | 2035 | 1 |
| A5 | WCSD_PMG_1_073123 | 1 | WCSD | PMG | H1N1 | 91.85 | 7551 | 219 | 7332 | 2025 | 1 |
| B5 | WCSD_PMG_2_073123 | 1 | WCSD | PMG | H1N1 | 54.17 | 7340 | 122 | 7218 | 2025 | 1 |
| C5 | WCSD_PMG_3_073123 | 1 | WCSD | PMG | H1N1 | 81.49 | 8240 | 203 | 8037 | 2025 | 1 |
| A5 | WCSD_PMG_1_090523 | 2 | WCSD | PMG | H1N1 | 11.42 | 8216 | 30 | 8186 | 2035 | 1 |
| B5 | WCSD_PMG_2_090523 | 2 | WCSD | PMG | H1N1 | 16.36 | 8122 | 41 | 8081 | 2035 | 1 |
| C5 | WCSD_PMG_3_090523 | 2 | WCSD | PMG | H1N1 | 13.08 | 8261 | 33 | 8228 | 2035 | 1 |
| A5 | WCSD_PMG_1_073123 | 1 | WCSD | PMG | H3N2 | 260.6 | 7551 | 605 | 6946 | 2025 | 1 |
| B5 | WCSD_PMG_2_073123 | 1 | WCSD | PMG | H3N2 | 133 | 7340 | 296 | 7044 | 2025 | 1 |
| C5 | WCSD_PMG_3_073123 | 1 | WCSD | PMG | H3N2 | 238 | 8240 | 579 | 7661 | 2025 | 1 |
| A5 | WCSD_PMG_1_090523 | 2 | WCSD | PMG | H3N2 | 13.7 | 8216 | 36 | 8180 | 2035 | 1 |
| B5 | WCSD_PMG_2_090523 | 2 | WCSD | PMG | H3N2 | 21.16 | 8122 | 53 | 8069 | 2035 | 1 |
| C5 | WCSD_PMG_3_090523 | 2 | WCSD | PMG | H3N2 | 23.02 | 8261 | 58 | 8203 | 2035 | 1 |
| A7 | WCSD_PMG_1_073123 | 1 | WCSD | PMG | N1 | 8915.4 | 7455 | 7007 | 448 | 2025 | 1 |
| B7 | WCSD_PMG_2_073123 | 1 | WCSD | PMG | N1 | 6728.4 | 8087 | 7088 | 999 | 2025 | 1 |
| C7 | WCSD_PMG_3_073123 | 1 | WCSD | PMG | N1 | 8532.9 | 8215 | 7603 | 612 | 2025 | 1 |
| A7 | WCSD_PMG_1_090523 | 2 | WCSD | PMG | N1 | 2120.8 | 8263 | 4030 | 4233 | 2035 | 1 |
| B7 | WCSD_PMG_2_090523 | 2 | WCSD | PMG | N1 | 2692.2 | 8252 | 4678 | 3574 | 2035 | 1 |
| C7 | WCSD_PMG_3_090523 | 2 | WCSD | PMG | N1 | 3182.4 | 8179 | 5074 | 3105 | 2035 | 1 |
| A7 | WCSD_PMG_1_073123 | 1 | WCSD | PMG | OC43 | 6266.5 | 7455 | 6422 | 1033 | 2025 | 1 |
| B7 | WCSD_PMG_2_073123 | 1 | WCSD | PMG | OC43 | 2901.5 | 8087 | 4805 | 3282 | 2025 | 1 |
| C7 | WCSD_PMG_3_073123 | 1 | WCSD | PMG | OC43 | 5268.2 | 8215 | 6562 | 1653 | 2025 | 1 |
| A7 | WCSD_PMG_1_090523 | 2 | WCSD | PMG | OC43 | 1498.5 | 8170 | 3077 | 5093 | 2035 | 1 |
| B7 | WCSD_PMG_2_090523 | 2 | WCSD | PMG | OC43 | 1891.5 | 8252 | 3668 | 4584 | 2035 | 1 |
| C7 | WCSD_PMG_3_090523 | 2 | WCSD | PMG | OC43 | 3117.4 | 8179 | 5012 | 3167 | 2035 | 1 |
| G1 | WCSD_S_1_073123 | 1 | WCSD | S | A6 | 311.6 | 8236 | 779 | 7457 | 2025 | 1 |
| H1 | WCSD_S_2_073123 | 1 | WCSD | S | A6 | 152.5 | 8013 | 390 | 7623 | 2025 | 1 |
| A2 | WCSD_S_3_073123 | 1 | WCSD | S | A6 | 213 | 7439 | 493 | 6946 | 2025 | 1 |
| G1 | WCSD_S_1_090523 | 2 | WCSD | S | A6 | 58.33 | 8138 | 150 | 7988 | 2035 | 1 |
| H1 | WCSD_S_2_090523 | 2 | WCSD | S | A6 | 45.98 | 8171 | 122 | 8049 | 2035 | 1 |
| A2 | WCSD_S_3_090523 | 2 | WCSD | S | A6 | 56.2 | 8200 | 147 | 8053 | 2035 | 1 |
| G3 | WCSD_S_1_073123 | 1 | WCSD | S | AdV2 | 159 | 6246 | 300 | 5946 | 2025 | 1 |
| H3 | WCSD_S_2_073123 | 1 | WCSD | S | AdV2 | 1.14 | 8266 | 3 | 8263 | 2025 | 1 |
| A4 | WCSD_S_3_073123 | 1 | WCSD | S | AdV2 | 65.6 | 8178 | 170 | 8008 | 2025 | 1 |
| G3 | WCSD_S_1_090523 | 2 | WCSD | S | AdV2 | 32.73 | 8235 | 83 | 8152 | 2035 | 1 |
| H3 | WCSD_S_2_090523 | 2 | WCSD | S | AdV2 | 4.95 | 8256 | 13 | 8243 | 2035 | 1 |
| A4 | WCSD_S_3_090523 | 2 | WCSD | S | AdV2 | 91.51 | 8207 | 237 | 7970 | 2035 | 1 |
| G3 | WCSD_S_1_073123 | 1 | WCSD | S | AdV5 | 54.2 | 8175 | 136 | 8039 | 2025 | 1 |
| H3 | WCSD_S_2_073123 | 1 | WCSD | S | AdV5 | 0.38 | 8266 | 1 | 8265 | 2025 | 1 |
| A4 | WCSD_S_3_073123 | 1 | WCSD | S | AdV5 | 40.74 | 8178 | 106 | 8072 | 2025 | 1 |
| G3 | WCSD_S_1_090523 | 2 | WCSD | S | AdV5 | 16.13 | 8235 | 41 | 8194 | 2035 | 1 |
| H3 | WCSD_S_2_090523 | 2 | WCSD | S | AdV5 | 3.426 | 8256 | 9 | 8247 | 2035 | 1 |
| A4 | WCSD_S_3_090523 | 2 | WCSD | S | AdV5 | 46.77 | 8207 | 122 | 8085 | 2035 | 1 |
| G1 | WCSD_S_1_073123 | 1 | WCSD | S | B5 | 55.27 | 8242 | 144 | 8098 | 2025 | 1 |

| | | | | | | | | | | | |
|---|---|---|---|---|---|---|---|---|---|---|---|
| H1 | WCSD_S_2_073123 | 1 | WCSD | S | B5 | 28.34 | 8020 | 74 | 7946 | 2025 | 1 |
| A2 | WCSD_S_3_073123 | 1 | WCSD | S | B5 | 19.27 | 7439 | 46 | 7393 | 2025 | 1 |
| G1 | WCSD_S_1_090523 | 2 | WCSD | S | B5 | 11.19 | 8138 | 29 | 8109 | 2035 | 1 |
| H1 | WCSD_S_2_090523 | 2 | WCSD | S | B5 | 13.12 | 8170 | 35 | 8135 | 2035 | 1 |
| A2 | WCSD_S_3_090523 | 2 | WCSD | S | B5 | 7.587 | 8200 | 20 | 8180 | 2035 | 1 |
| G5 | WCSD_S_1_073123 | 1 | WCSD | S | H1N1 | 0.399 | 8215 | 1 | 8214 | 2025 | 1 |
| H5 | WCSD_S_2_073123 | 1 | WCSD | S | H1N1 | 0 | 8250 | 0 | 8250 | 2025 | 1 |
| A6 | WCSD_S_3_073123 | 1 | WCSD | S | H1N1 | 0.412 | 7536 | 1 | 7535 | 2025 | 1 |
| G5 | WCSD_S_1_090523 | 2 | WCSD | S | H1N1 | 0 | 8169 | 0 | 8169 | 2035 | 1 |
| H5 | WCSD_S_2_090523 | 2 | WCSD | S | H1N1 | 0 | 8236 | 0 | 8236 | 2035 | 1 |
| A6 | WCSD_S_3_090523 | 2 | WCSD | S | H1N1 | 0 | 8182 | 0 | 8182 | 2035 | 1 |
| G5 | WCSD_S_1_073123 | 1 | WCSD | S | H3N2 | 1.198 | 8215 | 3 | 8212 | 2025 | 1 |
| H5 | WCSD_S_2_073123 | 1 | WCSD | S | H3N2 | 0 | 8250 | 0 | 8250 | 2025 | 1 |
| A6 | WCSD_S_3_073123 | 1 | WCSD | S | H3N2 | 0 | 7536 | 0 | 7536 | 2025 | 1 |
| G5 | WCSD_S_1_090523 | 2 | WCSD | S | H3N2 | 0 | 8169 | 0 | 8169 | 2035 | 1 |
| H5 | WCSD_S_2_090523 | 2 | WCSD | S | H3N2 | 0 | 8236 | 0 | 8236 | 2035 | 1 |
| A6 | WCSD_S_3_090523 | 2 | WCSD | S | H3N2 | 0 | 8182 | 0 | 8182 | 2035 | 1 |
| G7 | WCSD_S_1_073123 | 1 | WCSD | S | N1 | 53.53 | 8231 | 132 | 8099 | 2025 | 1 |
| H7 | WCSD_S_2_073123 | 1 | WCSD | S | N1 | 0 | 8259 | 0 | 8259 | 2025 | 1 |
| A8 | WCSD_S_3_073123 | 1 | WCSD | S | N1 | 12.58 | 8152 | 33 | 8119 | 2025 | 1 |
| G7 | WCSD_S_1_090523 | 2 | WCSD | S | N1 | 1.2 | 8279 | 3 | 8276 | 2035 | 1 |
| H7 | WCSD_S_2_090523 | 2 | WCSD | S | N1 | 0.762 | 8232 | 2 | 8230 | 2035 | 1 |
| A8 | WCSD_S_3_090523 | 2 | WCSD | S | N1 | 4.5 | 8273 | 12 | 8261 | 2035 | 1 |
| G7 | WCSD_S_1_073123 | 1 | WCSD | S | OC43 | 229.9 | 8231 | 552 | 7679 | 2025 | 1 |
| H7 | WCSD_S_2_073123 | 1 | WCSD | S | OC43 | 5.7 | 8259 | 15 | 8244 | 2025 | 1 |
| A8 | WCSD_S_3_073123 | 1 | WCSD | S | OC43 | 58.74 | 8152 | 153 | 7999 | 2025 | 1 |
| G7 | WCSD_S_1_090523 | 2 | WCSD | S | OC43 | 0 | 8279 | 0 | 8279 | 2035 | 1 |
| H7 | WCSD_S_2_090523 | 2 | WCSD | S | OC43 | 2.667 | 8232 | 7 | 8225 | 2035 | 1 |
| A8 | WCSD_S_3_090523 | 2 | WCSD | S | OC43 | 6.753 | 8273 | 18 | 8255 | 2035 | 1 |
| B10 | EBMUD_IP_1_072623 | 1 | EBMUD | IP | crassphage | 2374.1 | 7489 | 4344 | 3145 | 2043 | 5 |
| C10 | EBMUD_IP_2_072623 | 1 | EBMUD | IP | crassphage | 2241.5 | 6600 | 3583 | 3017 | 2043 | 5 |
| D10 | EBMUD_IP_3_072623 | 1 | EBMUD | IP | crassphage | 3962 | 6328 | 4730 | 1598 | 2043 | 5 |
| D9 | EBMUD_NT_1_072623 | 1 | EBMUD | NT | crassphage | 2361.8 | 8282 | 4247 | 4035 | 2029 | 50 |
| E9 | EBMUD_NT_2_072623 | 1 | EBMUD | NT | crassphage | 1750.2 | 8289 | 3408 | 4881 | 2029 | 50 |
| A9 | EBMUD_PMG_1_072623 | 1 | EBMUD | PMG | crassphage | 14.2 | 8125 | 40 | 8085 | 2043 | 5 |
| B9 | EBMUD_PMG_2_072623 | 1 | EBMUD | PMG | crassphage | 1.348 | 8240 | 4 | 8236 | 2043 | 5 |
| C9 | EBMUD_PMG_3_072623 | 1 | EBMUD | PMG | crassphage | 52.67 | 6570 | 117 | 6453 | 2043 | 5 |
| G9 | EBMUD_S_1_072623 | 1 | EBMUD | S | crassphage | 970.3 | 8270 | 2151 | 6119 | 2029 | 50 |
| H9 | EBMUD_S_2_072623 | 1 | EBMUD | S | crassphage | 773.1 | 8226 | 1811 | 6415 | 2029 | 50 |
| A10 | EBMUD_S_3_072623 | 1 | EBMUD | S | crassphage | 642.7 | 8075 | 1504 | 6571 | 2029 | 50 |
| B10 | SMCSD_IP_1_080223 | 1 | SMCSD | IP | crassphage | 2891.9 | 8256 | 4892 | 3364 | 2027 | 5 |
| C10 | SMCSD_IP_2_080223 | 1 | SMCSD | IP | crassphage | 4288.3 | 8261 | 6057 | 2204 | 2027 | 5 |
| D10 | SMCSD_IP_3_080223 | 1 | SMCSD | IP | crassphage | 3438.8 | 8270 | 5400 | 2870 | 2027 | 5 |
| D9 | SMCSD_NT_1_080223 | 1 | SMCSD | NT | crassphage | 982 | 8265 | 2136 | 6129 | 2027 | 50 |
| E9 | SMCSD_NT_2_080223 | 1 | SMCSD | NT | crassphage | 866.2 | 8180 | 1886 | 6294 | 2027 | 50 |

| | | | | | | | | | | | |
|---|---|---|---|---|---|---|---|---|---|---|---|
| F9 | SMCSD_NT_3_080223 | 1 | SMCSD | NT | crassphage | 893.4 | 8261 | 1962 | 6299 | 2027 | 50 |
| A9 | SMCSD_PMG_1_080223 | 1 | SMCSD | PMG | crassphage | 11756 | 8257 | 8069 | 188 | 2027 | 5 |
| B9 | SMCSD_PMG_2_080223 | 1 | SMCSD | PMG | crassphage | 14467.1 | 8184 | 8094 | 90 | 2027 | 5 |
| C9 | SMCSD_PMG_3_080223 | 1 | SMCSD | PMG | crassphage | 8473.5 | 8280 | 7683 | 597 | 2027 | 5 |
| G9 | SMCSD_S_1_080223 | 1 | SMCSD | S | crassphage | 481.8 | 8277 | 1150 | 7127 | 2027 | 50 |
| H9 | SMCSD_S_2_080223 | 1 | SMCSD | S | crassphage | 289.5 | 8267 | 735 | 7532 | 2027 | 50 |
| A10 | SMCSD_S_3_080223 | 1 | SMCSD | S | crassphage | 267.8 | 8240 | 678 | 7562 | 2027 | 50 |
| B10 | WCSD_IP_1_073123 | 1 | WCSD | IP | crassphage | 764.9 | 8260 | 1746 | 6514 | 2025 | 5 |
| C10 | WCSD_IP_2_073123 | 1 | WCSD | IP | crassphage | 990.2 | 8249 | 2169 | 6080 | 2025 | 5 |
| D10 | WCSD_IP_3_073123 | 1 | WCSD | IP | crassphage | 729.2 | 8253 | 1659 | 6594 | 2025 | 5 |
| D9 | WCSD_NT_1_073123 | 1 | WCSD | NT | crassphage | 4647.4 | 8274 | 6264 | 2010 | 2025 | 50 |
| E9 | WCSD_NT_2_073123 | 1 | WCSD | NT | crassphage | 4595.1 | 8258 | 6202 | 2056 | 2025 | 50 |
| F9 | WCSD_NT_3_073123 | 1 | WCSD | NT | crassphage | 4094.1 | 8281 | 5891 | 2390 | 2025 | 50 |
| A9 | WCSD_PMG_1_073123 | 1 | WCSD | PMG | crassphage | 7509.6 | 8245 | 7509 | 736 | 2025 | 5 |
| B9 | WCSD_PMG_2_073123 | 1 | WCSD | PMG | crassphage | 3664.7 | 8178 | 5569 | 2609 | 2025 | 5 |
| C9 | WCSD_PMG_3_073123 | 1 | WCSD | PMG | crassphage | 3554.1 | 8277 | 5530 | 2747 | 2025 | 5 |
| G9 | WCSD_S_1_073123 | 1 | WCSD | S | crassphage | 1462.6 | 8209 | 2996 | 5213 | 2025 | 50 |
| H9 | WCSD_S_2_073123 | 1 | WCSD | S | crassphage | 10.54 | 8274 | 28 | 8246 | 2025 | 50 |
| A10 | WCSD_S_3_073123 | 1 | WCSD | S | crassphage | 1108.3 | 8047 | 2407 | 5640 | 2025 | 50 |
| B12 | EBMUD_IP_1_072623 | 1 | EBMUD | IP | pmmov | 528.9 | 8142 | 1272 | 6870 | 2029 | 1 |
| C12 | EBMUD_IP_2_072623 | 1 | EBMUD | IP | pmmov | 327.1 | 8284 | 822 | 7462 | 2029 | 1 |
| D12 | EBMUD_IP_3_072623 | 1 | EBMUD | IP | pmmov | 839.1 | 6736 | 1779 | 4957 | 2044 | 1 |
| D11 | EBMUD_NT_1_083023 | 1 | EBMUD | NT | pmmov | 9941.3 | 8265 | 7888 | 377 | 2034 | 1 |
| E11 | EBMUD_NT_2_083023 | 1 | EBMUD | NT | pmmov | 1121 | 8205 | 2391 | 5814 | 2034 | 1 |
| A11 | EBMUD_PMG_1_072623 | 1 | EBMUD | PMG | pmmov | 17221.6 | 8060 | 8030 | 30 | 2029 | 1 |
| C11 | EBMUD_PMG_3_072623 | 1 | EBMUD | PMG | pmmov | 10779.7 | 6180 | 5963 | 217 | 2029 | 1 |
| B11 | EBMUD_PMG_2_072623 | 1 | EBMUD | PMG | pmmov | 4884.1 | 7878 | 6447 | 1431 | 2044 | 1 |
| H11 | EBMUD_S_2_072623 | 1 | EBMUD | S | pmmov | 416.4 | 8238 | 1016 | 7222 | 2029 | 1 |
| A12 | EBMUD_S_3_072623 | 1 | EBMUD | S | pmmov | 347.1 | 7645 | 816 | 6829 | 2029 | 1 |
| G11 | EBMUD_S_1_072623 | 1 | EBMUD | S | pmmov | 15.05 | 7863 | 41 | 7822 | 2044 | 19 |
| B12 | SMCSD_IP_1_080223 | 1 | SMCSD | IP | pmmov | 2265.8 | 8252 | 4266 | 3986 | 2027 | 1 |
| C12 | SMCSD_IP_2_080223 | 1 | SMCSD | IP | pmmov | 1247.1 | 8179 | 2688 | 5491 | 2027 | 1 |
| D12 | SMCSD_IP_3_080223 | 1 | SMCSD | IP | pmmov | 740.1 | 8273 | 1737 | 6536 | 2027 | 1 |
| D11 | SMCSD_NT_1_080223 | 1 | SMCSD | NT | pmmov | 9862.1 | 8278 | 7891 | 387 | 2027 | 1 |
| E11 | SMCSD_NT_2_080223 | 1 | SMCSD | NT | pmmov | 10120.7 | 8273 | 7904 | 369 | 2027 | 1 |
| F11 | SMCSD_NT_3_080223 | 1 | SMCSD | NT | pmmov | 14106.8 | 8270 | 8160 | 110 | 2027 | 1 |
| A11 | SMCSD_PMG_1_080223 | 1 | SMCSD | PMG | pmmov | 6916.2 | 8262 | 7388 | 874 | 2027 | 1 |
| B11 | SMCSD_PMG_2_080223 | 1 | SMCSD | PMG | pmmov | 11345.8 | 8287 | 8050 | 237 | 2027 | 1 |
| C11 | SMCSD_PMG_3_080223 | 1 | SMCSD | PMG | pmmov | 9035.8 | 8250 | 7752 | 498 | 2027 | 1 |
| G11 | SMCSD_S_1_080223 | 1 | SMCSD | S | pmmov | 329.4 | 8256 | 800 | 7456 | 2027 | 1 |
| H11 | SMCSD_S_2_080223 | 1 | SMCSD | S | pmmov | 204 | 8246 | 515 | 7731 | 2027 | 1 |
| A12 | SMCSD_S_3_080223 | 1 | SMCSD | S | pmmov | 92.69 | 8151 | 242 | 7909 | 2027 | 1 |
| C12 | WCSD_IP_2_073123 | 1 | WCSD | IP | pmmov | 340.2 | 8282 | 853 | 7429 | 2025 | 1 |
| D12 | WCSD_IP_3_073123 | 1 | WCSD | IP | pmmov | 267.9 | 8157 | 667 | 7490 | 2025 | 1 |
| B12 | WCSD_IP_1_073123 | 1 | WCSD | IP | pmmov | 29.84 | 7415 | 75 | 7340 | 2044 | 19 |

| | | | | | | | | | | | |
|---|---|---|---|---|---|---|---|---|---|---|---|
| D11 | WCSD_NT_1_073123 | 1 | WCSD | NT | pmmov | | | | | 2044 | |
| E11 | WCSD_NT_2_073123 | 1 | WCSD | NT | pmmov | | | | | 2044 | |
| F11 | WCSD_NT_3_073123 | 1 | WCSD | NT | pmmov | | | | | 2044 | |
| A11 | WCSD_PMG_1_073123 | 1 | WCSD | PMG | pmmov | 1880.3 | 7495 | 3598 | 3897 | 2044 | 19 |
| B11 | WCSD_PMG_2_073123 | 1 | WCSD | PMG | pmmov | 1595.2 | 8132 | 3354 | 4778 | 2044 | 19 |
| C11 | WCSD_PMG_3_073123 | 1 | WCSD | PMG | pmmov | 1453.3 | 7821 | 2941 | 4880 | 2044 | 19 |
| G11 | WCSD_S_1_073123 | 1 | WCSD | S | pmmov | 2072.7 | 6669 | 3157 | 3512 | 2025 | 1 |
| H11 | WCSD_S_2_073123 | 1 | WCSD | S | pmmov | 978 | 4332 | 1152 | 3180 | 2025 | 1 |
| A12 | WCSD_S_3_073123 | 1 | WCSD | S | pmmov | 1121 | 7975 | 2436 | 5539 | 2025 | 1 |
| B10 | EBMUD_IP_1_083023 | 2 | EBMUD | IP | crassphage | 3562.8 | 8223 | 4612 | 1492 | 2034 | 1 |
| C10 | EBMUD_IP_2_083023 | 2 | EBMUD | IP | crassphage | 4297 | 8253 | 6057 | 2196 | 2034 | 1 |
| D10 | EBMUD_IP_3_083023 | 2 | EBMUD | IP | crassphage | 2658.8 | 8266 | 4619 | 3647 | 2034 | 1 |
| F9 | EBMUD_NT_3_083023 | 2 | EBMUD | NT | crassphage | 4145.2 | 8201 | 6203 | 1998 | 2043 | 50 |
| D9 | EBMUD_NT_1_083023 | 2 | EBMUD | NT | crassphage | 2687.3 | 7681 | 4607 | 3074 | 2043 | 50 |
| E9 | EBMUD_NT_2_083023 | 2 | EBMUD | NT | crassphage | 2421.3 | 7823 | 4432 | 3391 | 2043 | 50 |
| A9 | EBMUD_PMG_1_083023 | 2 | EBMUD | PMG | crassphage | 967.4 | 7971 | 2136 | 5835 | 2043 | 50 |
| B9 | EBMUD_PMG_2_083023 | 2 | EBMUD | PMG | crassphage | 1752.7 | 8249 | 3501 | 4748 | 2043 | 50 |
| C9 | EBMUD_PMG_3_083023 | 2 | EBMUD | PMG | crassphage | 846.1 | 7787 | 1841 | 5946 | 2043 | 50 |
| G9 | EBMUD_S_1_083023 | 2 | EBMUD | S | crassphage | 14614.5 | 3924 | 3882 | 42 | 2034 | 1 |
| H9 | EBMUD_S_2_083023 | 2 | EBMUD | S | crassphage | 23116.3 | 6775 | 6771 | 4 | 2034 | 1 |
| A10 | EBMUD_S_3_083023 | 2 | EBMUD | S | crassphage | 16530 | 8221 | 8180 | 41 | 2034 | 1 |
| B10 | SMCSD_IP_1_091923 | 2 | SMCSD | IP | crassphage | 2518.9 | 8256 | 4479 | 3777 | 2038 | 5 |
| C10 | SMCSD_IP_2_091923 | 2 | SMCSD | IP | crassphage | 2345.6 | 8279 | 4260 | 4019 | 2038 | 5 |
| D10 | SMCSD_IP_3_091923 | 2 | SMCSD | IP | crassphage | 2869.3 | 8280 | 4856 | 3424 | 2038 | 5 |
| D9 | SMCSD_NT_1_091923 | 2 | SMCSD | NT | crassphage | 1567.7 | 8194 | 3110 | 5084 | 2038 | 50 |
| E9 | SMCSD_NT_2_091923 | 2 | SMCSD | NT | crassphage | 2231.4 | 8272 | 4061 | 4211 | 2038 | 50 |
| F9 | SMCSD_NT_3_091923 | 2 | SMCSD | NT | crassphage | 1252.2 | 8144 | 2575 | 5569 | 2038 | 50 |
| A9 | SMCSD_PMG_1_091923 | 2 | SMCSD | PMG | crassphage | 4311.5 | 6611 | 4931 | 1680 | 2043 | 5 |
| B9 | SMCSD_PMG_2_091923 | 2 | SMCSD | PMG | crassphage | 4952.7 | 7390 | 5868 | 1522 | 2043 | 5 |
| C9 | SMCSD_PMG_3_091923 | 2 | SMCSD | PMG | crassphage | 3008.7 | 7610 | 4760 | 2850 | 2043 | 5 |
| G9 | SMCSD_S_1_091923 | 2 | SMCSD | S | crassphage | 157.4 | 8259 | 394 | 7865 | 2038 | 50 |
| H9 | SMCSD_S_2_091923 | 2 | SMCSD | S | crassphage | 111.7 | 8217 | 290 | 7927 | 2038 | 50 |
| A10 | SMCSD_S_3_091923 | 2 | SMCSD | S | crassphage | 1736.2 | 7699 | 3420 | 4279 | 2043 | 50 |
| B10 | WCSD_IP_1_090523 | 2 | WCSD | IP | crassphage | 2305.5 | 8273 | 4229 | 4044 | 2035 | 5 |
| C10 | WCSD_IP_2_090523 | 2 | WCSD | IP | crassphage | 2641 | 8258 | 4598 | 3660 | 2035 | 5 |
| D10 | WCSD_IP_3_090523 | 2 | WCSD | IP | crassphage | 1340.8 | 8104 | 2740 | 5364 | 2035 | 5 |
| D9 | WCSD_NT_1_090523 | 2 | WCSD | NT | crassphage | 618.1 | 8249 | 1415 | 6834 | 2035 | 50 |
| E9 | WCSD_NT_2_090523 | 2 | WCSD | NT | crassphage | 1379.7 | 8163 | 2786 | 5377 | 2035 | 50 |
| F9 | WCSD_NT_3_090523 | 2 | WCSD | NT | crassphage | 838 | 8224 | 1847 | 6377 | 2035 | 50 |
| A9 | WCSD_PMG_1_090523 | 2 | WCSD | PMG | crassphage | 1833.1 | 8264 | 3682 | 4582 | 2035 | 5 |
| B9 | WCSD_PMG_2_090523 | 2 | WCSD | PMG | crassphage | 1778.9 | 8222 | 3500 | 4722 | 2035 | 5 |
| C9 | WCSD_PMG_3_090523 | 2 | WCSD | PMG | crassphage | 5127.2 | 8253 | 6572 | 1681 | 2035 | 5 |
| G9 | WCSD_S_1_090523 | 2 | WCSD | S | crassphage | 74.21 | 8254 | 188 | 8066 | 2035 | 50 |
| H9 | WCSD_S_2_090523 | 2 | WCSD | S | crassphage | 13.22 | 8252 | 35 | 8217 | 2035 | 50 |
| A10 | WCSD_S_3_090523 | 2 | WCSD | S | crassphage | 1202.7 | 8181 | 2618 | 5563 | 2035 | 50 |

| | | | | | | | | | | | |
|---|---|---|---|---|---|---|---|---|---|---|---|
| B12 | EBMUD_IP_1_083023 | 2 | EBMUD | IP | pmmov | 382.2 | 8277 | 956 | 7321 | 2034 | 1 |
| C12 | EBMUD_IP_2_083023 | 2 | EBMUD | IP | pmmov | 343.8 | 8276 | 861 | 7415 | 2034 | 1 |
| D12 | EBMUD_IP_3_083023 | 2 | EBMUD | IP | pmmov | 745.2 | 8218 | 1736 | 6482 | 2034 | 1 |
| D11 | EBMUD_NT_1_072623 | 2 | EBMUD | NT | pmmov | 8496 | 7165 | 6653 | 512 | 2029 | 1 |
| E11 | EBMUD_NT_2_072623 | 2 | EBMUD | NT | pmmov | 8252.7 | 5670 | 5221 | 449 | 2029 | 1 |
| F11 | EBMUD_NT_3_083023 | 2 | EBMUD | NT | pmmov | 1376.3 | 8185 | 2815 | 5370 | 2034 | 1 |
| A11 | EBMUD_PMG_1_083023 | 2 | EBMUD | PMG | pmmov | 404.1 | 7837 | 948 | 6889 | 2044 | 50 |
| B11 | EBMUD_PMG_2_083023 | 2 | EBMUD | PMG | pmmov | 514.8 | 7984 | 1235 | 6749 | 2044 | 50 |
| C11 | EBMUD_PMG_3_083023 | 2 | EBMUD | PMG | pmmov | 341.3 | 7988 | 871 | 7117 | 2044 | 50 |
| G11 | EBMUD_S_1_083023 | 2 | EBMUD | S | pmmov | 460.7 | 8212 | 1091 | 7121 | 2034 | 1 |
| H11 | EBMUD_S_2_083023 | 2 | EBMUD | S | pmmov | 559.3 | 8059 | 1306 | 6753 | 2034 | 1 |
| A12 | EBMUD_S_3_083023 | 2 | EBMUD | S | pmmov | 493.1 | 8256 | 1223 | 7033 | 2034 | 1 |
| B12 | SMCSD_IP_1_091923 | 2 | SMCSD | IP | pmmov | 242.5 | 8262 | 619 | 7643 | 2038 | 1 |
| C12 | SMCSD_IP_2_091923 | 2 | SMCSD | IP | pmmov | 147.9 | 8214 | 379 | 7835 | 2038 | 1 |
| D12 | SMCSD_IP_3_091923 | 2 | SMCSD | IP | pmmov | 210.7 | 8214 | 533 | 7681 | 2038 | 1 |
| D11 | SMCSD_NT_1_091923 | 2 | SMCSD | NT | pmmov | 139 | 8282 | 350 | 7932 | 2038 | 1 |
| E11 | SMCSD_NT_2_091923 | 2 | SMCSD | NT | pmmov | 148.2 | 8244 | 367 | 7877 | 2038 | 1 |
| F11 | SMCSD_NT_3_091923 | 2 | SMCSD | NT | pmmov | 148.9 | 8233 | 367 | 7866 | 2038 | 1 |
| A11 | SMCSD_PMG_1_091923 | 2 | SMCSD | PMG | pmmov | 995.8 | 8164 | 2256 | 5908 | 2038 | 1 |
| B11 | SMCSD_PMG_2_091923 | 2 | SMCSD | PMG | pmmov | 2218.4 | 8263 | 4139 | 4124 | 2038 | 1 |
| C11 | SMCSD_PMG_3_091923 | 2 | SMCSD | PMG | pmmov | 1448.9 | 8213 | 2977 | 5236 | 2038 | 1 |
| G11 | SMCSD_S_1_091923 | 2 | SMCSD | S | pmmov | 9.801 | 8257 | 25 | 8232 | 2038 | 1 |
| H11 | SMCSD_S_2_091923 | 2 | SMCSD | S | pmmov | 6.932 | 8224 | 18 | 8206 | 2038 | 1 |
| A12 | SMCSD_S_3_091923 | 2 | SMCSD | S | pmmov | 6.74 | 8222 | 18 | 8204 | 2038 | 1 |
| B12 | WCSD_IP_1_090523 | 2 | WCSD | IP | pmmov | 1094.8 | 8221 | 2437 | 5784 | 2035 | 1 |
| C12 | WCSD_IP_2_090523 | 2 | WCSD | IP | pmmov | 1229.4 | 8207 | 2666 | 5541 | 2035 | 1 |
| D12 | WCSD_IP_3_090523 | 2 | WCSD | IP | pmmov | 1026.5 | 8227 | 2294 | 5933 | 2035 | 1 |
| D11 | WCSD_NT_1_090523 | 2 | WCSD | NT | pmmov | 858 | 8046 | 2139 | 5907 | 2044 | 19 |
| E11 | WCSD_NT_2_090523 | 2 | WCSD | NT | pmmov | 902 | 8080 | 2140 | 5940 | 2044 | 19 |
| F11 | WCSD_NT_3_090523 | 2 | WCSD | NT | pmmov | 809.8 | 7924 | 1821 | 6103 | 2044 | 19 |
| A11 | WCSD_PMG_1_090523 | 2 | WCSD | PMG | pmmov | 1013.6 | 7994 | 2215 | 5779 | 2043 | 20 |
| B11 | WCSD_PMG_2_090523 | 2 | WCSD | PMG | pmmov | 1446.1 | 8096 | 3024 | 5072 | 2043 | 20 |
| C11 | WCSD_PMG_3_090523 | 2 | WCSD | PMG | pmmov | 529.6 | 8223 | 1375 | 6848 | 2043 | 20 |
| G11 | WCSD_S_1_090523 | 2 | WCSD | S | pmmov | 1207.6 | 8195 | 2555 | 5640 | 2035 | 1 |
| H11 | WCSD_S_2_090523 | 2 | WCSD | S | pmmov | 1250.4 | 8141 | 2658 | 5483 | 2035 | 1 |
| A12 | WCSD_S_3_090523 | 2 | WCSD | S | pmmov | 1692.3 | 8268 | 3499 | 4769 | 2035 | 1 |
